# Triangulation fails when neither linguistic, genetic, nor archaeological data support the Transeurasian narrative

**DOI:** 10.1101/2022.06.09.495471

**Authors:** Zheng Tian, Yuxin Tao, Kongyang Zhu, Guillaume Jacques, Robin J. Ryder, José Andrés Alonso de la Fuente, Anton Antonov, Ziyang Xia, Yuxuan Zhang, Xiaoyan Ji, Xiaoying Ren, Guanglin He, Jianxin Guo, Rui Wang, Xiaomin Yang, Jing Zhao, Dan Xu, Russell D. Gray, Menghan Zhang, Shaoqing Wen, Chuan-Chao Wang, Thomas Pellard

## Abstract

Robbeets et al.^1^ argue that the dispersal of the so-called “Transeurasian” languages, a highly disputed language superfamily comprising the Turkic, Mongolian, Tungusic, Koreanic, and Japonic language families, was driven by Neolithic farmers in the West Liao River region of China. They adduce evidence from linguistics, archaeology, and genetics to support their claim. An admirable feature of the Robbeets et al.’s paper is that all their datasets can be accessed. However, a closer investigation of all three types of evidence reveals fundamental problems with each of them. Robbeets et al.’s analysis of the linguistic data does not conform to the minimal standards required by traditional scholarship in historical linguistics and contradicts their own stated sound correspondence principles. A reanalysis of the genetic data finds that they do not conclusively support the farming-driven dispersal of Turkic, Mongolian, and Tungusic, nor the two-wave spread of farming to Korea. Their archaeological data contain little phylogenetic signal, and we failed to reproduce the results supporting their core hypotheses about migrations.

Given the severe problems we identify in all three parts of the “triangulation” process, we conclude that there is neither conclusive evidence for a Transeurasian language family nor for associating the five different language families with the spread of Neolithic farmers from the West Liao River region.

## Linguistics

The hypothesis of common ancestry between the five language families included in Robbeets et al.’s Transeurasian hypothesis has a long history, and it has always been a highly controversial question. The authors’ specific version of this hypothesis rests on a list of putative cognate words shared between these branches. *Cognate words* have evolved from a common ancestor, similar to homologous genes in biology. In order to prove that words are cognate, scholars need to present *regular sound correspondences*, which are considered the only valid proof of linguistic genealogical relationships^2–4^. In order to identify regular sound correspondences, both borrowed words and chance resemblances must be excluded.

Out of the 3166 cognate sets listed by Robbeets et al. in support of the Transeurasian hypothesis, only 317 are found in more than one language family, and only 50 of these are shared by more than two families and thus could be taken as evidence for the Transeurasian hypothesis. Only two cognate sets are shared by all five families. Among the 50 cognate sets, five contain words marked as borrowings by the authors but strangely have not been excluded from their analysis. Two contain forms wrongly attributed to a different language, most likely due to copy-paste mistakes. A systematic computer-assisted analysis^5^ of the remaining cognate sets reveals that only 17 etymologies follow the criteria for the identification of regular sound correspondences outlined by the authors (Figure 1a, Supplementary Information 1). This means that only 17 out of 3166 cognate sets might support the hypothesis that the five language families have sprung from a common source, even though most suffer from other problems (Supplementary Information 1). The core evidence presented is thus inadequate, and the Transeurasian hypothesis remains unwarranted.

**Figure 1:**
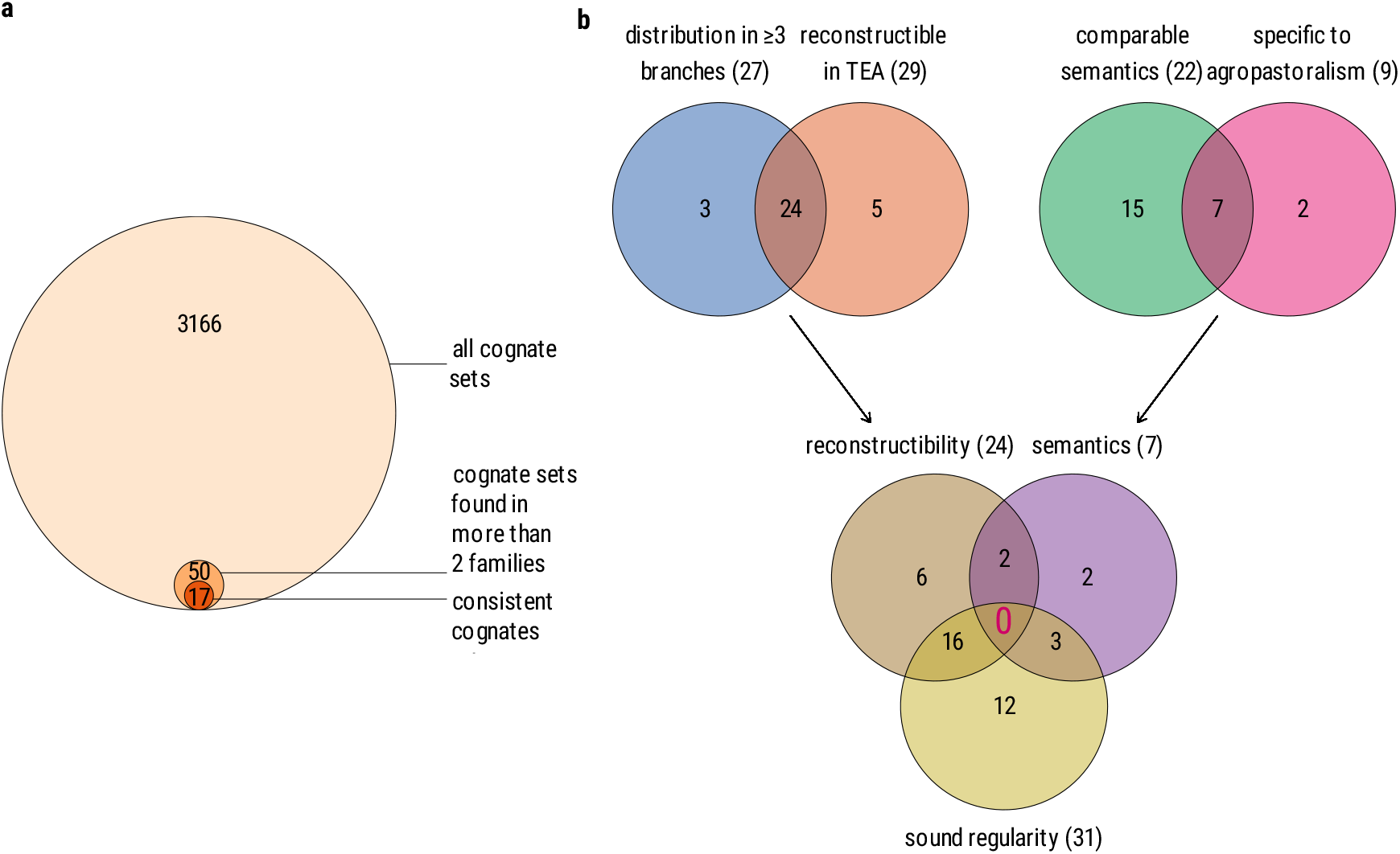
Number of lexical comparisons in Robbeets et al. (2021) that satisfy the different necessary criteria to support **a** the existence of the Transeurasian language family, **b** the hypothesis that the Proto-Transeurasian speakers were early farmers.

Similar problems can be found in the 43 agropastoral comparisons proposed in support of the hypothesis that the Proto-Transeurasian speakers were early farmers. Although 31 comparisons follow the proposed sound correspondences, only 29 could be reconstructed at the Proto-Transeurasian level, only 27 appear in more than three branches, only 22 have comparable semantics, and only 9 items belong to the realm of agropastoral vocabulary. With all these problems, none of the 43 items listed fulfills the criteria laid out by the authors themselves (Figure 1b; Supplementary Information 2). The identification of Proto-Transeurasian speakers with early millet farmers of the West Liao River area is thus not supported by empirical linguistic evidence.

A pervasive issue with the Transeurasian hypothesis, already noted by several specialists^6,7^, is the general opacity of its methodology, which also often ignores the known history of the languages involved. Our detailed investigation of the linguistic data the authors used for their triangulation confirms earlier criticisms and shows that the authors ignore their own principles of historical language comparison and bend the evidence to their needs (Supplementary Information 1 and 2).

We, therefore, conclude that the linguistic evidence is insufficient to distinguish between chance resemblance, contact and inheritance and therefore does not support Robbeets et al.’s claims that the five Transeurasian families share common ancestry and that their dispersal can be correlated with the spread of Early Neolithic millet farmers across Northeast Asia.

## Genetics

Robbeets et al. claim that the early spread of Transeurasian speakers was driven by agriculture based on their finding of West Liao River farmer ancestry with Yellow River farmer admixture in ancient samples collected from Korea and Japan. This assumption is unjustified. There was a long-term genetic continuity of Amur-related hunter-gatherer ancestry without West Liao River and Yellow River farmer admixture in the vast region covering the Mongolian Plateau, Lake Baikal, Amur River Basin and Russian Far East from the Late Palaeolithic to the Iron Age (14 kya to 2 kya)^8–11^. The widespread pattern of this Amur ancestry was already formed at least 8 kya, but agriculture had just started to appear in northeast Asia at that time^11^. The explanation of Robbeets et al. regarding this early widespread Amur ancestry crucially depends on the conjecture that the early Neolithic farmers in the West Liao River, such as the Xinglongwa (8200–7400 BP) people were genetically Amur-like without influence from Yellow River. This conjecture is contradicted by their claim that both the spread of Japonic and that of Koreanic were induced by West Liao River farmer ancestry with Yellow River farmer admixture. However, the genetic modelling of ancient Korean and Japanese formation in Figure 3 of Robbeets et al. is problematic. They associated the spread of farming to Korea with different waves of Amur and Yellow River gene flow, modelled by Hongshan for the Neolithic introduction of millet farming and Upper Xiajiadian for the Bronze Age addition of rice agriculture. This is, however, contradicted by the authors’ own statement in the supplementary material that their data lack the resolution to distinguish between competing admixture models (SI 13 in Robbeets et al.).

To test the robustness of this inference, we reanalyzed their data using *f* _4_ statistics and *qpWave* and found that Hongshan and Upper Xiajiadian were genetically equally related to the ancient Korean and Japanese (Figure 2a and b). The populations they selectively assigned with Upper Xiajiadian ancestry in their Figure 3 could also be explained by using Hongshan instead of Upper Xiajiadian as a genetic source (Figure 2c).

**Figure 2:**
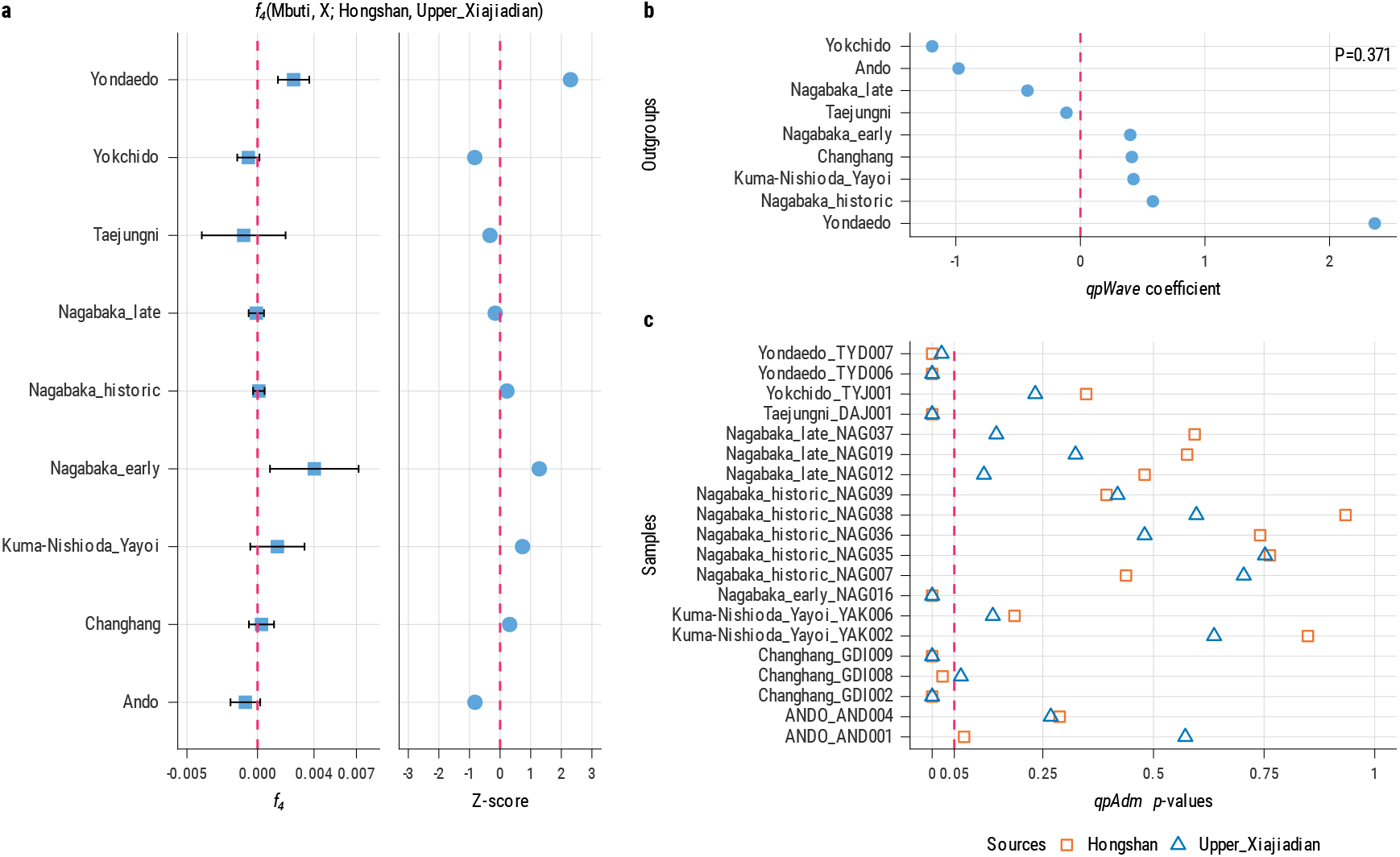
**a**, the *f* _4_ statistics in the form *f* _4_ (Mbuti, X; Hongshan, Upper_Xiajidian), in which there are no significant Z-scores. **b**, the homogeneity test using *qpWave* shows that Hongshan and Upper Xiajiadian are homogenous to all the newly generated samples with *P* = 0.371. **c**, the *p* values of *qpAdm* modelling of two-way admixture of ancient populations in Korea and Japan, in which we were not able to distinguish the Hongshan and Xiajiadian ancestry in the formation of targeted populations.

**Figure 3:**
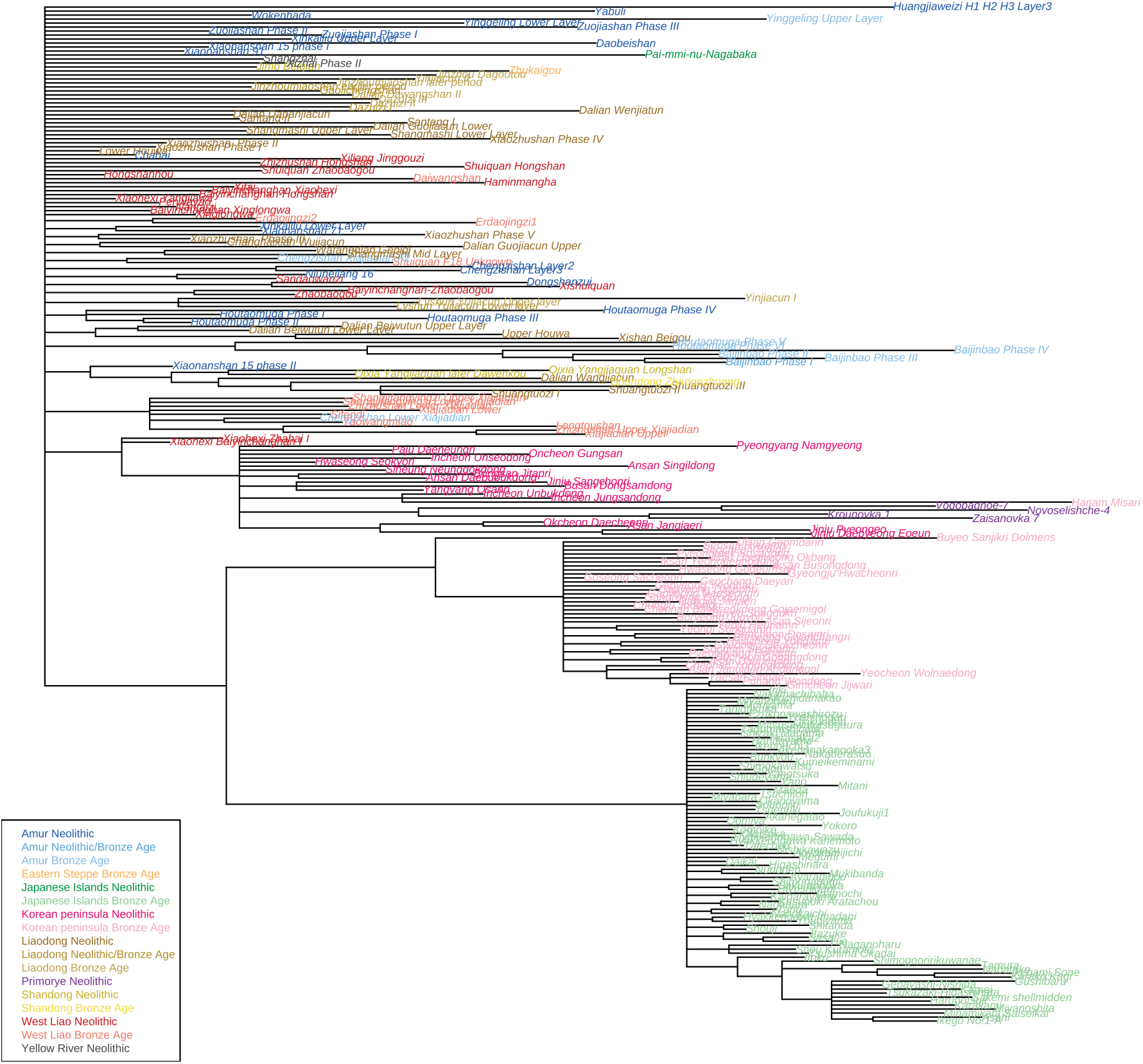
Majority rule consensus tree including only those clades that are present in the majority (≥ 50%) of the trees in our analysis of the corrected archaeological data. Colours indicate the region and period of the different archaeological sites.

Thus, from a genetic perspective, we conclude that Robbeets et al’s claims rest on an unjustified assumption and a selective modelling of migration hypotheses that excludes viable alternative hypotheses.

## Archaeology

Robbeets et al. (2021) collected 171 archaeological features for 255 Neolithic and Bronze Age sites in northern China, the Primorye, Korea and Japan and produced a phylogenetic analysis of these data using BEAST. Their analysis assumes that a phylogenetic tree is a good representation of the history of the archaeological sites. Phylogenies are commonly used for genes and languages, and they have also been used successfully for cultural data. However, a justification that a tree structure is appropriate for the data remains essential. For these data, no evidence is given that the traits considered have evolved along a tree (with vertical transmission from ancestor to descendant, rather than horizontal transmission between neighbours), nor that a single phylogeny can adequately capture the history of the very diverse traits considered. We tried to reproduce their results and conducted several additional analyses (Supplementary Information 3). We found very little evidence of tree-like signal in the data and that different subsets of the data have different histories.

Robbeets et al. (2021) use the results of their Bayesian analysis to support two major hypotheses. First, they argue for a Neolithic forking dispersal of millet agriculture from the West Liao river basin to Korea by 5500 BP (or 6500 BP, two distinct dates are given) and Primorye by 5000 BP. The first migration from West Liao to Korea is supported in their results by a clade comprising two West Liao river sites, Xiaohexi Chahai I (site #8 in SI 6 of Robbeets et al.) and Xiaohexi Bayinchanhan (#7) and the Chulmun culture sites and is interpreted as corresponding to the spread of the Koreanic language family. The second migration to Primorye (Zaisanovka-Yabuli culture) is supported by a clade formed by the Primorye Zaisanovka-Yabuli culture sites with Liaodong and Amur sites and is associated with the Tungusic family.

In contrast, in our results (Figure 3), we find a clade comprising the West Liao river sites Xiaohexi Chahai I (#8) and Xiaohexi Bayinchanhan (#7) together with Chulmun (Korean peninsula Neolithic), Zaisanovka and one Mumun site (#117). We thus find no support for the first hypothesis of two migrations corresponding to distinct putative branches of Transeurasian.

Second, Robbeets et al. (2021) argue for a Bronze Age migration from the West Liao river and Shandong to Korea and Japan (Mumun and Yayoi cultures) by 3500 BP that would correspond to the spread of the Japonic family. This is supported in their results by a clade comprising Mumun and Yayoi sites as well as Bronze Age West Liao sites such as Xiajiadian. However, in our results (Figure 3), though there is support for a clade comprising both the Yayoi culture (Japanese Islands Bronze Age) and Mumun (Korean Islands Bronze Age) sites, we find no evidence in favour of a close relationship with Bronze age West Liao river sites.

We also performed analyses of different subsets of the data, following Robbeets et al.’s feature classification into ceramics, stone tools, food remains, buildings/houses, shell and bone artifacts, and burials (Supplementary Information 3). Our analysis restricted to ceramic features yields a highly unresolved, rake-shaped tree that does not support any of the hypotheses laid out by Robbeets et al. Other analyses of subsets of the data show even less phylogenetic signal. The analysis restricted to buildings/houses, shell and bone artifacts and burials did not recover any clade corresponding to known archaeological cultures. Our results show that the archaeological features selected by Robbeets et al. are highly heterogeneous and evolved through different historical processes. Simply combining all of them is unlikely to yield results reflecting their complex history.

We, therefore, conclude that the archaeological evidence is weak, contains conflicting signals, and does not support the central claims of Robbeets et al.

## Conclusions

Our attempts to replicate Robbeets et al.’s results show significant discrepancies on all three fronts. Moreover, Robbeet’s et al. claim to use “triangulation” to aggregate the results of the three studies, but they neither define nor describe the method of “triangulation”. It is thus impossible to follow their method and combine the three corrected analyses. Taken together, our re-evaluation of all three datasets completely undermines Robbeets et al.’s narrative. The Transeurasian hypothesis remains unsubstantiated. We respectfully suggest the paper be withdrawn.

## Acknowledgements

The work was funded by the Department of Linguistic and Cultural Evolution, Max Planck Institute for Evolutionary Anthropology, Leipzig, Germany, the National Natural Science Foundation of China (32070576, 32070577, 91731303, 31771325, 31801040, and T2122007), the National Key R&D Program of China (2020YFC1521607), the Major Project of National Social Science Foundation of China (21&ZD285, 20&ZD248, and 20&ZD212), the “Double First-Class University Plan” key construction project of Xiamen University (the origin and evolution of East Asian populations and the spread of Chinese civilization, 0310/X2106027), Nanqiang Outstanding Young Talents Program of Xiamen University (X2123302), the Major Project of National Social Science Foundation of China (2021MZD014), the European Research Council (ERC) grant to Dan Xu (ERC-2019-ADG-883700-TRAM), the “Shuguang Program” supported by Shanghai Education Development Foundation and Shanghai Municipal Education Commission (20SG06), the Scientific and Technology Committee of Shanghai Municipality (18490750300), the National Key Research and Development Program (2020YFE0201600), Shanghai Municipal Science and Technology Major Project (2017SHZDZX01), the 111 Project (B13016), and the China Postdoctoral Science Foundation (2021M691879, 2021M691882). We thank Johann-Mattis List for providing help with the code for checking the sound regularity of the deep cognate sets and the distribution of cognate sets across different language families. We thank Johannes Krause and Simon Greenhill for comments on the draft manuscript.

## Author Contributions

CCW, MHZ, RDG, SQW, GJ and TP initiated the study. CCW, MHZ, YXT, and SQW wrote the first draft. TP, GJ, RJR and RDG extended the first draft. TP, GJ, JAAF, AA, RJR, YXT, and ZX checked, compiled, and analyzed linguistic data. MHZ and YXT contributed code for linguistic analyses. TP, ZT, KZ and RJR contributed code for the figures. TP, GJ, JAAF and AA wrote the etymological comments in the Supplementary Information 1 and 2. ZT, YZ, XJ, XR, and SQW compiled the archaeological data. ZT, YZ, SQW, GJ, RJR and TP analyzed the archaeological data and wrote the Supplementary Information 3. KZ, ZX, GH, JG, RW, XY, JZ, and CCW compiled and analyzed the genetic data. KZ and CCW wrote the Supplementary Information 2. DX was involved in the discussion. All authors have read and agree with the final version of this study.

