## Supplementary Information for "Triangulation fails when neither linguistic, genetic, nor archaeological data support the Transeurasian narrative"

June 9, 2022

The data and computer code used in this project are available online at `https://osf.io/jdvp3/?view_only=3bbd0c50f59247ea9fed160beedf2bf7`.

**Abbreviations 63**

**References 65**

### 1 Evaluation of the core linguistic evidence for the Transeurasian hypothesis

#### 1.1 Introduction

Since the so-called "Transeurasian" hypothesis of a phylogenetic relationship between the Japonic, Koreanic, Tungusic, Mongolic, and Turkic languages, has always been controversial and still remains a minority view, we set out to try replicating the results of Robbeets et al. (2021) and testing the validity of their evidence and the consistency of their methods. As recognized by Robbeets et al. (2021) themselves, the central issue is to distinguish genuine cognates, that would constitute evidence for a phylogenetic relationship, from borrowings and chance resemblances. To this end, Robbeets et al. (2021) list seven criteria (SI5: 1–2, see also Robbeets 2020). We propose to concentrate on the first two criteria, which are the only ones that are applicable to all comparisons and which can be tested automatically. These criteria can be formulated as follows:

1. a cognate set constitutes core evidence for the Transeurasian hypothesis if it is attested in three or more language families;
2. a cognate set that violates the sound correspondences for the initial (Consonant-)Vowel(-Consonant) sequence is irregular and cannot be used as evidence for the Transeurasian hypothesis.

The remaining five criteria require expert knowledge and extensive qualitative analysis, and we did not use them to reject proposed comparisons since they are not easily replicable.

The correspondence patterns are reported by Robbeets et al. (2021) in the form of several tables, in which columns correspond to the five language families and the hypothetical Transeurasian proto-language, and rows correspond to correspondence patterns, as exemplified in Table 1.1.

| ID | proto-Japonic | proto-Koreanic | proto-Tungusic | proto-Mongolic | proto-Turkic | proto-Transeurasian |
|---|---|---|---|---|---|---|
| 1 | *p- | *p- | *p- | *p- | *b-/*p- | *p- |
| 13 | *k- | *k- | *k- | *k- | *k- | *k- |
| 15 | *k- | *k- | *g- | *g- | *k- | *g- |
| 16 | *-k- | *-k- (-h-) | *-g- | *-g- | *-g- | *-g- |

Table 1.1: Examples of sound correspondences (from SI2 Table 3.11)

This table can be interpreted in two ways. On the one hand, when we start from the Transeurasian entry, we expect that for a given pTEA form, the reflexes in the five families—if attested—should exhibit the sounds listed in the table. This means, that if a pTEA form starts with a *p, we expect to find a *b or a *p in pTk, and a *p in all other proto-languages. Similarly, if a pTEA form contains a medial *g, we expect to find a *g in pTk, pM, and pTg, and a *k in pJ, as well as a *k or a *h in pK.

It is also possible to proceed in a bottom-up manner, and check whether a cognate set involving three or more of the five language families instantiates a valid correspondence pattern. Thus, if there is a cognate set with reflexes in Tungusic and Mongolic in which the pTg and pM forms start with a *k-, we could identify this comparison as "regular" according to the correspondence table provided by the authors (corr. #13). However, if the pM form had a *g- instead, we would classify this comparison as "irregular", since we do not find this pattern in the correspondence table.

#### 1.2 Data

In order to try replicating Robbeets et al.'s (2021) results, we adopted the bottom-up procedure mentioned in the previous paragraph. We first collected all cognates distributed in at least three language families (criterion #1), based on the data by Robbeets et al. (2021) used for the phylogenetic analysis (called SI1 in the following).[1] In order to collect the cognate sets, we first converted SI1 to the spreadsheet formats recommended by the Cross-Linguistic Data Formats Initiative (`https://cldf.clld.org`, Forkel et al. 2018) and implemented as part of the Lexibank wordlist repository (List et al. 2021). The resulting CLDF datasets of Robbeets et al.'s (2021) comparative wordlist is curated on GitHub (`https://github.com/digling/triangulation`, Version 0.1). We then searched for all cognate sets in the data which occur in at least three different language families. This first step yielded 50 cognate sets which constitute the core evidence for the putative Transeurasian hypothesis. Out of these 50 forms, 43 qualify for our analysis, since seven contain mistakes due to obvious coding errors, or borrowings that were marked as such but apparently not excluded by the authors in their phylogenetic analysis.

The proto-forms for the different families were extracted from the supplementary material provided by Robbeets et al. (2021) (called SI2 in the following).[2] When a proto-form was missing, we proposed a tentative reconstruction on the basis of the sound correspondence tables whenever possible. Such forms were marked with a double asterisk (**) in our annotations. The resulting table of 50 cognate sets (seven marked for exclusion) along with our proposed proto-forms and code that illustrates how they could be extracted from the CLDF dataset derived from SI1 is available at `https://osf.io/jdvp3/?view_only=3bbd0c50f59247ea9fed160beedf2bf7`.

#### 1.3 Method

In order to test the validity of the proposed comparisons in terms of the regularity of their sound correspondences, we examined the first three sounds of each word in a cognate set, which correspond to a $C_1VC_2$ template. Words starting with a vowel were labeled as `- V` $C_2$, words consisting of a vowel alone as `- V -`, and words consisting of a consonant and a vowel alone as $C_1$ `V -`. For each slot in a given cognate set, we aligned the sounds found in the different proto-languages to the $C_1VC_2$ template, which in turn yields a multiple alignment for all forms in a cognate set (see Wu et al. 2020 for details on template alignment techniques). These aligned sounds constitute a correspondence pattern (see List 2019 for details on the notion of correspondence patterns), and it was annotated as regular if it matched any pattern (row) in the correspondence tables of SI2. A cognate set was marked as valid if all slots contained a regular correspondence involving three or more proto-languages. The whole process of comparing if sound correspondences recur in the correspondence table was carried out automatically.

[1] File `16_Eurasia3angle_synthesis_SI 1_BV 254_REV2021.09.22.xlsx` in the supplement on Figshare.
[2] File `17_Eurasia3angle_synthesis_SI 2_basic etymologies.pdf` in the supplementary material on FigShare.

Our procedure was permissive in so far as we did not penalize those cases in which the authors provided more than one possible sound in one of the five proto-languages for a given correspondence pattern (as in correspondence #16 above for pK). In such cases, we broke down the different alternatives into distinct correspondence patterns, which increases the probability of a match. Likewise, when several alternative reconstructed forms were listed for a proto-language, we counted the cognate set as regular if regular correspondences obtained with any of the alternative forms.

We then measured the overall regularity of a given cognate set by counting the proportion of slots with regular correspondence patterns. We also automatically reconstructed proto-forms on the basis of the aligned forms and the correspondence patterns listed in the tables of SI2 and compared them with those provided in SI1. For the case of pTg and pM *p and pTk *b, this would yield a *p in pTEA (correspondence #1).

#### 1.4 Results

Table 1.3 lists the selected 50 comparisons together with our automatically computed consistency judgments. Rows in which indices are indicated with an asterisk mark forms which were excluded due to errors in the data (borrowings, copy-paste errors, etc.). The column RECONSTRUCTION gives our automated reconstruction based on the identified sound correspondences. Question marks are used to indicate that no correspondence pattern could be found in the correspondence tables. The column REFLEXES summarizes the number of reflexes that we identified. The column SCORE provides our score based on the number of sound segments following the sound correspondences of SI2, and the column HITS indicates the number of correspondence patterns matching the sound correspondences of SI2.

Out of the 43 cognate sets we analyzed, only 17 exhibit regular sound correspondences as defined above. On average, only 65% of the segments in the 43 cognate sets we analyzed are regular, which must be considered as low, given that we only looked at the first three segments of each reflex only, although proto-forms are at times much longer.

In addition to checking the consistency of cognate sets, our method allows us to reconstruct putative proto-forms in the system by Robbeets et al. (2021) When comparing these proto-forms with the proto-forms for Transeurasian (or putative intermediate stages, which are not distinguished by the authors), we can see many discrepancies as well, which indicate that their reconstruction of the putative Transeurasian proto-language is at its best premature.

The judgments of Robbeets et al. (2021) regarding whether a comparison constitutes valid evidence for the Transeurasian hypothesis fail to replicate most of the time. The low number of cognate sets that follow the criteria of sound regularity and distribution over three or more families seriously undermines the Transeurasian hypothesis.

As a final analysis, we compared the cognate set distributions reported by Robbeets et al. (2021) with the distributions found in well-established language families of similar age, such as Sino-Tibetan or Indo-European (Table 1.4). The number of cognate sets proposed for Transeurasian is exceptionally low, which provides additional evidence that the "Transeurasian" hypothesis does not rest on solid grounds.

| ID | PROTO_TEA | CONCEPT | RECONSTRUCTION | REFLEXES | SCORE | HITS |
|---|---|---|---|---|---|---|
| 12_1 | *koko | breast (n.) | x o k | 3 | 1.00 | 3 |
| 14_2 | *bi | 1SG | b i - | 3 | 1.00 | 2 |
| 25_3 | *ki- | do (make) (v.) | k ? ? / x ? ? | 3 | 0.33 | 1 |
| 26_4 | *di:ba | house (n.) | d iː b | 3 | 1.00 | 3 |
| *46_5 | *keme- | bite (v.) | | | | |
| 47_6 | *ar-ka | back (n.) | - a r | 3 | 1.00 | 2 |
| 55_7 | *da-k-,*da-n- | burn (v.) | d a ? | 4 | 0.67 | 2 |
| 56_8 | *an- | not | - aː n | 3 | 1.00 | 2 |
| 63_9 | *tur | soil (n.) | t ? r | 3 | 0.67 | 2 |
| *64_10 | *nab-či | leaf (n.) | | | | |
| 65_11 | *pula- | red | p ? l / p ? lč | 3 | 0.67 | 2 |
| 79_12 | *pur- | blow (v.) | p ? ? | 3 | 0.33 | 1 |
| 92_13 | *kole | shade (n.) | k ? l / k ? lč | 3 | 0.67 | 2 |
| 99_14 | *kata- | hard | k a t | 5 | 1.00 | 3 |
| 111_15 | *nali | raw | ? ? l / ? ? lč | 3 | 0.33 | 1 |
| 123_16 | *kira | edge (n.) | k i r | 3 | 1.00 | 3 |
| 126_17 | *pul | cheek (n.) | p ? ? | 3 | 0.33 | 1 |
| 126_18 | *čira | cheek (n.) | č ? r | 3 | 0.67 | 2 |
| 130_19 | *ur(e)- | grow (v.) | - u r / - ʊ r | 3 | 1.00 | 2 |
| 130_20 | *ös- | grow (v.) | - iu s | 3 | 1.00 | 2 |
| 135_21 | *e- | which? | - ? - | 3 | 0.00 | 0 |
| *144_22 | *nar- | thin | | | | |
| 146_23 | *köp- | foam (n.) | k iu p / x iu p | 3 | 1.00 | 3 |
| 152_24 | *siara- | white | s iaː r | 4 | 1.00 | 3 |
| 157_25 | *mɔrɔ | woods (n.) | m ? ? | 3 | 0.33 | 1 |
| 165_26 | *diu- | four | ? ? ? | 4 | 0.00 | 0 |
| 169_27 | *ta-, te- | that | t ? - | 3 | 0.50 | 1 |
| 171_28 | *ama, eme | mother (n.) | - ? m | 4 | 0.50 | 1 |
| 171_29 | *ana, ene | mother (n.) | - a n | 3 | 1.00 | 2 |
| 175_30 | *beyin | brain (n.) | m ? ? | 3 | 0.33 | 1 |
| 176_31 | *dula- | warm | d ? ? | 4 | 0.33 | 1 |
| 181_32 | *ama, eme | female (of an animal) (n.) | - ? m | 5 | 0.50 | 1 |
| *184_33 | *mangil | forehead (n.) | | | | |
| 185_34 | *sola | left | s ? l / s ? lč | 3 | 0.67 | 2 |
| 188_35 | *e- | how? | - ? - | 3 | 0.00 | 0 |
| 190_36 | *xa- | where? | x a - | 3 | 1.00 | 2 |
| *191_37 | *tol- | spin (v.) | | | | |
| 191_38 | *tɔmu- | spin (v.) | t ɔ m | 3 | 1.00 | 3 |
| 191_39 | *pɔrɔ- | spin (v.) | p ? r | 3 | 0.67 | 2 |
| 196_40 | *magu | bad | m ? ? | 3 | 0.33 | 1 |
| 197_41 | *kap | bark (n.) | k aː p / g aː p / x aː p | 3 | 1.00 | 3 |
| *199_42 | *toga-la- | count (v.) | | | | |
| 202_43 | *to:ŕa | dust | t ? ? | 3 | 0.33 | 1 |
| 204_44 | *aba | father (n.) | - a b | 3 | 1.00 | 2 |
| *212_45 | *nogo-ɣan | green | | | | |
| 216_46 | *aba-la- | hunt (v.) | - ? b | 3 | 0.50 | 1 |
| 239_47 | *tondo, torto | straight | t ɔ ? | 3 | 0.67 | 2 |
| 249_48 | *bu-PL | 1PL pronoun | ? u - | 3 | 0.50 | 1 |
| 249_49 | *bi-PL | 1PL pronoun | b i - | 3 | 1.00 | 2 |
| 253_50 | *siara- | yellow | ? ? ? | 3 | 0.00 | 0 |
| ALL | | | | 2.76 | 0.65 | 1.67 |

Table 1.3: Results of our consistency check.

Table 1.4: Distribution of cognates across a collection of datasets

| | Transeurasian (Robbeets et al. 2021) | Sino-Tibetan (Sagart et al. 2019) | Sino-Tibetan (Zhang et al. 2019) | Indo-European (Dunn & Tresoldi 2021) |
|---|---|---|---|---|
| Major subgroups | 5 | 7 | 10 | 10 |
| Concepts | 254 | 250 | 100 | 207 |
| Cognates in at least 2 subgroups | 317 | 603 | 347 | 748 |
| Cognates in 60% of subgroups | 38 (0.15) | 47 (0.19) | 19 (0.19) | 52 (0.25) |
| Cognates in 80% of subgroups | 5 (0.02) | 23 (0.09) | 11 (0.11) | 32 (0.15) |
| Cognates in 100% of subgroups | 2 (0.01) | 19 (0.08) | 7 (0.07) | 12 (0.06) |

#### 1.5 Detailed qualitative analysis

Here we give detailed qualitative comments about irregularities in the sound correspondences for all comparisons in SI1, as well as other problems with the presentation and analysis of the data by Robbeets et al. (2021).[3]

Properly evaluating the different comparisons in SI1 proved difficult due to a variety of problems. First, the data sources for the 98 languages involved are not indicated anywhere, which makes it difficult to check that the forms cited actually exist and have the meaning listed. During our analysis, we discovered that some forms in SI1 were attributed to the wrong language (e.g. comparisons #46 and #191) and that some others are only attested at a later stage than indicated (e.g. comparisons #126 and #130). In addition, we could not easily verify the meanings of the different comparanda since in many cases the only gloss provided was a number without any corresponding reference. We presume that these were copy-pasted from the online version of Starostin et al. (2003) without the corresponding list of meanings.

Moreover, the reconstructions given in SI1 suffer from a general lack of expliciteness, comprehensiveness and consistency. For instance, several languages are missing from the correspondence tables, and it is thus impossible to evaluate the phonological regularity of cognates involving these languages. For those languages that are listed in the tables, some segments are however absent, preventing proper evaluation of cognate sets that contain such segments. For instance, long vowels are attested and reconstructed in Mongolic, Tungusic and Turkic, but they do not appear anywhere in the correspondence tables and remain unaccounted for. This explains why the Manchu vowel transcribed as *ū* or *û* (usually interpreted as *ʊ*) is missing from the tables too: it was mistakenly interpreted as a long vowel *u:* and confused with *u*, even though this mistake has already been pointed in Alonso de la Fuente (2016). Tonal distinctions in Japonic are also completely ignored.

In addition, the proposed proto-forms often contradict not only the listed correspondences but also the known history of the languages involved, and they generally substantially differ from the standard reconstructions in the different families. The state of the art of the historical linguistics of each language family is too often ignored.

[3] Linguistic forms are given as they appear in the SI1, but we tacitely correct typos in our comments. Numbered sound correspondences and tables refer to those found at the end of SI2. Reconstructions cited from other sources were sometimes converted to a unified transcription.

##### #12 'breast (n.)' *koko

*Comments* According to SI2 Table 3.11, corr. #21, the pTEA reconstruction should be *xoko rather than *koko. According to SI2 Table 3.5, pTg *x- > Man. *w-* or zero (not *x-*) and NanB. *x-* or *s-* (not *k-*). This incongruence in sound correspondences may be pointing to the existence of two different words at the Tungusic level.

**Mongolic** *kökö-n

1. Kh. *xöx*
2. Bur. *xüxe(n)*
3. Kalm. *kökn*
4. ShYu. *hkön* ~ *hgön*
5. Bao. *kugo*
6. Dgx. *gogo*
7. Mnh. *kugo*
8. Huz. *kugo*
9. MMoSH *kokan*
10. MMoMuq. *köken*

*Comments* Nugteren (2011: 425) *köken 'breast'.

**Tungusic** *xökö-n

1. Udi. *oko*
2. NanA. *kun*
3. Ork. *qu:(n)* ~ *qo:(n)*
4. Jur. *gugu* ~ *xuxun*
5. Man. *xuxun*
6. Xib. *xuxuŋ*
7. Evn. *okan*

**Turkic** *kökü-$r_2$

1. BTat. *kökräk*
2. CC *köksug*
3. NAlt. *kögüs*
4. OT *kögüz*
5. SAlt. *kögüs*
6. Aze. *köküs*
7. Bas. *kükräk*
8. Chu. $kə^{w}gə^{w}r$
9. CTat. *koküs*
10. Gag. *gü:s*
11. KKal. *kökürek*
12. KBal. *kökürek*
13. Kar. *kökräk*
14. Kaz. *kökrek*
15. Khk. *kögɘs*
16. Khl. *ki:es*
17. Kir. *kökürök*
18. Kum. *kökürek*
19. Nog. *kökirek*
20. Sal. *küpräχ*
21. WYug. *gös*
22. KTat. *kükräk*
23. Tks. *göγüs*
24. Tkm. *kükrek*
25. Uyg. *kökräk*
26. Uzb. *kọkrak*

### #14 '1SG' *bi

*Comments* This is the same etymon as #249 *bi-PL '1PL pronoun' without the plural suffix.

**Mongolic** *bi

1. Kh. *bi*
2. Bur. *bi*
3. Kalm. *bi*
4. Oir. *bi*
5. Kmg. *bi*
6. Dag. *bi*
7. ShYu. *bə*
8. Bao. *bə*
9. Dgx. *bi*
10. Mnh. *bi*
11. Huz. *bu*
12. Kgj. *bi*
13. Mog. *bi*
14. MMoSH *bi*
15. MMoMuq. *bi*

*Comments* Nugteren (2011: 281) suggests a back vowel *bï to account for the relationship with plural (inclusive) *bïda.

**Tungusic** *bi

1. Hez. *bi* (*min-*)
2. Orc. *bi* (*min-*)
3. Udi. *bi* (*min(ə)-*)
4. NanA. *mi* (*min-* ~ *mimbi-*)
5. KU *mi* (*min-*)
6. NanB. *bi* (*min-*)
7. Ork. *bi:* (*min-*)
8. Ulc. *bi* (*min-*)
9. Jur. *bi* (*min-*)
10. Man. *bi* (*min-*)
11. Xib. *bi* (*min-*)
12. Evn. *bi:* (*min-*)
13. Sol. *bi* (*min-*)
14. Evk. *bi:*
15. StEvk. *bi* (*min-*)
16. SEChi. *bi*
17. NETut. *bi:*
18. NETur. *bi*
19. Neg. *bi* (*min-*)
20. Orq. *bi:* (*min-*)

**Turkic** *bi

1. BTat. *min*
2. Bas. *min*
3. Dol. *min*
4. Khk. *min*
5. KTat. *min*
6. Yak. *min*

*Comments* Though it cannot be observed in the forms provided, the majority of Turkic languages have a vowel *-e-* or *-ä-*, which led other specialists to reconstruct pTk *ben (e.g. Janhunen 2013: 219). Thus, according to SI2 Table 3.10, a form like Kir. *men* can only go back to a proto-form with *-e-. SI2 assumes instead that only *-i-* forms and Chu. go back to *bi-, and that other forms go back to *be-, a distinct root with the same consonant and the same meaning.

##### #25 'do (make) (v.)' *ki-

*Comments* No account is provided for the coda *-l in pTk and *-r in pJ. In addition, pTk *-ï is not compatible with the possible Japonic proto-forms (corr. #40.1)

**Japonic** **ker-, **kəir-, **kair-, **kiər-, **kiar-

1. Yng. *khírun*

*Comments* Yng. *khírun* is listed in SI1 but is not discussed in SI2. It has no cognate in any other Japonic language, and its etymology is unknown. In any case, the lack of mutation in the initial consonant indicates that the vowel cannot come from pJ *i (compare with pJ *kir- > *ccun* 'cut'), the expected outcome of pTEA *i (corr. #40).

**Mongolic** *ki-

1. Kh. *xiy-*
2. Bur. *xe-*
3. Kalm. *ke-*
4. Oir. *ke-*
5. Kmg. *ki-*
6. Dag. *kiː-*
7. ShYu. *gə*
8. Bao. *кә-*
9. Dgx. *giə*
10. Mnh. *ge*
11. Huz. *gə*
12. Kgj. *gi*
13. Mog. *kə-*, *ki-*
14. MMoSH *ki-*
15. MMoMuq. *ki-*

*Comments* Nugteren (2011: 415) *ki- 'to do'.

**Turkic** *kïl-

1. BTat. *ḳïl*
2. NAlt. *ḳïl*
3. OT *kïl*
4. Dol. *gïn*
5. Kar. *ḳïl*
6. Kir. *ḳïl*
7. WYug. *ḳïl*
8. Tof. *kïl*
9. Tuv. *ḳïl*
10. Uyg. *ḳil*
11. Uzb. *ḳil*
12. Yak. *gïn*

##### #26 'house (n.)' *diːba

*Comments* SI2 Table 3.12 does not include long vowels.

**Japonic** *(y)ipi

1. Jpn *ie*
2. OJ *ipye*
3. Kag. *ie*
4. Kos. *je*
5. Kum. *ije*
6. Fuk. *ie*
7. Hac. *je*

*Comments* Martin (1987: 421): ?*ipa-Ci, ?*ipi-a, ?*ipo 2.3 'house'. The reconstruction of initial *y- is not supported by any internal evidence (SI2: 12).

**Koreanic** *cip < ? *cipi

1. HH *cip*
2. SHG *cip*
3. NHG *cip*
4. SPA *cip*
5. NPA *cip*
6. JJ *cip*
7. SJL *cip*
8. NJL *cip*
9. SGS *cip*
10. NGS *cip*
11. SCC *cip*
12. NCC *cip*
13. GG *cip*
14. GW *cip*
15. MK *tip* ~ *cip*

*Comments* Martin (1996: 45) *cipɨ, *cipʌ ‘house’; Vovin (2010: 171) *cipɨ ‘house’. The final vowel cannot be *i, which does not undergo apocope in that context.

**Tungusic** *ji:b

1. Hez. *ʤo*
2. Orc. *ʤu̯:(g)* ~ *ʤu̯bba*
3. Udi. *ʤugdi*
4. NanA. *ʤo:(g)-*
5. KU *ʤo:*
6. NanB. *ʥo:* ~ *ʥo:g*
7. Ork. *duhu* (*dug-*)
8. Ulc. *ʤo:* (*ʤoɣ-*)
9. Evn. *ʤu:*
10. Sol. *ʤu*
11. Evk. *ʥu:g*
12. StEvk. *ʤu:*
13. SEChi. *ďukča:* / *ďu*
14. NETut. *ďu:*
15. NETur. *dû*
16. Neg. *ʤo:*
17. Orq. *ʤu:*

##### #46 ‘bite (v.)’ *keme-

**Japonic** *kam-

1. Jpn *kam-*
2. OJ *kam-*
3. Yng. *khàmun*
4. Kag. *kan*, *kantsu?*
5. Kos. *kamu*

*Comments* Martin (1987: 703): *kama- B ‘chew, bite, eat’
*Missing cognates* 1. Hac. *kam-*; 2. Fuk. *kam-*; 3. Kum. *kam-*; 4. Yam. *xam-*; 5. Ynm. *ham-*; 6. Ira. *kam-*; 7. Ish. *kam-*; 8. Htm. *kam-*.

**Koreanic**

1. MK *kamuŋ*

*Comments* There is no such Koreanic form, and <*ŋ*> is never used for transcribing Koreanic in Robbeets et al. (2021). It has probably been erroneously copied and pasted from a Japonic language.

**Mongolic** *keme-

1. Bao. *kaməl-*
2. MMoMuq. *kemile-*

*Comments* Nugteren (2011: 410) *kemile- ‘to gnaw’, “[p]robably derived from a noun *kemi ‘soft bone’”.

##### #47 ‘back (n.)’ *ar-ka

**Mongolic** **arka

1. Kmg. *ara*
2. Dag. *arxen*

*Comments* Nugteren (2011: 274) reconstructs *aru- ‘back, posterior side’ and argues that Dag. *arxen* is either a diminutive form from *aru-, or borrowed from Tungusic.
*Missing cognates* 1. MMoSH *aru*; 2. MMoMuq. *āru-dur*; 3. Kh. *aru*; 4. Bur. *ara*; 5. Kalm. *ar*; 6. ShYu. *a:r*.

**Tungusic** **arkan

1. Orc. *akka(n)*
2. Udi. *aka*

*Comments* SI2 Table 3.5 does not include sound correspondences involving Orc. *-kk-* and Udi. *-k-*. It is only with the help of additional cognates, like for example Literary Ewenki *arka/n* 'back (anat.)', that we can posit pTg *-rk-, which regularly yields Orc. *-kk-* and Udi. *-k-*.

**Turkic** **arka

1. BTat. *arɣa*
2. CC *archa*
3. NAlt. *arḳa*
4. OT *arḳa*
5. Aze. *arχa*
6. Bas. *arḳa*
7. CTat. *arḳa*
8. Gag. *arka*
9. KKal. *arḳa*
10. Kar. *arka*
11. Kaz. *arḳa*
12. Khk. *arɣa*
13. Khl. *arḳa*
14. Kir. *arḳa*
15. Kum. *arḳa*
16. Nog. *arḳa*
17. Sal. *arɣa*
18. WYug. *arḳa*
19. KTat. *arḳa*
20. Tks. *arka*
21. Tkm. *arḳa*
22. Uyg. *arḳa*
23. Uzb. *ɔrḳa*

##### #55 'burn (v.)' *da-k-, *da-n-

*Comments* This cognate set mixes transitive ('to burn something') and intransitive ('to get burned') forms.

**Japonic** *tak-

1. Jpn *yake-*
2. OJ *yake-*

*Comments* Martin (1987: 784): *daka-Ci- A 'get burned/roasted'. The given pJ form *tak- is transitive and is a different root.
*Missing cognates* Cognates are widely attested throughout Japonic but are not always listed in the sources. The root *yak- is even more widely found. 1. Kum. *jake-*; 2. Yam. *jehe-*; 3. Shu. *'jaki-* 1; 4. Ynm. *'jaʁii-*; 5. Ira. *jaki-*; 6. Ish. *jaki-*.

**Koreanic** *tʌhʌ- < *takʌ-

1. HH *thaywu-*
2. SHG *thaywu-*
3. NHG *thaywu-*
4. SPA *thaywu-*
5. NPA *thaywu-*
6. SGS *thaywu-*
7. NGS *thaywu-*
8. SCC *thaywu-*
9. NCC *thaywu-*
10. GG *thaywu-*
11. GW *thaywu-*
12. MK *tho-*

*Comments* Vovin (2010: 114) *hʌtʌ-, *tʌhʌ- 'to burn'. It could also be reconstructed as *kʌtʌ- or *tʌkʌ-. The bare root has an intransitive meaning, and the modern forms listed are derived from the causative form (MK *thóyGwó-*).

**Tungusic** **da-

1. Man. *da-*

*Comments* The Man. form is isolated within Tungusic.

**Turkic** *ya-k-

1. BTat. *yan-*
2. Aze. *yan*
3. Bas. *yan*
4. Chu. *śun*
5. CTat. *yan*
6. Gag. *yan*
7. KKal. *žan*
8. Kar. *yan*
9. Kaz. *žan*
10. Kir. ? *ǰan*
11. Kum. *yan*
12. Nog. *yan*

13. KTat. *yan*
14. Tks. *yan*
15. Tkm. *yan*
16. Uyg. *yan*
17. Uzb. *yɔn*

*Comments* The pTk reconstruction *ya-k- in SI2 is the transitive form, whereas intransitive forms are used in SI1.

##### #56 'not' *an-

*Comments* In Vovin (2018: 126-131), it is argued that Old Korean *an-ti* and *an-kʌy* could be the origin via borrowing of, among others, Man. *akû*. This explanation would account for the two Tungusic forms cited as well.

**Japonic** *ana-

1. Jpn *(a)-na-i*
2. OJ *(a)zu*
3. Yam. *aɴ*
4. Asa. *aɴ*
5. Yor. *aɴnu*
6. Ynm. *aɴ*
7. Shu. *aɴ*
8. Ira. *(a)n*
9. Ish. *anu*
10. Htm. *anu*
11. Yng. *-anu-n*
12. Kag. *n-*
13. Kos. *n-*
14. Kum. *n-*
15. Fuk. *n-*
16. Hac. *nnaka-*

**Koreanic** *an-

1. HH *ani-*
2. SHG *ani-*
3. NHG *ani-*
4. SPA *ani-*
5. NPA *ani-*
6. JJ *ani-*
7. SJL *ani-*
8. NJL *ani-*
9. SGS *ani-*
10. NGS *ani-*
11. SCC *ani-*
12. NCC *ani-*
13. GG *ani-*
14. GW *ani-*
15. MK *ani*

**Tungusic** *a:na-

1. Jur. *(a)kua*
2. Xib. *qu*

*Comments* The proposed pTg reconstruction cannot be derived from the Jur. and Xib. forms according to the correspondence tables.

##### #63 'soil (n.)' *tur

*Comments* According to SI2 Table 3.12, there is no sound correspondence accounting for pTg *ö and pM *-o-. None of the corr. #35, #37, or even #40d works with this particular set of cognates.

**Japonic** **toro, **tərə

1. Yor. *duru*

*Comments* Martin (1987: 391): *ntoro 2.3 'mud'; Vovin (2010: 165) *nVtoro, *mVtoro. The voiced initial is probably secondary, and the quality of the vowel cannot be determined since the word is not attested in OJ. Though the meaning 'soil' is sometimes listed, the primary meaning is usually 'mud'. Other Japonic cognates such as Fuk. *doro* and Kos. *doro* are assigned to a different cognate set (*yer).

**Mongolic** **tor

1. Dgx. *tura* 2. Kgj. *turu*, *turʁu*

*Comments* This is the same etymon as in #202 'dust' *to:rá. Nugteren (2011: 520) *toarag, *tobarag 'earth; dust, dust cloud, speck of dust', a loan from Turkic *toprak 'earth'. Dgx. *-u-* can originate from pM *o, *ö, *u, or *ü (SI2 Table 3.8), but the cognate forms in Nugteren (2011: 520) unambiguously indicate pM *o (e.g. Dag. *tʷa:rəl*, Kh. *toorog*).

**Tungusic** **tö:r

1. Evn. *tor* 2. Neg. *tu:j*

*Comments* According to SI2 Table 3.6, Evn. *-o-* and Neg. *-u-* can only go back to pTg. *-ö-. This table contains no information on the correspondence of Neg. long vowels.

#### #64 'leaf (n.)' *nab-či

**Mongolic** **nabči

1. Kh. *nabšan*
2. Bur. *namč*
3. Kalm. *namči*/*navči*
4. Oir. *nabči*
5. Kmg. *larči*
6. Dag. *ɬabʤəg*
7. ShYu. *labtɕaŋ*
8. Bao. *latʂɯn*
9. Mnh. *labdʐə*
10. Huz. *lafʤɔ*, *lar̥tʃɔ*
11. MMoMuq. *nabcin*

*Comments* Nugteren (2011: 450) *nabčïn. Bur. *namč* and Kalm. *namči* go back to a distinct proto-form *namčï.
*Missing cognates* 1. Dgx. *latʂin.*

**Tungusic** borrowing

1. Evk. *napči* (bor.) 2. Orq. *nabu̥če* (bor.)

**Turkic** borrowing

1. WYug. *lapčïk̠* (bor.)

#### #65 'red' *pula-

*Comments* pK *ɨ does not correspond to pM *u and pTg *u (corr. #39, pK *ʌ is expected instead), but to pM *ö and pTg *ö (corr. #37). Corr. #39b does not apply since the second vowel is *a in pM and pTg.

**Koreanic** *pɨl-kɨ-

1. MK *pulk-*

**Mongolic** *pula

1. Kh. *ula:n*
2. Bur. *ula:n*
3. Kalm. *ulan*
4. Oir. *ula:n*
5. Kmg. *ulān*
6. Dag. *ulān*
7. ShYu. *ɬaan*
8. Bao. *fulaŋ*
9. Dgx. *xulan*
10. Mnh. *hulang*
11. Huz. *fulaan*
12. Kgj. *fulɔ*
13. Mog. *ulōn*
14. MMoSH *xula'an*
15. MMoMuq. *hula:n*

*Comments* Nugteren (2011: 363) *hulaan.

**Tungusic** *pula

1. Hez. *fulgian*
2. Udi. *hulaligi*
3. NanB. *folgä:(n)*
4. Jur. *ful[g]ian*
5. Man. *fulgijan*
6. Xib. *fəlgian*
7. Evn. *hulańa ~ hulati*
8. Sol. *u̯la.riŋ*
9. Evk. *ula.rin*
10. StEvk. *hula-ma ~ hula-rin*
11. SEChi. *ho:lbama*
12. NETut. *xolbama*
13. NETur. *hulama*
14. Neg. *hola-ji:n*
15. Orq. *u̯la:-ren*

#### #79 'blow (v.)' *pur-

*Comments* No explanation is provided for the irregular absence of the expected final *r in pTg (corr. #29).

**Koreanic** *pulɨ-

1. HH *pwun-*
2. SHG *pwun-*
3. NHG *pwu-*
4. SPA *pwu-*
5. NPA *pwun-*
6. JJ *pwun-*
7. SJL *pwun-*
8. NJL *pwul-*
9. NGS *pwul-*
10. SCC *pwun-*
11. NCC *pwun-*
12. GG *pwun-*
13. GW *pwun-*
14. MK *pwul-*

*Comments* Martin (1996: 41) *pulɨ- 'blow'

**Tungusic** *pu:

1. Orc. *hu:-*
2. NanA. *pu:-*
3. Ork. *pu:-*
4. Ulc. *pu:-*
5. Evn. *hu:-*
6. Sol. *u:gu-*
7. Evk. *hu:wu-*
8. StEvk. *huw-*
9. SEChi. *hup-*
10. NETut. *hu:p-*
11. NETur. *huvû-*

**Turkic** **pür

1. OT *ür*
2. SAlt. *ür*
3. Bas. *ör*
4. Chu. *və$^w$r*
5. Dol. *ür*
6. KKal. *ürle*
7. KBal. *ür*
8. Kar. *ir*, *ür*
9. Kaz. *ürle*
10. Khk. *ür*
11. MChu. *ür*
12. Nog. *ür*
13. KTat. *ör*
14. Tof. *ür*
15. Tuv. *ür*
16. Yak. *ür*

#### #92 'shade (n.)' *kole

*Comments* From the correspondence pK *ɨ :: pTk *ö (corr. #37), pTg *ö is expected instead of *e.

**Koreanic** **kɨlɨmey

1. NGS *kulungci*
2. GW *kulungci*
3. MK *kulumey*

**Tungusic** **kel

1. Jur. *həlmə*
2. Man. *həlmən*
3. Xib. *xəlmən*

**Turkic** **köl

1. CC *coläge*
2. NAlt. *kölötkö etc.*
3. OT *kölige*
4. SAlt. *kölötki*
5. Aze. *kölkä*
6. Bas. *külägä*
7. CTat. *kol'ge*
8. Dol. *külük*
9. Gag. *gölgä*
10. KKal. *köleŋke*
11. KBal. *kölekke*
12. Kar. *kölege*
13. Kaz. *köleŋke*
14. Khk. *kölek*, *köletkə*
15. Kir. *kölökö*
16. Kum. *gölentgi*
17. MChu. *ke:ləg*
18. Nog. *köletki*
19. WYug. *külehke* (Mal.), *kelehki*

(Ten.)
20. Sho. *kölek*
21. KTat. *külägä*
22. Tof. *hölege*
23. Tks. *gölge*
24. Tkm. *kölege*
25. Tuv. *χölege*
26. Uyg. *köläŋkä*
27. Yak. *külük*

##### #99 'hard' *kata-

*Comments* The reconstructed root *kata- instantiates corr. #32b. (*CaCa), which implies that the expected pK reflex should be *kʌtʌ-. In turn, pK *kʌtʌ- should yield MK †*thó-* rather than *kata- > MK †*kata-*, which is not attested as such anyway.

**Japonic** *kata-

1. Jpn *kata-*
2. OJ *kata-*
3. Shu. *katasaɴ*
4. Yng. *khatan*
5. Kag. *kate*
6. Kos. *kataka*
7. Kum. *kataka*
8. Fuk. *kataka*
9. Hac. *katakja*

*Comments* Martin (1987: 831): *kata- A 'hard'

**Koreanic** *kata-

1. HH *kwut-*
2. MK *kwut-*

**Mongolic** *kata

1. Kh. *xatu:*
2. Bur. *xatu:*
3. Kalm. *xatu:*
4. Oir. *xatu:*
5. Kmg. *xatu:*
6. Dag. *kato:*
7. ShYu. *Gadu*
8. Bao. *ħdɵŋ*
9. Dgx. *qɯdun*
10. Mnh. *katoŋ*
11. Huz. *xadɔŋ*
12. Kgj. *χutuŋ*, *χuduŋ*
13. MMoMuq. *qatawu:*

*Comments* Nugteren (2011: 405) *katau 'hard', from *kata- 'to be or become hard or dry'.
*Missing cognates* 1. MMoSH *qatanggin.*

**Tungusic** **kat

1. Man. *kataŋ səmə*
2. Sol. *hat*
3. Neg. *katan*
4. Orq. *katan*

**Turkic** *kat

1. BTat. *ḳattï*
2. CC *chati*
3. NAlt. *ḳadu(:)*, *ḳadïɣ*
4. OT *ḳatïɣ*
5. SAlt. *ḳatu*
6. Bas. *ḳatï*
7. Chu. *χïdə*
8. CTat. *ḳattï*
9. Dol. *kïta:nak*
10. KKal. *ḳattï*
11. KBal. *ḳatï*
12. Kar. *ḳatï*
13. Kaz. *ḳattï*
14. Khk. *χatïɣ*
15. Kir. *ḳatu:*
16. Kum. *ḳattï*
17. MChu. *ḳa:dəɣ*
18. Nog. *ḳatï*
19. Sal. *χït(t)i*, *χïǯi*
20. WYug. *ḳatïɣ*
21. Sho. *ḳadïɣ*
22. KTat. *ḳatï*
23. Tof. *ḳaˁtïɣ*
24. Tks. *katï*
25. Tkm. *ġatï*
26. Tuv. *ḳadïɣ*
27. Uyg. *ḳattiḳ*
28. Uzb. *ḳattiḳ*
29. Yak. *kïta:naχ*

##### #111 'raw' *nali

*Comments* pTg *ia and pTk *ia correspond to pK *ia > MK *ye* (corr. #40b) and not to pK *ʌ > MK *o*.

**Koreanic** **nʌl(ʌ)

1. MK *nol*

**Tungusic** **niali

1. Hez. *nialkin*
2. Orc. *ńa:li*
3. Udi. *ńa-ligi*
4. NanA. *ńa:lo:n ~ nealo:n*
5. KU *ńalkịn*
6. NanB. *ńä:lo(n)*
7. Ork. *na:lu:*
8. Ulc. *ńe:lo(n)*
9. Evn. *ńa:lak.ča*
10. Neg. *ńali.hin*

**Turkic** **ja:lč

1. BTat. *yäš*
2. OT *yaš*
3. Aze. *yaš*
4. Bas. *yäš*
5. Gag. *yaš*
6. KBal. *ǯaš, žaš, zaš*
7. Kar. *yaš*
8. Kaz. *žas*
9. Khk. *čas*
10. Kir. *ǯaš*
11. Nog. *yas*
12. Tkm. *yāš*
13. Tuv. *čaʃ*

##### #123 'edge (n.)' *kira

**Mongolic** **kira

1. Kalm. *kirə*
2. Mnh. *ćirē*
3. Huz. *ćirē*

*Comments* Nugteren (2011: 415–416) *kirbei 'edge'

**Tungusic** **kira

1. Orc. *kia*
2. Udi. *kä*
3. NanA. *kera*
4. KU *kera-nị*
5. NanB. *kịra ~ kara*
6. Ork. *kira*
7. Ulc. *kera-*
8. Evn. *kirag, kiran*

**Turkic** **kïrgak

1. SAlt. *ḳïrï*
2. Aze. *ġïraġ*
3. Bas. *ḳïr*
4. Chu. *χərə*
5. Dol. *kïrï:*
6. Kar. *kïrïy*
7. Khk. *χïri, χïrïɣ*
8. Kum. *ḳïrïy*
9. MChu. *ḳïr*
10. Sal. *ḳïrɣï*
11. Sho. *ḳïr, ḳïrïy*
12. KTat. *ḳïrïy*
13. Uyg. *ḳir, ḳirɣak*
14. Uzb. *ḳirɣɔḳ*
15. Yak. *x́ar*

##### #126 'cheek (n.)' *čira

*Comments* This cognate set is not included in SI2. pJ *i and not *u is expected from the correspondence with pM *i and pTg *i (corr. #40).

**Japonic** **tura

1. Yam. *cïra*
2. Asa. *cɨraɴtusɨɨ, biɴtaa*
3. Ynm. *huuziraa ~ ciraabuʁuu*
4. Shu. *huuzira*

*Comments* Martin (1987: 556): *tura 2.3 'cheeks, face'. Yam. *cïra* means 'face' and not 'cheek'. Ynm. *ciraa* LLH means 'face' too, and the meaning 'cheeks' exists only in compounds, such as with *-bukuu* 'something puffy'. Ynm. *huuziraa* HHLLL, with initial *h-* instead of expected †*p-*, is probably a loan from Shu. *huuz̧ira* 0, a compound involving *huu* 0 'cheek'. Asa. *cɨraɴtusɨɨ* is also a compound, though the identity of second part is obscure, and the meaning 'cheek' cannot be attributed to the *cɨra-* part, which means 'face' when used alone (*cɨra:*).

**Mongolic** **čirai

1. Mnh. *qirai*
2. Huz. *qirai*

*Comments* Nugteren (2011: 303–304) *čïraï 'face; facial expression'. The meaning 'cheek' is not securely attested in the languages cited. Lǐ (1988: 449) has only *qiree* 'facial expression, face'.

**Tungusic** **čira

1. Hez. *čira*

##### #126 'cheek (n.)' *pul

*Comments* pJ *o and pK *o correspond to pTg *o (corr. #35) and not *u. The absence of *-r in Japonic is not accounted for.

**Japonic** *po

1. Jpn *hoho*
2. OJ *popo*
3. Kag. *fu*, *fu̥tabura*, *ho:*
4. Kos. *fu:*, *fu:ta*
5. Kum. *hobeta*
6. Fuk. *ho:tan*, *ho:benta*
7. Hac. *hoppeta*

*Comments* Martin (1987: 414): *po-po 2.3. OJ *popo* is not attested (first attestation 934).
*Missing cognates* 1. Asa. *huu* LH 'cheeks (in proverbs)'; 2. Shu. *huu* 0, *huuẓira* 0 'cheeks'.

**Koreanic** *pol

1. HH *polttayki*
2. SHG *polttayki*
3. NHG *polttayki*
4. SPA *polttayki*
5. NPA *polttayki*
6. SJL *polttayki*
7. NJL *polthayki*
8. NGS *polthayki*
9. SCC *polttayki*
10. NCC *polttayki*
11. GW *ppolttaci*

**Tungusic** **pul

1. Ork. *pulči(n)*
2. Jur. *funči*
3. Man. *fulǯin*
4. Xib. *dərəj filčin*

*Comments* The attested forms imply pTg *u and not *o, which makes the pTg irregular.

##### #130 'grow (v.)' *ös-

**Mongolic** **ös-

1. Kh. *ös-*
2. Kalm. *ös-*
3. Oir. *ös-*
4. Dag. *euse*
5. Bao. *osi*
6. Dgx. *osɯ*
7. Mnh. *wosi*
8. Huz. *wuusi*
9. Kgj. *usɯ*
10. MMoSH *os-*
11. MMoMuq. *ös-*

*Comments* Nugteren (2011: 477) *ös 'to grow'. Dag. reflects *eüs- 'to originate, arise, to be started', a distinct etymon (Nugteren 2011: 334–335).

**Tungusic** **üsə-

1. Udi. *ju-*
2. Neg. *isəw-*
3. Orq. *ju:-*

**Turkic** **ös-

1. BTat. *ö(y)s-*
2. CC *os-*
3. NAlt. *ö(:)s-*
4. SAlt. *ös*
5. Bas. *üθ*
6. Chu. *üs*
7. CTat. *ös*
8. KKal. *ös*
9. KBal. *ös*
10. Kar. *ös*, *es*
11. Kaz. *ös*
12. Khk. *ös*
13. Kir. *ös*
14. Kum. *ös*
15. MChu. *ös*
16. Nog. *ös*
17. Sho. *ös*
18. KTat. *üs*
19. Tof. *öˤs*
20. Tkm. *ös*
21. Tuv. *ös*
22. Uyg. *ös*
23. Uzb. *ọs*

##### #130 'grow (v.)' *ur(e)-

**Japonic** *ura-

1. Jpn *ure-*
2. OJ *ure-*

*Comments* Martin (1987: 556): *ura-Ci- ?B 'ripen, get ripe'. OJ *ure-* does not exist (first attestation 1603). Its late attestation and the absence of Ryukyuan cognates suggest it is a recent development.
*Missing cognates* 1. Fuk. *ururu*; 2. Kum. *ururu*; 3. Kag. *uruʔ*.

**Mongolic** *ur-

1. Kh. *urga-*
2. Bur. *urga-*
3. Kalm. *urh-*
4. Oir. *urha-*
5. Kmg. *urga*
6. Bao. *wɵr*
7. Mog. *urɣu-*
8. MMoSH *urqu-*

*Comments* Nugteren (2011: 533) *urgu 'to come up, appear (usu. of celestial bodies); to grow, sprout'.
*Missing cognates* 1. Dag. *orɣʷ-*.

**Tungusic** *ure

1. NanA. *urə-*
2. KU *urə-*
3. Ork. *urə-*
4. Ulc. *urə-*
5. Evk. *urug-*

##### #135 'which?' *e-

*Comments* pK *e and pTg *e can correspond to either pJ *a (corr. #41b.) or pJ *ə (corr. #42) but not to pJ *i or *e.

**Japonic** **i, **e

1. Jpn *dore*
2. OJ *idure*
3. Yam. *dïru*
4. Asa. *diɴ*
5. Yor. *idu*
6. Ynm. *canu*
7. Shu. *ziru*
8. Ira. *nzi*
9. Ish. *ziri*
10. Htm. *nuu*, *ziri*
11. Yng. *ndí*
12. Kag. *doi*
13. Kos. *doi*
14. Kum. *doru*
15. Fuk. *dore*
16. Hac. *doi*

*Comments* Martin (1987: 430): *intu/o-ra-Ci 3.6 'which one'

**Koreanic** **e

1. HH *etten*
2. SHG *etten*
3. NHG *etten*
4. SPA *etten*
5. NPA *etten*
6. JJ *etten*
7. SJL *etten*
8. NJL *etten*
9. SGS *etten*
10. NGS *etten*
11. SCC *etten*
12. NCC *etten*
13. GG *etten*
14. GW *etten*
15. MK *enu*

**Tungusic** **e

1. StEvk. *edynma*

##### #144 ‘thin’ *nar-

**Mongolic** *nari-

1. Kh. *nariyn*
2. Bur. *narin*
3. Kalm. *nərxn*
4. Oir. *nəren*
5. Kmg. *narin*
6. Dag. *narin*
7. ShYu. *narən*
8. Bao. *na:raŋ*
9. Dgx. *narun*
10. Mnh. *narəŋ*
11. Huz. *narin*
12. Kgj. *narɔ*
13. MMoSH *narin*
14. MMoMuq. *narin*

*Comments* Nugteren (2011: 452) *narïn ‘thin, fine (not coarse)’.
*Missing cognates* 1. Mog. *nɔrin*.

**Tungusic** **nar-

1. Man. *narhuːn*

*Comments* According to Rozycki (1994: 161), Man. *narhûn-* is a Mongolic loanword.

**Turkic** *yarï-

1. Kaz. *žara-*
2. Kir. *ǯarō*
3. Tuv. *čarɨɣ-da-*

*Comments* SI2: 52 gives meanings that differ from the concept ‘thin’ for the forms cited in this cognate set: Kaz. ‘to be poor’, Kir. ‘lean, skinny’, Tuv. ‘to spend, consume’. SI2 Table 3.9 gives no correspondence for the initial consonants involved here.

##### #146 ‘foam (n.)’ *köp-

*Comments* The correspondence pM *-ö- :: pTk *-ö- implies pK *-ɨ- and not *-e- (corr. #37).

**Koreanic** **kepɨkum, **kekɨpum

1. MK *kephwum*

*Comments* Besides MK *kèphwúm*, the forms *kephom* and *tèphwúm* are also attested. The cognates in modern dialects such as *pekhum* (SGS, JJ), *thephwum* (HH) or *pekhem* (NJL) make this form difficult to reconstruct.

**Mongolic** **köɣ-

1. Kh. *xö:s(ön)*
2. Bur. *xö:hen*
3. Kalm. *kö:sn*
4. Oir. *kö:sen*
5. Kmg. *kö:sün*
6. Dag. *xwə:s*
7. ShYu. *køwəg*
8. Bao. *Gə*
9. Huz. *koosə*
10. MMoMuq. *kö:sün*

*Comments* This cognate set confuses forms reflexes of *höesün (*höersün) ‘pus, matter’ (Nugteren 2011: 361) with reflexes of *köersün ‘foam, froth’ (Nugteren 2011: 423).

**Turkic** **köp-

1. CC *kobelek*
2. NAlt. *köbük*
3. SAlt. *köbük*
4. Aze. *köpük*
5. Bas. *kübek*
6. Chu. *kə$^{w}$bə$^{w}$k*
7. CTat. *köpük*
8. Gag. *köpük*
9. KKal. *köbik*
10. KBal. *kömük*
11. Kar. *köbük*
12. Kaz. *köbĭk*
13. Khk. *köbək*
14. Kir. *köbük*
15. Kum. *göbük*
16. MChu. *ko:β'ük*
17. Nog. *köbik*
18. WYug. *kivek*, (*kevĭk*)
19. Sho. *köbük*
20. KTat. *kübek*
21. Tof. *kö$^{ʕ}$pük*
22. Tks. *köpük*
23. Tkm. *köpür*
24. Uyg. *köpük*, *kövük*
25. Uzb. *kọpik*

#### #152 'white' *siara-

*Comments* This is the same root as #253 'yellow' *siara-. Long vowels in Tungusic and Turkic are unaccounted for in the sound correspondence tables.

**Japonic** *sero-, *sera

1. Jpn *siro-i*
2. OJ *siro-si*
3. Yam. *sirusari*
4. Asa. *sɨruuhaN*
5. Yor. *sjuusaN*
6. Ynm. *siruseN*
7. Shu. *sirusaN*
8. Ira. *ssukam*
9. Ish. *ssusaaN*
10. Htm. *ssooN*
11. Yng. *ccudari*
12. Kag. *ɕire*
13. Kos. *ɕiroka*
14. Kum. *ɕiroka*
15. Fuk. *ɕiroka*
16. Hac. *ɕirokja*

*Comments* Martin (1987: 840): *siro- B

**Koreanic** *siala-

1. HH *huy-*
2. SHG *huy-*
3. NHG *huy-*
4. SPA *huy-*
5. NPA *huy-*
6. JJ *heyyeng-*
7. SJL *hwuye-*
8. NJL *hayah-*
9. SGS *hwuyeth-*
10. NGS *hyaya-*
11. SCC *hyaya-*
12. NCC *hyaya-*
13. GG *haya-*
14. GW *huy-*
15. MK *hoy-*, *huy-*, *haya ho-*

**Tungusic** *sia:ra-

1. Hez. *čiangi*
2. Orc. *ča:gʤa(n)*
3. Udi. *caligi*, *cagʤa*
4. NanA. *ča:gʤan*
5. KU *čagʤan*
6. NanB. *ca:gʥa(n)*
7. Ork. *ta:gda*
8. Ulc. *ča:gʤa(n)*
9. Jur. *šaŋgia*
10. Man. *šaŋgijan* / *šanjan*
11. Xib. *saŋən*

**Turkic** *sia:rï-

1. Chu. *šurə*

#### #157 'woods (n.)' *mɔrɔ

*Comments* pM *mo-dun and pTg *mo: imply a monosyllabic root, but no explanation is given for the presence of a second syllable in Japonic, and no segmentation is proposed. SI2 Table 3.12 does not include long vowels, and vowel length in Tungusic is unaccounted for.

**Japonic** *morɨ-

1. Jpn *mori*
2. OJ *mori*
3. Yng. *muri*

*Comments* Martin (1987: 485): ?*mor[a-C]i 2.1 'woods, wooded place/hill, shrine woods'. OJ *mori* is not attested phonographically. However, Arisaka's First Law forbids the coexistence of pJ *o amd *ə (and *ɨ in the seven-vowel hypothesis). The reconstruction cannot thus be *morɨ-, but only *moro-, *mərə-, *mɨrɨ- (or perhaps *mərɨ- or *mɨrə-), though there are other problems too.

*Missing cognates* 1. Kum. *mori*; 2. Ira. *muï*.

**Mongolic** *mo-dun

1. Kalm. *modn*
2. Kmg. *modon*
3. Dag. *xu:rt mo:d*

*Comments* Nugteren (2011: 444–445) *modun 'tree, wood', which "may contain the (collective?) suffix *-dUn, in which case the pM root is *mo-".

**Tungusic** *mo:

1. Orc. *mo:*
2. Udi. *mo:hi bua*
3. NanB. *mo:*
4. Ork. *mo:so*
5. SEChi. *mo:ha*
6. NETut. *mo:l*
7. Neg. *mo:*
8. Orq. *mɔ:ša*

#### #165 'four' *diu-

*Comments* In SI2: 57, we read: "The initial consonant does not correspond regularly in Turkic, I do not include these forms in the etymology". Indeed, from pJ *y- :: pM *d- :: pTg *d-, we would expect pTk *y- (corr. #9) and not *t-. SI2 Table 3.12 does not include long vowels, and vowel length in Turkic is unaccounted for.

**Japonic** *yə

1. Jpn *yon*, *yo*, *yottu*
2. OJ *yo*, *yotu*
3. Yam. *juucï*
4. Asa. *juucɨ*
5. Yor. *juuci*
6. Ynm. *juuci*
7. Shu. *juuci*
8. Ira. *juucï*
9. Ish. *juucï*
10. Htm. *juuci*
11. Yng. *duuci*
12. Kag. *jottsu*
13. Kos. *jottsu*
14. Kum. *jottsu̥*
15. Fuk. *jottsu*
16. Hac. *jottsu*

*Comments* Martin (1987: 575): *də 1.1 'four'

**Mongolic** *dö-

1. Kh. *döröv(ön)*
2. Bur. *dürben*
3. Kalm. *dörvn*
4. Oir. *dörven*
5. Kmg. *dürben*
6. Dag. *durwe*
7. ShYu. *dørwen*
8. Bao. *deraŋ*
9. Dgx. *ʥiəran*
10. Mnh. *dierang*
11. Huz. *deerang*
12. Kgj. *derɔ*
13. Mog. *durbɔn*
14. MMoSH *dorben*
15. MMoMuq. *dörben*

*Comments* Nugteren (2011: 318) *dörben 'four', *döčin 'forty'

**Tungusic** *dü-gin

1. Hez. *duin*
2. Orc. *di:(n)*
3. Udi. *di:*
4. NanA. *duin*
5. KU *diin ~ diɣin*
6. NanB. *dui:(n)*
7. Ork. *ʤi:n*
8. Ulc. *dui(n)*
9. Jur. *duin*
10. Man. *duin*
11. Xib. *dujin*
12. Evn. *digen*
13. Sol. *digiŋ*
14. Evk. *digin*
15. StEvk. *dygin*
16. SEChi. *dɨgin*
17. NETut. *dɨgin*
18. NETur. *dugin*
19. Neg. *digin ~ dijin*
20. Orq. *dijin*

**Turkic** *tö:rt

1. BTat. *tört*
2. CC *tort*
3. NAlt. *tört*
4. OT *tört*
5. SAlt. *tört*
6. Aze. *dörd*
7. Bas. *dürt*
8. Chu. *təʷvattə*, *təʷvadə*, *təʷvat*
9. CTat. *dört*
10. Dol. *tüört*
11. Gag. *dört*

12. KKal. *tört*
13. KBal. *tört*
14. Kar. *dört*
15. Kaz. *tört*
16. Khk. *tört*
17. Khl. *tö:ört*, *tü:ört*
18. Kir. *tört*
19. Kum. *dört*
20. MChu. *tört*
21. Nog. *dört*
22. WYug. *dürt*, *türt*, *tört*
23. Sho. *tört*
24. KTat. *dürt*
25. Tof. *dört*
26. Tks. *dört*
27. Tkm. *dört*
28. Tuv. *dört*
29. Uyg. *tört*
30. Uzb. *tọrt*
31. Yak. *tüört*

*Comments* Tkm. has a long vowel in this word: *dö:rt* (Baskakov 1968: 283a)

##### #169 'that' *ta-, te-

*Comments* There is no correspondence pM *e (corr. #33, 34) :: pTg *a (corr. #1) :: pTk *i (corr. #40).

**Mongolic** **te-re

1. Kh. *ter*
2. Bur. *tere*
3. Kalm. *ter*
4. Oir. *tere*
5. Kmg. *ter*
6. Dag. *tərə*
7. ShYu. *tere*
8. Bao. *tə*, *tər*
9. Dgx. *tərə*
10. Mnh. *tige*
11. Huz. *te*, *tenga*, *tengi*
12. Kgj. *te*
13. Mog. *te*
14. MMoSH *tere*
15. MMoMuq. *tere*

*Comments* Nugteren (2011: 519) *tere 'that; s/he, it'.

**Tungusic** **ta-r

1. Hez. *ti*
2. Orc. *təi* ~ *ti:*
3. Udi. *təji*
4. NanA. *təj*
5. KU *tij* (*ti*)
6. NanB. *ti*
7. Ork. *tari*
8. Ulc. *te*
9. Man. *tərə*
10. Xib. *tər*
11. Evn. *tar* ~ *tarak*
12. Sol. *tari* (*ta-*)
13. Evk. *tari*
14. StEvk. *tar* ~ *tara:*
15. SEChi. *tar*
16. NETut. *tawar*
17. NETur. *ta:-*
18. Neg. *taj* (*ta:-*)
19. Orq. *tare*

**Turkic** **ti-ki

1. Bas. *tege*
2. Kir. *tigi*
3. MChu. *täɣ*
4. KTat. *tege*
5. Tuv. ? *döö*

##### #171 'mother (n.)' *ama, *eme

*Comments* This is a nursery word, which makes it not really reliable for phylogenetic inference. This is also probably the same root as #181 'female (of an animal) (n.)' *ama, *eme.

**Japonic** *əmə

1. Yam. *ʔaɴma*
2. Asa. *ʔaama*
3. Ynm. *ʔamu*
4. Shu. *ʔaɴma*

*Comments* The Ryukyuan data adduced does not support the reconstrution of a pJ vowel other than *a in initial position. The reconstruction *əmə is a mechanical projection from *omo*, an Eastern Old Japanese dialectal form, but in could also be from *omo.

**Koreanic** *ema

1. HH *ommani*
2. SHG *ommani*
3. NHG *ommani*
4. SPA *ommani*
5. NPA *ommani*
6. JJ *emeng*
7. SJL *emay*
8. NJL *emei*
9. SGS *emeng*
10. NGS *ema*
11. SCC *aymi*
12. NCC *aymi*
13. GG *emeni*
14. GW *emeni*
15. MK *am*, *em*, *emuy*

**Mongolic** *eme

1. Bao. *amo*, *amə*
2. Huz. *aama*

*Comments* Nugteren (2011: 328) *eme 'woman', but he gives different cognates: Huz. *imu*, *yæmu* 'girl', Bao. *emə* 'wife', *imə* 'woman, wife'.

**Tungusic** *eme

1. Jur. *əmə*
2. Man. *əmə*

##### #171 'mother (n.)' *ana, ene

**Japonic** **ana

1. Ira. *anna*

*Comments* This form is isolated within Japonic.

**Mongolic** **ana

1. ShYu. *ana*
2. Bao. *ana*
3. Dgx. *ana*
4. Mnh. *ana*
5. Kgj. *ana*

*Comments* Not in Nugteren (2011)

**Turkic** **ana, **eńe

1. BTat. *ĕnä*, *inä*
2. CC *anna*
3. NAlt. *ene*, *ana*, *ane*
4. SAlt. *ene*
5. Aze. *ana*
6. Bas. *inä*
7. Chu. *anne*
8. CTat. *ana*
9. Dol. *in'e*
10. Gag. *ana*
11. KKal. *ana*, <*ene*>
12. KBal. *ana*
13. Kar. *ana*
14. Kaz. *ana*
15. Kir. *ene*
16. Kum. *ana*
17. Nog. *ana*
18. WYug. *ana*
19. Sho. *ene*
20. KTat. *ana*, (*äni*)
21. Tks. *anne*, *ana*
22. Tkm. *ene*
23. Uyg. *ana*, [dial. *inä*]
24. Uzb. *ɔna*

##### #175 'brain (n.)' *beyin

*Comments* ShYu. and Hez. are not included in SI2 Table 3.5 and SI2 Table 3.5. Thus, we don't know what the origin of ShYu. *-ŋ-* is within the Transeurasian framework.

**Mongolic** ?

1. ShYu. *məŋi*

*Comments* Not in Nugteren (2011). Both Bǎo (1984) and Nugteren & Roos (1996: #41) argue that ShYu. *məŋi* is a loan from the Turkic language West Yugur *muŋe*. Its Turkic origin seems to be confirmed by the fact that *ŋ* usually only appears in syllable-coda position in the native lexicon of ShYu.. The putative correspondence made with pTk *y leads to expect a development of pM *y to ShYu. *ŋ*, but no such change is described by Nugteren (2011), and none seems to exist.

**Tungusic** ?

1. Hez. *meyi*

*Comments* Neither SI2 Table 3.5 nor SI2 Table 3.6 for Tungusic sound correspondences includes Hezhe, and it is unclear to what pTg form it could go back. The isolated attestation of this word within Tungusic may indicate it is a loanword from a Turkic language.

**Turkic** **beyiŋ

1. BTat. *miyä*
2. CC *meŋ*
3. NAlt. *mees*, (P) *nee*
4. OT *meyi*
5. SAlt. *mee*
6. Aze. *beyin*
7. Bas. *meye*
8. Chu. *mime*
9. CTat. *miy*
10. Gag. ? *beyin*
11. KKal. *miy*
12. KBal. *mïyï*
13. Kar. *miy*
14. Kaz. *mïy*
15. Khk. *mii*
16. Khl. *mein*
17. Kir. *mee*
18. Kum. *miy*
19. MChu. *miis*
20. Nog. *mïy*
21. Sal. *meŋes*
22. Sho. *mi:s*
23. KTat. *mi*
24. Tof. *me:*
25. Tks. *beyin*
26. Tkm. *beyni*
27. Tuv. *mee*
28. Uyg. *meyä*, *miŋä*
29. Uzb. *miya*
30. Yak. *meyi:*

*Comments* SI2 Table 3.9 does not include the medial -*y*- present in most of the reflexes. We assume it is the same as for initial *y*- and reconstruct pTk *y. Various alternative reconstructions have been proposed in the past in the past (e.g., *bejŋi by Starostin et al. 2003, *béñī by Clauson 1972: 348).

##### #176 'warm' *dula-

*Comments* No explanation is given for the absence of the expected *l in pK (corr. #31).

**Koreanic** *tʌ- ~ *tɨ- ~ *ta- ~ *te-

1. HH *ttattus ha-*
2. SHG *ttattus ha-*
3. NHG *ttattus ha-*
4. SPA *ttattus ha-*
5. NPA *ttattus ha-*
6. JJ *testes he-*
7. SJL *taswup-*
8. NJL *ttaswup-*
9. SGS *ttasi-*
10. NGS *ttasi-*
11. SCC *ttattusha-*
12. NCC *ttattusha-*
13. GG *ttattusha-*
14. GW *ttattusha-*
15. MK *tos-*

*Comments* Only pK *tʌs- can be reconstructed.

**Mongolic** *dula-

1. Kh. *dula:n*
2. Bur. *dula:n*
3. Kalm. *dulan*
4. Oir. *dula:n*
5. Kmg. *dula:n*
6. Dag. *dulān*
7. ShYu. *dulaan*
8. Dgx. *untʂu*
9. Kgj. *dʐamadʐu*

*Comments* Nugteren (2011: 319) *dulaan 'warm'.
*Wrong cognates* 1. Dgx. *untʂu*; 2. Kgj. *dʐamadʐu*.

**Tungusic** *du:l-

1. Jur. *duluu*
2. Evn. *dụ:l-*, *du:lan*

**Turkic** *yïlï

1. NAlt. *yïlïɣ*
2. OT *yïlïɣ*
3. SAlt. *d'ïlu*
4. Bas. *yïlï*
5. KKal. *žïllï*
6. KBal. *ǰïlï*
7. Kar. *yïlï*
8. Kaz. *žïlï*
9. Khk. *čïlïɣ*
10. Kir. *ǰïluu*
11. Kum. *yïlï*
12. MChu. *čələɣ*
13. Nog. *yïlï*
14. WYug. *yilïɣ*
15. Sho. *čïlïɣ*
16. KTat. *ǰïlï*
17. Tof. *čïlïɣ*
18. Tks. *ïlïk*
19. Tkm. *yïlï*
20. Tuv. *čïlïɣ*
21. Uyg. *illiḳ*, dial. [*žiliḳ*]
22. Uzb. *iliḳ*
23. Yak. *sïla:s*

##### #181 'female (of an animal) (n.)' *ama, *eme

*Comments* This is the same root as #171 'mother (n.)' *ama, *eme. There is no explanation for the absence of an initial vowel in Japonic. pM. *e-, pTg *e- and pTk *e- ought to correspond to pK *e-, not *a- (corr. #41b, #42).

**Japonic** *me

1. Jpn *mesu*
2. OJ *mye*
3. Yam. *mïï*
4. Asa. *mɨɨmuɴ*
5. Yor. *mii*
6. Ynm. *miimuɴ*
7. Shu. *miimuɴ*
8. Ira. *miimunu*
9. Ish. *miimunu*
10. Htm. *miimu*
11. Yng. *mii+munu*
12. Kag. *men*
13. Kos. *mesu*
14. Kum. *mesu̥*
15. Fuk. *mettɕo*
16. Hac. *mesu*

*Comments* Martin (1987: 474): *miCa ?1.3b 'female'

**Koreanic** **amh

1. HH *amkhes*
2. SHG *amkhes*
3. NHG *amkhes*
4. SPA *amkhes*
5. NPA *amkhes*
6. JJ *amkhe*
7. SJL *amkhe*
8. NJL *amkhe*
9. SGS *amkhe*
10. NGS *amkhe*
11. SCC *amkhe*
12. NCC *amkhe*
13. GG *amkhes*
14. GW *amkhes*
15. MK *am*

*Comments* The MK form is *ámh*, which is congruent with the aspiration in the modern dialect forms.

**Mongolic** *eme

1. Kh. *em*
2. Bur. *eme*
3. Kalm. *em*
4. Dag. *əmɣun*
5. ShYu. *eme*
6. Bao. *ɛmə*
7. Dgx. *əmə*
8. Kgj. *eme*
9. MMoSH *eme*
10. MMoMuq. *eme*

*Comments* Nugteren (2011: 328) *eme 'woman'

**Tungusic** *eme

1. Jur. *əmilə*

**Turkic** *eme

1. Chu. *ama*

##### #184 'forehead (n.)' *mangil

**Mongolic** **maŋlai

1. Kh. *magnay*
2. Bur. *magnay*
3. Kalm. *maŋna*
4. Oir. *maŋna:*
5. Kmg. *mannay*
6. Dag. *mangil*
7. ShYu. *maŋlii*
8. Bao. *maŋləi*
9. Dgx. *manləi*
10. Mnh. *manglai*
11. Huz. *maŋlii*, *molii*
12. MMoSH *manglai*, *man*
13. MMoMuq. *manqlai*

*Comments* Nugteren (2011: 441) *maŋlaï 'forehead' ("[t]he central languages developed from an assimilated form *maŋnaï").

**Tungusic** **maŋgVl

1. Sol. *maŋgil*
2. Orq. *maŋge:la*

*Comments* According to Poppe (1931: 58a), Sol. *maŋgil* was borrowed from Dag. *mangil*, and according to Li & Whaley (2009: 544), Orq. *maŋge:la* is also a loanword from a Mongolic language.

**Turkic** **maŋlai

1. Bas. *maŋlay* (bor.)
2. CTat. *maŋlay* (bor.)
3. KKal. *maŋlay* (bor.)
4. KBal. *maŋïlay* (bor.)
5. Kar. *maŋlay* (bor.)
6. Kaz. *maŋday* (bor.)
7. Kir. *maŋday* (bor.)
8. Kum. *maŋalay* (bor.)
9. Nog. *maŋlay* (bor.)
10. KTat. *maŋgay* (bor.)
11. Tkm. *maŋlay* (bor.)
12. Uyg. *maŋlay* (bor.)
13. Uzb. *maŋlay* (bor.)

##### #185 'left' *sola

**Mongolic** **solagai

1. Oir. *solha:*
2. Kmg. *salagay*
3. Dag. *solgi:*
4. ShYu. *soloʁui*
5. Bao. *jələG*
6. Dgx. *soɣəi*, *coGi*
7. Mnh. *serghai*
8. Huz. *sulighui*

*Comments* Nugteren (2011: 500) *solagaï 'left, left hand side'
*Missing cognates* 1. Kh. *solgoy*; 2. Kalm. *solɣa*.

**Tungusic** ?

1. Xib. *sölhu*

*Comments* The Xib. form is most likely a graphic adaptation of *ɕœlχu* (Lǐ & Zhòng 1986: 144a), but cf. *syolχo* in Kubo et al. (2011a: 33), *śɑʁo* in Zikmundová (2013: 222). It is isolated in Tungusic and there is no trace of it in Man., where the regular term is *hashû*, for which cognates are attested in some modern dialects (Yamamoto 1969: 126b; Kim et al. 2008: 106b)). Provided there is such a sound in Xib., *ö* does not appear in SI2 Table 3.6. It is thus unclear to what sound would it correspond in pTg. In order to regularly correspond to pM *o and pTk *o, we would need pTg *o (corr. #43), which would yield Xib. *o* and not *ö* (SI2 Table 3.6).

**Turkic** **so:l

1. BTat. *sol*
2. CC *sol*
3. NAlt. *sol*
4. OT *sol*
5. SAlt. *sol*
6. Aze. *sol*
7. Bas. *hul*
8. Chu. *sulaɣay* (bor.)
9. CTat. *sol*
10. Gag. *sol taraf*
11. KBal. *sol*
12. Kaz. *sol*
13. Khk. *sol*
14. Kir. *sol*
15. Kum. *sol*
16. MChu. *sol*
17. Nog. *sol*
18. WYug. *sol*, (*sul*, *söl*)
19. Sho. *sol*
20. KTat. *sul*
21. Tks. *sol*
22. Tuv. *solaɣay* (bor.)
23. Uyg. *sol*

*Comments* The reconstruction with a long vowel is based on Tkm. *so:l* (Baskakov 1968: 584a), but this form is missing from the cognate list.

##### #188 'how?' *e-

**Japonic** **i, **e

1. Jpn *dou*
2. OJ *ika*
3. Yam. *xjasi*
4. Asa. *ʔissjɨɨ*
5. Yor. *icjasi*
6. Ynm. *caɴguτu*
7. Shu. *canugutu*
8. Kag. *dogen*
9. Kos. *doː*
10. Kum. *doː*
11. Fuk. *dogen*
12. Hac. *dogon*

**Koreanic** **e

1. HH *ettehkey*
2. SHG *ettehkey*
3. NHG *ettehkey*
4. SPA *ettehkey*
5. NPA *ettehkey*
6. JJ *ettekkhey*
7. SJL *ekhey*
8. NJL *eccicako*
9. SGS *ecci*
10. NGS *wuccey*
11. SCC *eccey*
12. NCC *eccey*
13. GG *etekhay*
14. GW *eccay*
15. MK *este*

**Tungusic** ?

1. Xib. *ərai*

*Comments* According to Kubo et al. (2011b: 55), Xib. *ərai* means 'what is this?'. It is a contraction of *ərə* 'this' and *ai* 'what?' and it appears in expressive collocations like for example *ərai χayriN* 'too bad'. The root is thus a demonstrative and not a manner interrogative.

##### #190 'where?' *xa-

**Mongolic** **kaɣa

1. Bur. *xaːna*
2. Kalm. *xama*
3. Oir. *xamaː*
4. Kmg. *xaana*
5. Dag. *xaːnə*
6. ShYu. *xana*
7. Bao. *ħala*
8. Dgx. *qala*
9. Mnh. *angji*
10. Huz. *anji*, *anjii*
11. Kgj. *χana*
12. Mog. *qana*
13. MMoSH *qa'a*
14. MMoMuq. *qaːna*

*Comments* Nugteren (2011: 395) *kaa, *kaana 'where?'.
*Missing cognates* 1. Kh. *xaa.*

**Tungusic** **xai-

1. Hez. *ia-du*
2. Orc. *i-du̱*, *jaː-la*
3. Udi. *je-du*
4. NanA. *haj-do ~ haj-du*
5. KU *eːdu̱ ~ ī̱du̱*
6. NanB. *hai̱-do*
7. Ork. *hai-du*
8. Ulc. *χaj-do*
9. Man. *aibi ~ aiba-də ~ aibi-də*
10. Xib. *ja-va*
11. Evn. *i-du*
12. Sol. *i-lə*
13. Evk. *i-ləː*, *iːr-bə*
14. StEvk. *iː-duː*
15. SEChi. *idu*
16. NETut. *iːdu*
17. NETur. *idu*
18. Neg. *iː-duː*
19. Orq. *i-du*

*Comments* The pronominal base *xai- (some languages point rather to *xia-, according to SI2 Table 3.6, though this is a general problem in Tungusic linguistics for which we have no satisfactory solution) acquires the meaning 'where' only when it has the (secondary) dative-locative case *-dOO attached to it.

**Turkic** **kay-

1. BTat. *ḳayda*
2. CC *chayda*, *kani*
3. NAlt. *ḳayt*, *ḳayda*
4. OT *ḳayuda*, *ḳanda*
5. SAlt. *ḳayda*
6. Aze. *harada*
7. Bas. *ḳayδa*
8. Chu. *əśta*
9. CTat. *ḳayda*
10. Dol. *kanna*
11. KKal. *ḳayda*
12. KBal. *ḳayda*, ? *ḳalayda*
13. Kar. *kayda*
14. Kaz. *ḳayda*
15. Khk. *χayda*
16. Khl. *ḳa:(nï)*
17. Kir. *ḳayda*
18. Kum. *ḳayda*
19. MChu. *ḳayda*
20. Nog. *ḳajda*
21. Sal. *ḳayda*, *ḳayta*
22. WYug. *ḳayta*, (*ḳayda*), *ḳan*
23. Sho. *ḳayda*
24. KTat. *ḳayda*
25. Tof. *ḳayda*
26. Tuv. *ḳayda*, *ḳaya:*
27. Uyg. *ḳeni*
28. Uzb. *ḳayerda*
29. Yak. *χanna*

##### #191 'spin (v.)' *pɔrɔ-

*Comments* pM *-o- and pTg *-o- do not correspond to pTk *-ö- (corr. #35–36).

**Mongolic** **por-

1. Kalm. *ə:r-*
2. Dag. *ə:r*
3. Dgx. *furəu*

*Comments* The intended cognate set here corresponds to *horïa 'to bind, wind, spin, wrap' in Nugteren (2011: 360), but the forms provided in SI1 do not originate from this proto-form, the correct reflexes being Dgx. *xoro-* and Kalm. *orax* (no Dag. cognate). The long vowel in Dag. *ə:r* and Kalm. *ə:r-* points to a sequence *ege, which would be incompatible with the other formations. Dgx. †*furəu-* should be corrected to *fura-* 'turn' (Mǎ & Chén 2012: 115), a form originating from *hurban 'turn (around)' (Nugteren 2011: 365), an unrelated etymon.

**Tungusic** **poro-

1. Man. *foro-*
2. Orq. *ɔrɔl-*

*Comments* According to Rozycki (1994: 79), Man. *foro-* is a Mongolic loanword.

**Turkic** **pör-

1. Yak. *ör*

*Comments* Yak. *-ö-* can only come from pTk *-ö- according to SI2 Table 3.10.

##### #191 'spin (v.)' *tol-

*Comments* pK *-o- does not correspond to pJ *-ə- (corr. #35–36).

**Japonic** **yər-

1. Kum. *joru*

*Comments* Martin (1987: 787): *dəra- B 'twist'
*Missing cognates* 1. OJ *yo$_2$r-* 'spin (a thread)'; 2. Jpn *yor-* 'spin (a thread)'; 3. Yam. *'jur-* 'spin (a thread)'; 4. Ira. *jur-*.

**Koreanic** **tolʌ-

1. HH *tolli-*
2. SHG *tolli-*
3. NHG *tolli-*
4. SPA *tolli-*
5. NPA *tolakamcci-*
6. JJ *tolli-*
7. SJL *tolli-*
8. NJL *tolli-*
9. SGS *tolli-*
10. NGS *tolli-*
11. SCC *tolli-*
12. NCC *tolli-*
13. GG *tolli-*
14. GW *tolli-*
15. MK *tol-*

*Comments* Vovin (2010: 127) *tolʌ-. The MK verb *twŏl-* is intransitive and means 'turn'. The MK form given in SI1 *tol-* is a mistake for *twŏl-*, whose vowel can only go back to pK *o. This is also supported by the reflexes in modern Korean dialects, which all unambiguously point to pK *o.

**Tungusic** —

1. Orq. *tolli-*

*Comments* There is no such form in Orq.. The most likely explanation is that there has been an error while copy-pasting forms in the neighbouring column where Korean data is found.

##### #191 'spin (v.)' *tɔmu-

*Comments* pM *-o- and pTg *-o- do not correspond to pJ *-u- (corr. #35–36).

**Japonic** *tumu

1. Jpn *tumug-*
2. OJ *tumug-*
3. Shu. *çing-*
4. Kag. *tsumuʔ*
5. Kos. *tsumugu*
6. Fuk. *tsumugu*
7. Hac. *tsumugu*

*Comments* Martin (1987: 775): *tumu$_{2.4}$-n[a]-ka- 'spin, make into yarn'
*Comments* The Ryukyuan forms unambiguously point to pJ *u and not *o (e.g. Shu. *çing-* and not †*tung-*).
*Missing cognates* 1. Yam. *cïɴg-* 'spin a thread with a spindle'; 2. Asa. *cïɴg-* 'spin'; 3. Ynm. *ciɴg-* 'spin'; 4. Ira. *tsïmk-*; 5. Ish. *tsïn-*.

**Mongolic** *tomu-, *tamu-

1. Kh. *tomo-*
2. Bur. *eryu:l-*
3. Oir. *tomo-*
4. Kmg. *tomo-*
5. ShYu. *tomə*
6. Bao. *təm*
7. Dgx. *tamu*
8. Huz. *tamu*
9. Kgj. *tumu*
10. MMoSH *tamu-*
11. MMoMuq. *tomu-/tamu-*

*Comments* According to SI2 Table 3.8, Dgx. *a* goes back only to pM *a. Other specialists, unable to decide about the vocalism, cautiously reconstruct both possibilities, cf. Nugteren (2011: 511) *tamu-, *tomu- (*toma-) 'to rub; to twist or spin thread or rope'. The Bur. form *eryu:l-* is obviously not cognate, and it should be replaced with *tomoxo* (Nugteren 2011: 511).
*Missing cognates* 1. Kalm. *tömx-* 'twist'.

**Tungusic** tomu-

1. Orc. *tompo-*
2. Udi. *tompo-*
3. NanA. *tompo-*
4. NanB. *tomfo-*
5. Ork. *tokpo-*
6. Ulc. *toŋpo-*
7. Evn. *tọmka-*
8. Sol. *tɔŋɔn-*
9. StEvk. *tomko-*
10. Neg. *tomko-*

##### #196 'bad' *magu

*Comments* The diphthong in Korean is unaccounted for (corr. #32). Furthermore, the expected correspondence of pK *-k- (-h-) and pTk *-k- is pM *-k- (corr. #14) and not *-ɣ-. The pTEA putative reconstruction *magu suggests corr. #16, but the Turkic evidence points to pTk *k and not *g.

**Koreanic** **maykay-

1. GW *mayhayta*

*Comments* There is no correspondence table for Korean dialects. Since there is no pK *h in the Transeurasian framework, we can tentatively reconstruct *k. The diphthong *ay* in the first syllable is unexpected and it is unclear what its origin might be. A plausible etymology is that it is related to MK *mànghòtá* 'to perish, to go to ruin' and P'yŏngan *mangheta* 'bad'. This would invalidate the comparison with other languages since it is a Sino-Korean loanword (亡).

**Mongolic** **maɣu

1. Kh. *mu:*
2. Bur. *mu:*
3. Kalm. *mu*
4. Oir. *mu:*
5. Kmg. *mu:*
6. Dag. *mo:*
7. ShYu. *muu*
8. Bao. *muŋ*
9. Dgx. *mao*
10. Mnh. *mao*
11. Huz. *muu*
12. Kgj. *mau*
13. MMoSH *mao'u(n)*, *mao'ui*
14. MMoMuq. *mu:*

*Comments* Nugteren (2011: 442) *mau- 'bad'. According to SI2 Table 3.7, pM *-g- is never lost in the daughter languages, so the consonant should be pM *ɣ instead, even though it does not appear in SI2 Table 3.7 but only in SI2 Table 3.11. Likewise, SI2 Table 3.8 doesn't account for vowel contractions after the loss of medial consonants. Janhunen (2013: 216) assumes *-x-, rather than *-g-, which would correspond to *-ɣ- in the Transeurasian framework.

**Turkic** **bak-, **biak-

1. Tof. *baˤḳ* (*baˤhay*)
2. Tuv. *baɣay*, (*baḳ*)

*Comments* SI2 Table 3.9 indicates that the final consonant should be pTk *k and not *g.

##### #197 'bark (n.)' *kap

*Comments* SI2 Table 3.12 does not include long vowels.

**Japonic** *kapa

1. OJ *kapa*
2. Yam. *xïɴxo*
3. Asa. *kɨɴkoo*
4. Yor. *siinuhoo*, *hiinuhoo*
5. Ynm. *kiinuhaa*
6. Shu. *kiinukaa*
7. Ira. *kiinukaa*
8. Ish. *kiinukaa*
9. Htm. *kiinukaa*
10. Yng. *khaa*
11. Kag. *kawa*
12. Kos. *ka:*
13. Kum. *kawa*
14. Fuk. *kawa* ~ *kinokawa*
15. Hac. *kawa*

*Comments* Martin (1987: 575): *kapa 2.3 'skin, fur' OJ *kapa* is not attested phonographically.
*Missing cognates* 1. Jpn *(ki no) kawa.*

**Koreanic** *kap(ʌ)-k

1. HH *namwukkepteyki*
2. SHG *namwukkepteyki*
3. NHG *namwukkepteyki*
4. SPA *namwukkepteyki*
5. NPA *namwukkepteyki*
6. JJ *kepcwuk*
7. SJL *kkepcil*
8. NJL *kkepteyki*
9. SGS *kkepteyki*
10. NGS *kkepttayki*
11. SCC *kkeptey*
12. NCC *kkepwul*
13. GG *kkepwul*
14. GW *kkemwuli*
15. MK *kepcil*

*Comments* The reconstruction with *a and not *e is not supported by the Koreanic evidence but only by external evidence (SI2: 28).

**Turkic** *ka:p-ïk

1. BTat. *ḳabïḳ*
2. CC *cabuc*
3. NAlt. *ḳaɣaš*
4. Aze. *ġabïġ*
5. Bas. *ḳabïḳ*
6. Chu. *χubə*
7. CTat. *ḳabuḳ*
8. Gag. *kabuk*
9. KKal. *ḳabïḳ*
10. KBal. *ḳabuḳ*
11. Kar. *kabuk*
12. Kaz. *ḳabïḳ*
13. Khk. *χaχpas*
14. Kir. *ḳabïḳ*
15. Kum. *ḳabuḳ*
16. MChu. *ḳaḳβač*
17. Nog. *ḳabïḳ*
18. Sal. *koχ*
19. Sho. *ḳaḳpaš*
20. KTat. *ḳabïḳ*
21. Tks. *kabuk*, (*aɣač*) *kabuɣu*
22. Tkm. *ġabïḳ*
23. Uyg. *ḳovzaḳ*

*Comments* The Tkm. form should be *ġa:bïḳ* (Baskakov 1968: 136a), with a long vowel which goes back to pTk.

##### #199 'count (v.)' *toga-la-

**Mongolic** **toa-la

1. Kh. *tool-*
2. Bur. *to:lo-*
3. Kalm. *to:l-*
4. Oir. *to:la-*
5. Kmg. *to:lo-*
6. Dag. *tuwa:l*
7. ShYu. *tu:la-*
8. Bao. *tɵ:la*
9. Dgx. *tolu*
10. Mnh. *tolo-*
11. Huz. *to:la-*
12. Kgj. *tula*
13. Mog. *toala-*
14. MMoSH *to'ola-*

*Comments* Nugteren (2011: 520) *toala- 'to count'.

**Tungusic** borrowing

1. Hez. *tolo-* (bor.)
2. Man. *tolo-* (bor.)
3. Xib. *tolu-* (bor.)

**Turkic** borrowing

1. Sho. *to:la-* (bor.)
2. Tks. *hesapla-*, *hesap et-* (bor.)

##### #202 'dust' *to:ŕa

*Comments* pM *-o- and pTk *-o- can only correspond to pTg *-o- (corr. #35–36), which in turn should correspond to Sol. *-o-* and not *-ɔ-*, which anyways does not appear in SI2 Table 3.6. SI2 Table 3.12 does not include long vowels.

**Mongolic** ?**torag

1. Oir. *towrog*
2. Kmg. *to:rog*
3. Dag. *tuarel*
4. Kgj. *turu*, *turʁu*

*Comments* This is the same etymon as in #63 'soil (n.)' *tur. Nugteren (2011: 520) *toarag-, *tobarag- 'earth; dust, dust cloud, speck of dust'.

**Tungusic** **t?r-

1. Sol. *tɔ:rɔl*
2. Orq. *tɔ:rag*

*Comments* Sol. and Orq. are obvious loanwords from Mongolic Li & Whaley (2009: 544). There is no vowel ɔ listed for Sol. in SI2 Table 3.6, and Orq. is not included at all. No tentative pTg reconstruction can be provided for the vowel.

**Turkic** **to:r$_2$

1. BTat. *tos*, *tosan*
2. CC *tos*
3. NAlt. *tozun* ~ *tozïn*
4. OT *toz*
5. Aze. *toz*
6. Bas. *tuδan*
7. Chu. *tuzan*
8. CTat. *toz*
9. Gag. *to:z*
10. Kar. *toz*
11. Kaz. *tozaŋ*
12. Khk. *tozïn*
13. MChu. *tozan*
14. WYug. *tos*
15. Sho. *tozun*
16. KTat. *tuzan*
17. Tks. *toz*
18. Tkm. *tozan*
19. Tuv. *do:zun*
20. Uyg. *tozaŋ*, *toz*

##### #204 'father (n.)' *aba

*Comments* pM *aba as well as the putative pTEA *aba have the structure *CaCa, which should correspond to pK *ʌpʌ (corr. #32b) and not *apa.

**Koreanic** **apa, **api

1. HH *apaci*
2. SHG *apaci*
3. NHG *apaci*
4. SPA *apaci*
5. NPA *apaci*
6. JJ *apang*
7. SJL *apssi*
8. NJL *apssi*
9. SGS *apay*
10. NGS *aypi*
11. SCC *aypi*
12. NCC *aypi*
13. GG *apa*
14. GW *apaci*
15. MK *api*, *apa*

**Mongolic** **aba

1. Kh. *a:v*
2. Bur. *aba*
3. Kalm. *a:v*
4. Oir. *a:v*
5. Kmg. *aba*
6. Bao. *awa*
7. Dgx. *aba*, *awi*
8. Mnh. *aba*
9. Huz. *aaba*
10. Kgj. *aba*

*Comments* Not in Nugteren (2011).

**Turkic** **aba, **apa

1. Sal. *apa*
2. Sho. *aba*

##### #212 'green' *nogo-ɣan

**Mongolic** **nogo-

1. Kh. *nogo:n*
2. Bur. *nogo:n*
3. Kalm. *nohan*
4. Oir. *noha:n*
5. Kmg. *nogo:n*
6. Dag. *nasən*
7. ShYu. *noʁoon*
8. Bao. *nɵGuŋ*
9. Dgx. *noɣon*
10. Mnh. *nuoghuang*
11. Huz. *nughuun*
12. Kgj. *nuʁun*
13. MMoMuq. *noga:n*

*Comments* Nugteren (2011: 461) *nogaan 'green'

**Tungusic** ? ? ?

1. Hez. *niuŋgian*
2. Orc. *ńu̜:gʤa(n)*
3. Udi. *njogje*
4. NanA. *ńoŋean*
5. KU *ńongian*
6. NanB. *noŋgä:(n)* ~ *ńoŋgä:(n)*
7. Ork. *ńo:gdo*
8. Ulc. *ńo:gʤo(n)*
9. Jur. *niəŋia*
10. Man. *niowaŋgijan*
11. Xib. *nüŋnian*

*Comments* No sound correspondences are given in SI2 Table 3.5 and SI2 Table 3.6 that would account for either the initial *ń-* in the various languages or Man. *o* :: Orc. *u* (the latter goes back to either pTg *u or *ö). Khabtagaeva (2022: 483-484) suggests that Dag. *nasun*, *nasen* is a borrowing from Sol. *nahun.* She reconstructs pTg *nagun, but at the same time admits that the base must be *ño- (following here the opinion of G. Doerfer). If pTg contains *ń-, there is no sound correspondence in SI2 Table 3.11 that would include this sound.

**Turkic** borrowing

1. Khk. *noɣan* (bor.)
2. Tuv. *noɣa:n* (bor.)

#### #216 'hunt (v.)' *aba-la-

*Comments* SI2 Table 3.12 does not include long vowels.

**Mongolic** **aba

1. Kh. *avl-*
2. Oir. *avala-*
3. Dag. *aula:*
4. MMoMuq. *abala-*

*Comments* Nugteren (2011: 189) *aba 'hunt', with the productive verbalizer *-lA:-.

**Tungusic** **aba

1. Man. *aba-la-*
2. Xib. *avələ-*

*Comments* Man. *aba-la-* has been indentified as a Mongolic borrowing (Rozycki 1994: 9), as suggested by the isolation of this etymon in Tungusic, and the fact that only the denominal verb is attested, while the base noun is not present.

**Turkic** **a:b-la-

1. CC *uula-*, *huala-*
2. OT *avla-*
3. Aze. *ova čïχ*, *ovčuluġ et*, *ovla*
4. Bas. *awla*
5. CTat. *avla*
6. Gag. *awla*
7. KKal. *awla*
8. KBal. *uwuɣa* + motion verb
9. Kar. *avla*
10. Kaz. *awla*
11. Kir. *uula*
12. Kum. *(ɣ)aw et*
13. KTat. *awla*
14. Tks. *avla*
15. Tkm. *awla*
16. Uyg. *ola*
17. Uzb. *ɔvla*

*Comments* Tkm. *aawla-* (Baskakov 1968: 18a) indicates pTk *a:b-.

#### #239 'straight' *tondo, *torto

**Mongolic** **tondV, **tundV

1. Dag. *tondo:xen*
2. Bao. *dʐaŋɕig*

*Comments* Not in Nugteren (2011). Bao. *dʐaŋɕig* is borrowed from Tibetan *drang.zhig* (Chén 1986: 206). As for Dag., specialists agree that *tondo* is a loanword from Man. (e.g. Kałużyński 1970: 138; Todaeva 1986: 168).

**Tungusic** **to?d?

1. Hez. *tondu*
2. Orc. *toŋno*
3. Udi. *toŋdo*
4. NanA. *toŋdo*
5. KU *toŋno*
6. NanB. *toŋgdo*
7. Ork. *toŋdo*
8. Ulc. *toŋdo*
9. Man. *tondo*
10. Xib. *tondoqun*
11. Sol. *tɔndɔ-hụŋ*

*Comments* SI2 Table 3.5 does not include any -*ŋ*-.

**Turkic** **tort, **tord

1. Tuv. *dort*

##### #249 '1PL pronoun' *bi-PL

*Comments* This is the same etymon as #14 *bi '1SG' with a plural suffix.

**Mongolic** **bi-da < **bi-ta

1. Kh. *bid*
2. Bur. *bide*, *bidener*
3. Kalm. *bidn*
4. Oir. *biden*, *bidnüːs*
5. Kmg. *bidə*
6. Dag. *bide*
7. Bao. *bədə*
8. Dgx. *bidʑiən*
9. Mog. *bidâ*, *bidâ-t*
10. MMoSH *bida*
11. MMoMuq. *bida*

*Comments* Nugteren (2011: 281) *bïda, *biden 'we (inclusive)'

**Tungusic** *bi-

1. Hez. *bəti*
2. Orc. *biti*
3. Udi. *minti*
4. Xib. *məs*
5. Evn. *mut*
6. Sol. *mit*
7. Evk. *miti*
8. StEvk. *mit*
9. SEChi. *mit*
10. NETut. *mut*
11. NETur. *mit*
12. Neg. *bittə* ~ *buttə*
13. Orq. *mitiː* ~ *mir*

**Turkic** *bi-z

1. BTat. *biz*
2. CC *bix*
3. NAlt. *pis*
4. OT *biz*
5. SAlt. *bis*
6. Aze. *biz*
7. Bas. *beδ*
8. Chu. *ebir*
9. CTat. *biz*
10. Dol. *bihigi*
11. Gag. *biz*
12. KKal. *biz*
13. KBal. *biz*
14. Kar. *biz*
15. Kaz. *bĭz*
16. Khk. *pəs*
17. Khl. *biz*
18. Kir. *biz*
19. Kum. *biz*
20. MChu. *pəs*
21. Nog. *biz*
22. Sal. *piser*
23. WYug. *mïs*, *mïster*
24. Sho. *pis*
25. KTat. *bez*
26. Tof. *bister*
27. Tks. *biz*
28. Tkm. *biz*
29. Tuv. *bis*
30. Uyg. *biz*
31. Uzb. *biz*
32. Yak. *bihigi*

##### #249 '1PL pronoun' *bu-PL

**Koreanic** **uli

1. HH *wuli*
2. SHG *wuli*
3. NHG *wuli*
4. SPA *wuli*
5. NPA *wuli*
6. JJ *wuli*
7. SJL *wuli*
8. NJL *wuli*
9. SGS *wuli*
10. NGS *wuli*
11. SCC *wuli*
12. NCC *wuli*
13. GG *wuli*
14. GW *wuli*
15. MK *wuli*

**Mongolic** **bu-, **bü-

1. ShYu. *budas*, *buda*
2. Mnh. *budasɛ*
3. Huz. *buda*, *budasge*, *bunaŋGʊla*

*Comments* Nugteren (2011: 281) *bïda, *biden 'we (inclusive)'.

**Tungusic** **bö-, **bu-

1. Hez. *bu ~ mun(u)*
2. Orc. *bu* (*mun-*)
3. Udi. *bu* (*mun-*)
4. NanA. *buə* (*bun- ~ bumbi-*)
5. KU *muː ~ mu* (*mun-*)
6. NanB. *buː* (*mun-*)
7. Ork. *buː* (*mumbeː- ~ mun-*)
8. Ulc. *buː ~ buə* (*mun-*)
9. Man. *bə*
10. Xib. *bo* (*mon-*)
11. Evn. *buː* (*mun-*)
12. Sol. *bu* (*mun-*)
13. Evk. *buː*
14. StEvk. *buː* (*mun-*)
15. SEChi. *buː*
16. NETut. *buː*
17. NETur. *buː*
18. Neg. *bu* (*mun-*)
19. Orq. *buː* (*mun-*)

##### #253 'yellow' *siara-

*Comments* This is the same root as #152 'white' *siara-. The Tungusic form should be excluded since it is marked as a borrowing.

**Mongolic** ?*siara

1. Kh. *šar*
2. Bur. *šara*
3. Kalm. *šar*
4. Oir. *šara*
5. Kmg. *šir*
6. Dag. *šara*
7. ShYu. *ʃəra*
8. Bao. *ɕira*
9. Dgx. *ʂɯra*
10. Mnh. *ʂira*, *ʂa*
11. Huz. *ɕira*
12. Kgj. *ʃira*
13. Mog. *šɪra*
14. MMoSH *šira*
15. MMoMuq. *šira*

*Comments* Nugteren (2011: 492) *sïra. pM *ia does not appear in SI2 Table 3.8.

**Tungusic** borrowing

1. Man. *sira* (bor.)

**Turkic** *sia:rï-

1. BTat. *sarï*
2. CC *sari*
3. NAlt. *sarï*
4. OT *sarïɣ*
5. SAlt. *sarï*
6. Aze. *sarï*
7. Bas. *harï*
8. CTat. *sarï*
9. Gag. *sarï*
10. KKal. *sarï*
11. KBal. *sarï*
12. Kar. *sarï*
13. Kaz. *sarï*
14. Khk. *sarïɣ*
15. Kir. *sarï*
16. MChu. *sa:rəɣ*
17. Nog. *sarï*
18. Sal. *sarï*
19. WYug. *sarïɣ*
20. Sho. *sarïɣ*
21. KTat. *sarï*
22. Tof. *sarïɣ*
23. Tks. *sarï*
24. Tkm. *sarï*
25. Tuv. *sarïɣ*
26. Uyg. *seriḳ*
27. Uzb. *sariḳ*
28. Yak. *araɣas*

#### Sources

In addition to the references cited above, we consulted the following sources:

**Japonic** Kokuritsu Kokugo Kenkyūjo (1963); Jōdaigo jiten henshū iinkai (1967); Osada & Suyama (1977–1980); Nakasone (1983); Hōsei Daigaku Okinawa Bunka Kenkyūjo (1987); Shōgaku Tosho (1989); Hirayama (1992–1994); Nihon Kokugo Daijiten Dai 2-han Henshū Iinkai, Shōgakkan Kokugo Jiten Henshūbu (NKD²); Ikema (2003); Miyagi (2003); Kiku & Takahashi

(2005); Fujimoto (2011); Tomihama (2013); Yonaguni Hōgen Jiten Henshū Iinkai (2019); Kajiku (2020); Pellard (2022)

**Koreanic** Nam (1997); Kungnip Kugŏwŏn (2016)

**Mongolic** Haenisch (1939); Cheremisov (1951); Weiers (1963); Bǎo (1984); Enhébātú (1984); Chén (1986); Todaeva (1986); Lǐ (1988); Bawden (1997); Saito (2008); Toadeva (2009); Mǎ & Chén (2012)

**Tungusic** Poppe (1931); Yamamoto (1969); Lǐ & Zhòng (1986); Hú (1994); Kim et al. (2008); Li & Whaley (2009); Kubo et al. (2011a,b); Zikmundová (2013)

**Turkic** Yudakhin (1965); Nadelyayev (1969); Baskakov (1968)

### 2 Evaluation of the agropastoral vocabulary

Robbeets et al. (2021) claim that the putative Transeurasian languages share a core of vocabulary related to millet cultivation, pig domestication, food preservation and textile production, which is taken to indicate that proto-Transeurasian can be dated back to the Early Neolithic, and that its speakers can be identified with early millet farmers in the West Liao River area. This hypothesis and the relevant evidence are detailed in "Supplementary Information 5: Inherited and borrowed correspondence sets for agropastoral vocabulary across the Transeurasian languages" (`20_Eurasia3angle_synthesis_SI 5_agropastoral_REV21.09.docx`), henceforth SI5, attached to Robbeets et al. (2021).

Within the framework of linguistic palaeontology (Blust 1988; Anthony 2007; Garnier et al. 2017), lexical comparisons can serve as evidence to reconstruct the material culture of the speakers of proto-languages and infer information about their homeland by comparison with archaeological evidence. However, specific criteria need to be met when making such inferences. First, the lexical comparisons need to be genuine cognates inherited from a common ancestor, and not borrowings, chance resemblances, or convergent evolution. Second, the meaning of the comparanda must denote a specific animal, plant, object, or activity related to a material culture, and this meaning needs to be identical across languages. Finally, the comparanda must be attested in enough languages to warrant reconstruction to their common ancestor.

Concerning the first aspect, in order to distinguish cognate words inherited from a putative common ancestor from borrowings and chance resemblances, Robbeets et al. (2021) set up several criteria (SI5: 1–2). The most important one is that of the regularity of sound correspondences, i.e. a cognate set including forms that do not correspond regularly with each other across languages cannot be taken as reliable evidence of inheritance. It is further specified that only the first (Consonant-)Vowel(-Consonant) portion of words should be examined for regular correspondences. In order to try replicating the analysis of Robbeets et al. (2021), we manually applied the same criteria as for sound correspondences in the core vocabulary (see Supplementary Information 1).[1]

Concerning the second aspect, Robbeets et al. (2021) specify that comparisons should involve words with comparable meanings and that these meanings should specifically concern agriculture or pastoralism (SI5: 1). We thus inspected all proposed comparisons and checked whether they involved a meaning specific to a Neolithic material culture across languages or not. For example, we excluded words meaning 'to spin' as a weaving activity when they had a more generic meaning not specific to any technological activity (e.g. 'to turn'). Comparisons involving words for nuts or dogs were also excluded because such words are also found in the lexicon of hunter-gatherer languages and are thus not specific to farming cultures. Such comparisons cannot be used to support the hypothesis that, if the Transeurasian languages are related, the speakers of the proto-Transeurasian language were Neolithic farmers.

An additional criterion is that a cognate set should be distributed in three or more language families (SI5: 1). All comparisons involving only two languages were thus discarded accordingly. We also rejected comparisons for which SI5 indicated a most recent common ancestor other than pTEA.

[1]Mention of tables and numbered sound correspondences in our detailed comments refer to those at the end of SI5.

In order to assess the Transeurasian West Liao River farming hypothesis, we thus evaluated each comparison listed in SI5 for the above criteria. Detailed comments are provided below, and we summarized our evaluation within a table (`Supplement_A-linguistics/agropastoral-vocabulary.tsv`). For each comparison proposed in SI5, we encoded:

1. whether the comparison was phonologically regular across all proto-languages for a which a cognate is listed (regularity criterion);
2. whether the meaning of the compared words were semantically compatible (comparability criteria);
3. whether the compared words belong specifically to the agropastoral lexicon in all languages for a which a cognate is listed (specificity criterion);
4. whether the comparison involved three or more language branches (distribution criterion);
5. whether the comparison allowed a reconstruction at the pTEA ancestor node in the phylogenetic tree (reconstructibility criterion).

Our results are as follows: out of the 43 comparisons presented, 31 etymologies follow the sound correspondences, only 22 have comparable semantics, only 9 really belong to the realm of agro-pastoral vocabulary, only 27 appear in more than three branches, and only 29 are reconstructible at the pTEA level. None of the comparisons satisfies all five criteria. The identification of Proto-Transeurasian speakers with early millet farmers of the West Liao River area is thus not supported by empirical evidence.

#### 2.1 Cultivation

##### (1) pTEA *pata 'field for cultivation'

*Comments* The pK form is irregular since roots with two vowels *a in pTEA should appear with two *ʌ in Koreanic (#32b.), i.e. we expect pK *pʌtʌ- and not *patʌ-. The agricultural meaning of the Turkic forms is debatable.

**Turkic** pTk *(p)ata 'delimited field irrigated for cultivation'
*Comments* The reconstruction *(p)ata actually conflates two different cognates sets, meaning 'island' and 'field', respectively. Alternative Turkic-internal etymologies are possible, analyzing e.g. OT *atïz* as derived via the collective suffix *-z* from the verb *at-* 'to throw, to remove' or 'to separate, to divide', since it basically means to set up portions of sowable land between two irrigation ditches (Nadelyayev 1969: 67). Also, the semantics of some attested forms go against the idea of fertile and cultivated land and are incompasible with agriculture, e.g. Kir. *adïr*, which means specifically 'hilly terrain *not suitable for irrigation*' (Yudakhin 1965: 23).

##### (2) pTEA *muda 'uncultivated field'

*Comments* None of the forms involved has the meaning 'field' and they do not belong to the agricultural lexicon. The sound correspondences are anyway irregular since the second consonant pTg *-d- should correspond to pK *-l- (#10) and not *-t- (#8). Also, the second vowel pJ *a and pTg *a should correspond to pK *a (#32) and not *ʌ (#39).

**Japonic** pJ *muta 'uncultivated land, marshland'
*Comments* The Japonic comparanda are problematic since there is confusion between two different roots, neither of which has the primary meaning of 'uncultivated field'. First, the Okinawan and Southern Ryukyuan forms cited all mean 'earth, soil', and not 'swamp, marshland', even less 'field'. They all go back to pR *mita (B) 'earth, soil',[2] which is cognate with Hac. *midʑa*, and not *muta (A) 'swamp, marshland',[3] which is a different etymon. SI5 gives Shodon *mutha* (*mùtʰâ:*) 'swamp' as a cognate, but Shodon also has *mítɕʰă:* 'earth'. Both roots are found in Yor. (*nìtɕá* 'red clay' vs. *mùtà* 'black ditch mud') and Kikai (*mitɕa* 'earth, soil' vs. *muta* 'black, sticky and watery earth') too. In Japanese dialects, there is indeed a form *muta* which usually means 'damp ground, marsh, ill-drained field', but attestations are restricted to Kyushu and one place in the neighbouring Yamaguchi prefecture.[4] There is also one isolated attestation of *muta* 'fallow paddy field' on Sado Island in Northern Japan. Japanese dictionaries usually consider *muta* to be related to *nuta* 2.1 'marsh, swamp, ill-drained field, mud', which is widely attested throughout Japan since the 12th c. Some dialects have a form *nita* which could be the result of contamination or something else. In any case, the original meaning of *muta* seems to be 'mud, muddy place', and it is not an agricultural word.

**Koreanic** pK *mutʌ 'dry land' + *-(ɨ/ʌ)k place suffix
*Comments* MK *mwùth* indeed means 'land, dry land, terra firma', and does not denote a field nor anything related to agriculture.

**Tungusic** pTg *muda 'plain, open field, highland'
*Comments* The Tungusic forms do not denote a field nor anything related to agriculture.

##### (3) pTEA *iuse- 'to plant, grow (plants)'

The Koreanic form are not semantically comparable with the other forms, which are not anyway specific to agriculture but generic verbs meaning 'to grow, to multiply, to increase'.

**Koreanic** pK *yes- 'to grow grain' + *-(ʌ/ɨ)k > *isak, *-(ʌ/ɨ)k 'edible plant suffix'
*Comments* The relationship proposed between MK *yés* 'candy' and *ìsàk* 'ear of grain' is speculative both phonologically and semantically. No verb 'to grow grain' can be reconstructed.

##### (4) pTEA *urə- 'to grow, ripen (of plants)'

*Comments* The Japonic form should be excluded from the comparison since it cannot be securely reconstructed in pJ.

**Japonic** pJ *ura- 'to mature, ripen (of plants)' + *-(C)i- causative-anticausative
*Comments* OJ "*ure-* 'to mature, ripen (of plants)'" does not exist, as it is first attested in 1603 only. Hatoma "*ʔurin* 'to ripen'" does not exist either, and the attested form is *ʔú:mùŋ*, which is cognate with EMJ *um-* 'fester, get ripe' and not with Japanese *ure-*. The late attestation of Japanese *ure* and the absence of cognates in Ryukyuan indicate it is a recent development, which forbids its comparison with other languages.

[2] Though forms such as Yng. *ǹtà* could theoretically go back to either *mita or *muta, tonal correspondences indicate they are cognate with *mita* (B).
[3] Attested in Amami Ryukyuan only.
[4] The 1275 etymological dictionary *Myōgoki* lists a dialectal word *muta* 'grassy marsh' but does not indicate its provenance.

##### (5) pTEA *pisi- 'sprinkle with the hands, sow'

*Comments* None of the forms compared has a meaning specific to agriculture but just mean 'to sprinkle, to scatter'. As noted in SI5, only the Koreanic forms also have the additional meaning of 'to sow'.

**Koreanic** pK *pis- 'to sprinkle, scatter, sow'
*Comments* The rationale behind the pK reconstruction *pis- is unclear. The verb 'to sprinkle' MK *spih-* probably contains the intensive prefix *s-*, the noun *pí* < *piWi* 'rain', and the verbalizer *ho-* 'to do' (Martin 1996: 47). Korean *ppuli* 'root' is surely unrelated since it comes from MK *pwùlhúy*, which reconstructs as pK *pulɨkɨy (Vovin 2006).

##### (6) pTEA *pisi-i (sow-INS.NMLZ) 'seed, seedling', *-i/ø instrumental deverbal noun suffix

*Comments* The Mongolic forms are not semantically comparable with those from other families and they do not belong to the agricultural lexicon anyway.

**Koreanic** pK *pisi 'seed; lineage'
*Comments* MK *psí* reconstructs as pK *pʌsi or *pɨsi, but not *pisi, whose only purpose is to make the comparison with other languages look regular.

**Mongolic** pM *pesi ~ *pisi 'origin or base of a plant'
*Comments* The Mongolic forms cited are generic words meaning 'basis, origin, stalk, trunk, handle' and have nothing to do with agriculcure, less millet and seeds.

**Tungusic** pTg *pisi-ke 'broomcorn millet (*Panicum miliaceum*)'
*Comments* Vovin (2006: 260) has suggested that Man. *fisike* 'millet' is a borrowing from Korean that then spread within Tungusic, but this hypothesis is not discussed in SI5. While Man. *fisihe* corresponds to Chinese *shŭ* (黍) 'broomcorn millet' in some translated texts, according to Hú (1994), it means *Setaria italica*, and it is *ira* that means 'broomcorn millet' instead.

##### (7) pTEA *kipi ~ kipe 'components that are removed from the grain harvest, barnyard grass (*Echinochloa crusgali*)'

*Comments* The vowels in the first syllable do not follow the given sound correspondences. pJ *i can correspond to pTg *i but incorrectly predicts pK *i (see below) and pTk *i or *ï (#40), while pK *ɨ or *ʌ should correspond to pJ *u, pTg *u and pTk *u, *ü or *ï (#38, 39), and pTk *e entails either pJ *a or *ə, pK *e, and pTg *e (#33, 34). The second consonant is irregular in Japonic, as *-p- is expected (#2) and not *-m- or *-Np- (#5, 6, 26).

**Japonic** pJ *kinpi ~ *kimi ~ *kipi-mi 'millets for human consumption such as barnyard millet (*Echinochloa esculenta*), broomcorn millet (*Panicum miliaceum*)'
*Comments* The hypothesis of "a morpheme boundary in Proto-Japonic, involving pJ *-mi, deriving from OJ *mi*$_2$, J *mi* 'fruit, nut, kernel'" implies that the root is just *ki, and that it cannot be compared to the forms from other languages. However, OJ *ki*$_1$*mi*$_1$ 'broomcorn millet' and *mi*$_2$ 'fruit' have different vowels, and *mi*$_2$ cannot go back to pJ *mi, which invalidates that hypothesis.

**Koreanic** pK *kipi > *phi 'barnyard millet (*Echinochloa esculenta*)'
MK *phí* 'barnyard millet' can go back to several proto-forms: *kɨpi, *kʌpi, *pɨki, or *pʌki, but not *kipi. Reconstructing *i in the first syllable has no basis and only serves to make the comparison with other languages look regular.

**Turkic** pTk *kẹpek 'bran (from millet, barley), chaff'
*Comments* The vowel *ẹ in pTk *kẹpek does not appear anywhere in the correspondence tables. This is probably due to the fact that it was copied from Starostin et al.'s (2003) database, while Robbeets et al. (2021) use a different reconstruction system. According to SI5 Table 5.10, the reconstruction should be *kepek instead.

##### (8) pTEA *amʊ 'cooked cereal, millet gruel'

*Comments* The Sino-Tibetan root meaning 'eat' and 'rice' is never used to denote broomcorn or foxtail millet. If, as assumed in SI5, Mongolic *amu-n was borrowed from this etymon, this would be contradictory with reconstructing the meaning 'millet' back to pTEA. The meaning 'broomcorn millet' is anyway only attested in Mongolian, and even if an etymological relationship between these etyma were possible, it would not be evidence for domestication of millet in pTEA. The second vowel is irregular in Japonic, since we would expect pJ *u (#39) and not *a(i).

**Japonic** pJ *amai 'cereal starch'
*Comments* In Ryukyuan, the meaning is always 'candy' as in Modern Japanese, which suggests it might be a loanword from Japanese. The word is anyway a transparent derivation from the adjective *ama-* 'sweet', as *ame*$_2$ is the result of the saccharification of starch. This word thus does not belong to the agricultural lexicon.

##### (9) pTEA *sarpa 'spade'

*Comments* The pTEA sequence *CaCa should yield pK *ʌ and not *a in the first syllable (#33). The proposed Sino-Tibetan *tsrop 'spade' is not based on known phonetic correspondences and is a pseudo-reconstruction (Fellner & Hill 2019). Aside from the fact that the meanings do not match, proto-Kuki-Chin *-op remains *-op* in Thadou and Tiddim (VanBik 2009), hence these words cannot be cognate with the Chinese etymon, which is isolated in the Sino-Tibetan family. Thus, no such etymon can be reconstructed back to the proto-Sino-Tibetan level.

##### (11) Proto-Mongolo-Tungusic *pure 'seed, sprout, offspring'

*Comments* The forms compared are generic ones for 'offspring, descendant' and do not belong to the agricultural lexicon.

##### (12) Proto-Japano-Koreanic *non 'field for agriculture'

*Comments* The semantics of the Japonic form ('uncultivated plain') invalivate the comparison with both pK *non 'irrigated field, rice paddy field' and Sinitic *nowng* < *nˤuŋ (農) 'agriculture'. Phonologically, the absence of a final nasal would remain unaccounted for in Japonic.

**Japonic** pJ *no 'field'

*Comments* The Japonic forms do not denote a field for agriculture but an uncultivated plain covered with vegetation, though in a few Japanese dialects it has acquired the meaning 'field'. This primary meaning is preserved in compounds with names of animals and plants, where *no-* adds the meaning 'wild', as opposed to 'domestic, cultivated'. In addition, the Okinawan *mo:*-like forms are irregular and probably represent a different etymon.

###### (13) Proto-Japano-Koreanic *mati 'delimited plot for cultivation'

*Comments* Since the second vowel in pK can only be *ʌ, the correspondence with pJ *i is irregular (#39, 40).

**Japonic** pJ *mati 'delimited plot for cultivation'
Though *mati* in both OJ and modern Japanese dialects often mean 'plot of a paddy field', it has other meanings too, especially in Ryukyuan, such as 'market, place where people gather, quarter of a town'. It can also mean 'lines carved on bones or turtle shells for divination', 'grade, class', 'boundary notch between the tang and the blade of a sword', and the reduplicated form *mati-mati* means 'delimited, diverse'. A plausible etymology would be *ma* 'space, interval' + *ti* 'path', hence the general meaning 'delimited area, section, plot', in which case there is no relationship with agriculture.

**Koreanic** pK *mat(i)-k 'delimited plot for cultivation'
*Comments* MK *màth* can only reconstruct as pK *matʌk or *makʌt, not *matik.

#### 2.2 Food production and preservation

###### (1) pTEA *saga- 'to ferment'

*Comments* The Mongolic and Turkic forms clearly mean 'to milk' and have nothing to do not with fermentation. The comparison with Koreanic 'to ferment, to rot' is thus problematic, as is the inclusion of Japonic. According to sound correspondence #32b., the expected pK form is *sʌkʌ- and not *sak-

**Japonic** pJ *saka- 'to ferment, be in heat, bloom'
*Comments* The relationship proposed in Japonic between 'rice wine', 'to be at peak, to be in rut', and 'to bloom (flowers)' is unclear and semantically not convincing. There is not related verb 'to ferment' attested.

**Mongolic** pM *saga- 'to ferment; to reduce (food); to milk'
*Comments* The meaning 'to ferment' is not attested for Written Mongolian *saɣa-*, nor in other Mongolic languages.

###### (2) pTEA *sugu- 'to ferment' (in vowel alternation to (1))

*Comments* The vowels do not correspond since pM *u and pTk *u only correspond to pK *ʌ and not *u (#39). The comparison is also semantically unjustified since verbs related to dairy production are compared with a noun for rice wine.

**Koreanic** pK *su(k)u- 'to make alcohol'

*Comments* The MK form *swùùl* is earlier attested in the *Kyerim yusa* (12th c.) with the Chinese transcription *su bwot* 酥孛, which shows that it originally contained a labial consonant and not a velar. Moreover, this etymon is a noun meaning 'rice wine', and reconstructing a verb 'making alcohol' by cutting off the final consonant is not justified.

##### (3) pTEA *sü:- 'to reduce/preserve food by fermentation'

*Comments* The vowel *ü reconstructed for pTEA does not appear in SI5 Table 5.12 and no correspondence is given for the suggested pM vowel *iü present in one of the several forms given. Moreover, only the Japonic and Koreanic forms are related to fermentation.

**Mongolic** pM *sü- ~ *siü- 'strain, filter, skim off, reduce (food)'
*Comments* None of the words listed is related to fermentation or more generally to food preservation.

**Turkic** pTk *sü:- to filter, strain (milk or soup), reduce (food)
*Comments* None of the words listed is related to fermentation or more generally to food preservation.

##### (4) pTEA *bilče- 'to ferment a liquid, mix with a liquid'

*Comments* The Japonic form poses several problems and should better be left out of the comparison, and the Mongolic form has nothing to do with fermentation and thus cannot be compared with the Koreanic form.

**Japonic** pJ *pisi 'fermented liquid'
*Comments* Japanese *pisipo* is not phonetically attested in OJ but in EMJ only. It is obviously formed on the root *sipo* 'salt', and the initial *pi-* could be 'dry, drain' since it is a liquid extracted from fermented barley, soy beans, and salt. Japanese *hisiko* 'Japanese anchovy' is not attested in OJ but only in the 15th c. onwards. Though its etymology is unsure, it is not plausible that such a recent word would preserve an otherwise unattested pJ root *pisi. There is thus no evidence for a root *pisi 'fermented liquid' in Japonic.

**Koreanic** pK *pici- 'ferment (liquid), brew, make dough'
*Comments* There is no evidence for reconstructing a final *i in pK for this root, as MK *pìc-* is a Class 1 verb stem and thus simply goes back to pK *pic-. Depending on one's theory of lenition and root structure in Korean, the reconstruction could also be *picɨ- or *picʌ-, but the final vowel cannot be *i.

**Mongolic** pM *bilca- 'mix a liquid with food'
*Comments* The meaning of this etymon cannot be reconstructed as 'mix a liquid with food' since all attestations mean 'to smear, to splash, to pound' or 'to become flat, thin'. This is not an agricultural word.

##### (5) pTEA *silɔ 'broth, soup, juice; liquid food extracted from vegetables, fruit or meat'

*Comments* The second vowel is irregular: pJ *u predicts pM *u (#38) or *ü (#39), and on the other hand pM *ö predicts pJ *ɨ (#37) and not *u.

##### (6) pTEA *niku- 'to crush to pulp'

*Comments* None of the forms compared are specific to agriculture but are generic verbs meaning 'to crush'.

##### (7) pTEA *suru- 'to grind, rub'

*Comments* None of the forms compared are specific to agriculture but are generic verbs meaning 'to rub'.

##### (8) pTEA *simi- 'to soak (food)'

*Comments* Cognates are proposed only for Japonic and Mongolic, but the forms do not denote anything related to food production or preservation.

**Japonic** pJ *sima- 'to soak, permeate'
*Comments* The Japonic forms have nothing to do with food production or preservation. They mean 'to get damp', 'to permeate'.

**Mongolic** pM *sime- 'to soak (food), moisten, suck up'
*Comments* The meaning of the root is clearly 'to suck, to sip', with the intransitive derived verb meaning 'to become soaked'. It has nothing to do with food production or preservation.

##### (9) pTEA *ulu- 'to soak, wet'

*Comments* Since the pJ form reconstructs as *oro and not *uru, the vowels are irregular since they do not follow correspondences #45 and #38. Moreover, the forms have nothing to do with food production or preservation.

**Japonic** pJ *uru- 'to irrigate, make wet' + *-pa- -pə- reflexive-anticausative
*Comments* Many of the Japanese forms cited are not attested in OJ proper, and the Ryukyuan forms are not verbs meaning 'to irrigate' but nouns meaning 'wetness (especially of a field)'. The Miyako Ryukyuan forms indicate that the pJ root should be reconstructed as *oro- rather than *uru-, otherwise we would expect a †*vv*- in Miyako (cf. pJ *ur-i 'to sell' > Miyako *vv*).

##### (10) pTEA *nɔr- 'to soak'

*Comments* pJ *u does not regularly correspond to pM *o (#35, 36, 38, 39). Moreover, the forms have nothing to do with food production or preservation.

**Japonic** pJ *nura- 'to be(come) wet''
*Comments* The vowel syncope in the first syllable in Shu. indicates that it comes indeed from pJ *u and not *o.

##### (11) proto-Altaic *deb- 'to soak'

*Comments* None of the forms compared are specific to agriculture but are generic verbs meaning 'to make wet' or 'to become wet'.

###### (12) Proto-Japono-Koreanic *kama- 'to soak, brew'

*Comments* The Japonic and Koreanic forms have incompatible semantics ('to bite' vs. 'to bathe'), and they anyway do not belong to the agricultural lexicon.

**Japonic** pJ *kamə- 'to chew, brew, dye'
*Comments* The primary meaning of Japanese *kam-* is 'to bite'. The meaning 'to brew' is derived from the fact that sake was made by chewing and spitting out rice, thereby using the amylase contained in saliva to break down the starch into sugar. This is not an agricultural word.

**Koreanic** pK *kʌmʌ- 'to put in a bath'
*Comments* Since MK *kóm-* is a Class 2a verb stem, the reconstruction should rather be simply *kʌm-.

#### 2.3 Durable wild food resources

None of these words can be taken as evidence of an agropastoral culture.

###### (1) pTEA *kuru 'edible nut used for starch production such as walnut, acorn, chestnut or pine nut'

*Comments* The Koreanic form denotes a tree and not a nut, so that it cannot be compared with the Japonic and Tungusic forms.

###### (2) pTEA *xʊsi 'edible nut used for starch production such as walnut, acorn, chestnut or pine nut'

*Comments* The Japonic form is problematic and the Mongolic one does not denote a nut but a tree. If valid, this comparison thus rather involves a kind of tree rather than a nut or acorn, and it is thus not directly related to food resource, less to agriculture.

**Japonic** pJ *kusi 'chestnut'
*Comments* OJ *kusi* is a hapax legomenon, probably a dialectal word cognate with *kuri* in (1).

**Mongolic** pM *kusi 'walnut'
*Comments* This root denotes a tree rather than its fruit, as is clear from the meaning of its reflexes in the different languages.

###### (3) pTEA *abu 'plant of the Althaea genus with roots rich of starch'

*Comments* The final *-k of pK is left unexplained, as is the final syllable in OJ. Since the Japonic form does not denote a nut, it is not comparable with the Koreanic form.

**Japonic** pJ *apu 'hollyhock (*Althaea rosa*)'
*Comments* NKD[2] identifies OJ *apupi*$_1$ with *Malva verticillata* instead. Segmenting and leaving out the final *-pi*$_1$ is not justified.

**Koreanic** pK *apok 'marshmallow (*Althaea officinalis*)'

*Comments* The pK reconstruction is not straightforward due to conflicting data. Though some varieties exhibit a *-p-*, others do not and have zero, *-h-*, or *-k-* instead.

**Mongolic** pM *abu 'marshmallow (Althaea officinalis)'
*Comments* There are no cognates in other Mongolic languages than Kh., which is suspect.

###### (4) Proto-Japano-Koreanic *pami 'edible true nut, i.e. dry fruit with only one seed such as acorn, hazelnut and chestnut'

*Comments* The Japonic form does not denote a nut, and the second vowel does not correspond regularly to the Koreanic one (#40, 39) once the pK form is reconstructed according to the known historical phonology.

**Japonic** pJ *pami 'edible true nut such as hazelnut or acorn'
*Comments* NKD[2] identifies EMJ *pami* (not attested in OJ) as *Lamium barbatum* rather than *Phlomis umbrosa*, but neither grows nuts. EMJ *katapami* (not attested in OJ) is *Oxalis corniculata*, a low-growing herbaceous plant which does not produce nuts either. On the other hand, OJ *turupami*$_1$ is *Quercus acutissima* and by extension the name of its fruit, the acorn, and of the pigment made from it, and EMJ *pasipami* is *Corylus heterophylla*, the Asian hazel. Reconstructing pJ **pami* 'edible nut' is not well motivated since it is found in many compounds for names of plants that do not produce nuts.

**Koreanic** pK *pami 'edible true nut such as chestnut or hazelnut'
*Comments* MK *păm* reconstructs as pK *pamʌ and not *pami. The umlauted form exhibited by some dialects are probably due to the influence of the nominative suffix *-i*, which is likely since non-umlauted forms are listed too.

#### 2.4 Domesticated animals

##### (1) pTEA *inu ~ *ina 'dog'

*Comments* Since the domestication of the dog precedes agriculture by several millennia (Liu & Chen 2012: 96–97), the etyma for 'dog', even if they are really related, are not relevant to this question. The proposed Tungusic reconstruction is marred with problems that prevent the evaluation of this comparison. The sound correspondences are thus not regular.

**Japonic** pJ *inu ~ *ina 'dog'
*Comments* The reconstruction pJ *ina rather than *inu is only supported by the unexpected *ina* form in Tarama, which is probably due to a diminutive suffix *-a* common in Ryukyuan for some animal names. All other Ryukyuan forms point to *inu, or perhaps *enu.

**Tungusic** pTg *ina ~ *inu 'dog'
*Comments* No cogent explanation is given for the presence of an initial nasal in the Tungusic cognates. In such cases, initial *ŋ- is usually reconstructed, but there is no such consonant in the correspondence tables. One correspondence for *g- in SI5 Table 5.5 lists initial *ŋ-* as a reflex in some languages, but it wrongly predicts an initial *g-* in Jurchen, Manchu, and Sibe, and it also lists either *g-* or *ŋ-* as

possible reflexes in other languages, but without any conditioning. Moreover, there is no basis for reconstructing the second vowel as *u as an alternative instead of simply *a.

###### (2) Proto-Altaic *toru 'young male pig'

*Comments* The forms are not semantically comparable since the Turkic forms do not denote an animal of the Suidae family but cattle, camels, or horses.

###### (3) Proto-Mongolo-Tungusic *uli 'pig'

*Comments* Within Mongolic, this form only appears in Khitan, which does not appear in the correspondence tables. The regularity of the sound correspondences has yet to be demonstrated.

#### 2.5 Textile production

Since needles are attested since the Palaeolithic (Wang et al. 2020), terms for 'needle' or 'sewing' are not diagnostic of an agricultural society.

###### (1) pTEA *nap- 'to make rope'

*Comments* The proposed Koreanic reconstruction poses too many problem and the sound correspondence with other families is not demonstrably regular. The semantics of Japonic 'to twist, to make rope' and Tungusic 'tiers' are not straightforwardly comparable.

**Koreanic** pK *nap- 'twist, spin'
*Comments* The relationship between MK *nàh-* 'to spin, to make yarn' and Modern Korean *kkunapul* 'piece of string' was first suggested by Martin (1966: 240), who however noted the difficulty in this derivation. The *napi*-like forms meaning 'to quilt' found in modern varieties are probably cognate with MK *nwùpí-* 'to quilt' instead. The reconstruction of a *p is speculative and fails to explain the presence of a *h* in the verb 'to make yarn'.

**Tungusic** pTg *nap- 'to make rope' + *-ki resultative nominalizer
*Comments* The Tungusic forms do not mean 'rope' or 'to make rope' and have nothing to do with textile production or twisting but mean 'tiers, straps'.

###### (2) pTEA *nup- 'to sew'

*Comments* The Tungusic form is not semantically comparable with the other forms since it means 'to pierce, to prick' and not 'to sew', and it is not specific to textile.

###### (3) pTEA *sili- 'to sew'

**Tungusic** pTg *sira- 'to sew together, tie together'
*Comments* The Tungusic forms are generic verbs meaning 'to connect, to lengthen, to tie' and the reconstruction of a verb specifically meaning 'to sew' is not justified.

##### (4) pTEA *pɔrɔ- 'to weave (cloth)'

*Comments* As noted in SI5, the Korean and Japonic forms are irregular since they fail to exhibit the initial *p- expected from correspondence #1. Moreover, the correspondence of the vowel is irregular too, since pTk has *ö instead of the expected *o (#35). The Tungusic and Mongolic are anyway generic verbs that do not mean 'to weave' but 'to spin'.

**Japonic** *orə- 'to weave'
*Comments* The pJ reconstruction *orə is phonotactically impossible. Arisaka's First Law entails that it should be *or(o)- or *ər(ə)-, and using external evidence to choose which vowel to reconstruct is begging the question.

**Tungusic** pTg *poro- 'to spin (nettle and hemp threads); to rotate, turn'
*Comments* This is a generic word meaning 'to spin, to turn'.

**Mongolic** *poro- 'to tie around, entwine; rotate, turn'
*Comments* Nugteren (2011: 360) reconstructs *horïa- 'to bind, wind, spin, wrap', with *h < *p. This root does not mean 'to weave' and is not directly related to textile production.

**Turkic** pTk *pö:r- 'to plait, weave'
*Comments* Initial *p- and its reflexes in the daughter languages are missing from SI5 Table 5.9.

##### (5) pTEA *tɔmʊ- 'to spin'

*Comments* pM *-o- and pTg *-o- do not correspond to pJ *-u- (corr. #35–36).

**Japonic** pJ *tumu 'spindle'
*Comments* Neither the noun *tumu* 'spindle' nor the verb *tumug*- are attested in OJ. Most of the Ryukyuan forms quoted are reflexes of a different etymon. In any case, it is a generic term for 'to spin' probably related to *tumu-zi* 'whorl of hair, whirlwind' and *tumu* ~ *tubu* 'spiral shellfish'.

##### (6) pTEA *giri- 'to cut (cloth)'

*Comments* Most of the forms quoted are not specialized forms related to textiles but just generic terms for cutting.

##### (7) Proto-Japano-Koreanic *parʌ- 'to sew'

*Comments* The correspondence of pJ *r with pK *n is irregular (#28, 29, 30, 31). Moreover the pJ reconstruction *paru- indicates that it is a case of corr. #39b., from which we should expect pK *pʌlʌ- or *pɨlɨ-, which makes the vowels irregular too.

**Japonic** pJ *paru-i 'needle' < *paru- 'to sew' + *-i deverbal noun suffix
*Comments* There is no verb 'sew' *paru reconstructible in Japonic, only the noun 'needle'. Nothing indicates that it should be reconstructed as *parui instead of *paroi or *pari. Since it is not an apophonic noun, it should be better reconstructed as *pari.

**Koreanic** pK *palʌ-l < *palʌ- ‘to sew’ + *-l deverbal noun suffix
*Comments* As noted by Vovin (2010: 96), LMK *palol* ‘needle’ “is in fact an Early Modern Korean *hapax legomenon* attested in the *Ma kyeng enhay*, dating from the Inco period (1623-49)”. All other Korean forms have an *n* and not an *l*.

##### (8) Proto-Japano-Koreanic *paca- ‘to weave (cloth) with a loom’

*Comments* Japonic ‘loom’ and Koreanic ‘to piece together’ are not semantically comparable.

**Koreanic** pK *pʌcʌ- > *pcʌ- ‘to weave’
*Comments* The MK form is *pcó-* and not *pcá-*. This is a generic verb meaning ‘to piece together, to assemble, to organize, to tie, to knit’ that is not specific to textile production.

##### (9) Proto-Japano-Koreanic *pu- ‘to spin, twist (thread)’

*Comments* Japonic ‘knot, stitch’ and Koreanic ‘twisted’ or ‘shuttle’ are not semantically comparable.

**Japonic** pJ *pu- ‘to spin, twist (thread)’
*Comments* The reconstruction of this etymon as a verb meaning ‘to spin, to twist’ is problematic, since the only attested forms are nouns meaning ‘node, knot of a plant’ or ‘stitch, knot of a mat or a woven fence’.

**Koreanic** pK *pu- ‘to twist (thread)’
*Comments* The MK form of ‘twisted’ is *pwuy-* and not *puy-*. The relationship with *pwùk* ‘shuttle’ via a nominalizing suffix is at best speculative, and the low tone in *pwùk* indicates it may be reconstructed as *pukɨ.

##### (10) Proto-Japano-Koreanic *ʌsa ‘hemp’

*Comments* No correspondence for pK initial *ʌ is given in SI5 Table 5.12, and from corr. #41 we would instead expect *a in pK. This comparison is thus irregular, which suggests that the alternative explanation suggested in SI5 that it is a *Wanderwort* is the correct one.

**Japonic** pJ *asa ‘hemp’
*Comments* The Kohama form *basa* does not come from a compound **bu: + asa* but is simply the Sino-Japanese (芭蕉) name of the Japanese fiber banana, a word unrelated to *asa ‘hemp’. The Japanese fiber banana is used to produce textile, hence perhaps the confusion, and a form *basa* is well attested in Southern Ryukyuan, and also in Northern Ryukyuan.

##### (11) Proto-Japano-Koreanic *mosi ‘ramie (cloth)’

*Comments* As noted in SI5, this word is usually thought to be a borrowing.

**Japonic** pJ *mosi ‘ramie (cloth)’

*Comments* The pR form for 'straw mat' cannot be *musi-su, otherwise palatalization should be blocked in Northern Ryukyuan and the final vowel would not be *u* is Southern Ryukyuan. It should be reconstructed as simply *musiro (or possibly *mosiro). In any case, the relationship between 'ramie cloth' and 'straw mat' is not straightforward since mats were made with stalks of bamboo, rush, or straw, but never with ramie fiber. It these words are unrelated, then *musi is not attested in Ryukyuan, which makes it likely that it is a loanword.

**Koreanic** pK *mosi 'ramie cloth'
*Comments* MK *mwòsì* has a double low tone with two light syllables and none of the two minimal vowels *o* and *u*, which is unusual. Since other such nouns are often loanwords (Ramsey 1991: 220–221), this might be the case of this word too.

#### Sources

In addition to the references cited above, we consulted the following sources:

**Japonic** Iwakura (1941); Jōdaigo jiten henshū iinkai (1967); Martin (1970); Shōgaku Tosho (1989); Hirayama (1992–1994); Nihon Kokugo Daijiten Dai 2-han Henshū Iinkai, Shōgakkan Kokugo Jiten Henshūbu (NKD[2]); Kiku & Takahashi (2005); Yonaguni Hōgen Jiten Henshū Iinkai (2019); Pellard (2022)

**Koreanic** Nam (1997); Lee & Ramsey (2011); Kungnip Kugŏwŏn (2016)

**Mongolic** Nugteren (2011)

**Turkic** Yudakhin (1965); Nadelyayev (1969)

### 3 Evaluation of the archaeological evidence

Robbeets et al. (2021) collected 171 archaeological features[1] for 255 Neolithic and Bronze Age sites in northern China, the Primorye, Korea and Japan, and coded those as present (1) or absent (0). They produced a phylogenetic analysis of these data using BEAST.

We attempted to replicate the results of the phylogenetic analysis of archaeological sites proposed by Robbeets et al., based on the supplementary file made available by the authors. Our analyses are all based on the XML made available by the authors at `https://github.com/rbouckaert/Eurasia3angle/blob/v1.0/41_Eurasia3angle_synthesis_SI%2021_XML%20files_cultures/pdcov-ucln-bbp.xml` and on their Excel data file.

We note some discrepancies between the XML file and the description of the analysis in the text of the Supplementary Information 8:

- the XML file includes 259 taxa, a discrepancy with the published paper which mentions 255 taxa; we removed four taxa (Shuiquan_Zhaobaogou_F1, Shuiquan_Zhaobaogou_F8, Daiwangshan2, Tongsamdong), as they seem to be left over from a previous version of the file; this leaves the expected 255 taxa;
- the text of the Supplementary Information 8 mentions two topological constraints which are imposed a priori, but these constraints are not present in the XML; our main analysis does not include these constraints, but we also ran an analysis with the prior constraints and found this had only a minor effect on the results;
- a small number of ages are different between the XML and the Excel data file; we rely on the ages provided in the Excel file.

All of our analyses are based on data for 255 taxa, using the same model as in the original study. The data are the same as in the original study, except that we correctly mark some data as missing, as mentioned in Section 3.1. We ran five analyses:

1. our main analysis includes all the data and no prior constraint;
2. one analysis with all the data and the two topological prior constraints listed in Section 3.2.1;
3. three analyses on subsets of the data, as detailed in Section 3.3. .

We wish to answer two questions:

1. is a tree model appropriate for these data?
2. if we accept the tree model, do the inferred phylogenies support the paper's conclusions? .

We answer both questions in the negative.

[1]Though Robbeets et al. state that they collected 172 features, their data only contains 171 features.

#### 3.1 Issues with missing data

An inspection of the data used shows that Robbeets et al. (2021) have treated missing data as zero values. We systematically checked the raw data for the Chinese archaeological sites reported in Robbeets et al. (109 out of 255 sites) and found 2,184 missing values that were improperly coded as zero values. For all sites where no burials were found, burial features were incorrectly coded as 0. This problem was also present in the sites with missing flora and fauna records. The corrected data are made available in the file `archeology_missingcorrected.csv`.

- grouping the Xinglongwa (8200–7400 BP), Zhaobaogou (7400–6500 BP) and Hongshang (6500–4900 BP) cultures from the West Liao together (sites #1–5, 10–13, 15, 18, 22, 23, 25, 34–38, 87, 88);

#### 3.2 Reconstructed topology

We reproduced the analysis using the complete corrected data set. The resulting consensus tree is shown in Figure 3.1. This consensus tree is highly multifurcating, which makes evident the very high uncertainty in the tree topology: only a small number of clades have support above 50%. This aspect of the posterior was not evident in the original study, as Robbeets et al. only produced a Maximum Clade Credibility (MCC) tree. A MCC tree is necessarily binary: using a MCC tree to summarize a distribution of trees with very high variance masks this variance. In this case, the consensus tree shows the variance better, and makes evident that there is very little tree-like signal in the data.

##### 3.2.1 Posterior support for key clades of interest

###### Prior clades

The following clades were proposed by Robbeets et al. Their Supplementary Information 8 lists these clades as being enforced by the prior, although we could not find these constraints in the XML file provided:

- grouping the Xinglongwa (8200–7400 BP), Zhaobaogou (7400–6500 BP) and Hongshang (6500–4900 BP) cultures from the West Liao together (sites #1–5, 10–13, 15, 18, 22, 23, 25, 34–38, 87, 88);
- grouping the Zaisanovka culture (5200–3300 BP) in the Southern Primorye (i.e., Zaisanovka 7, Krounovka 1, Novoselishche-4 and Vodopadnoe-7) together with the Yabuli (4500–4000 BP) culture in the adjacent part of the Amur region (sites #41–44 and 250).

In our main analysis with no prior topological constraints, both of these groupings have 0% posterior support in our analysis. We thus find no support in the data for these clades.

###### Individual cultures

The archaeological sites are grouped into “cultures” in the original data. Table 3.2.1 gives the posterior probability that each of these cultures is monophyletic in our analysis, as well as in the analyses of subsets mentioned in Section 3.3.

We find it striking that almost none of these cultures comes out as monophyletic in the posterior. Only the Yayoi and Zaisanovka culture are strongly supported as monophyletic (posterior support 100%

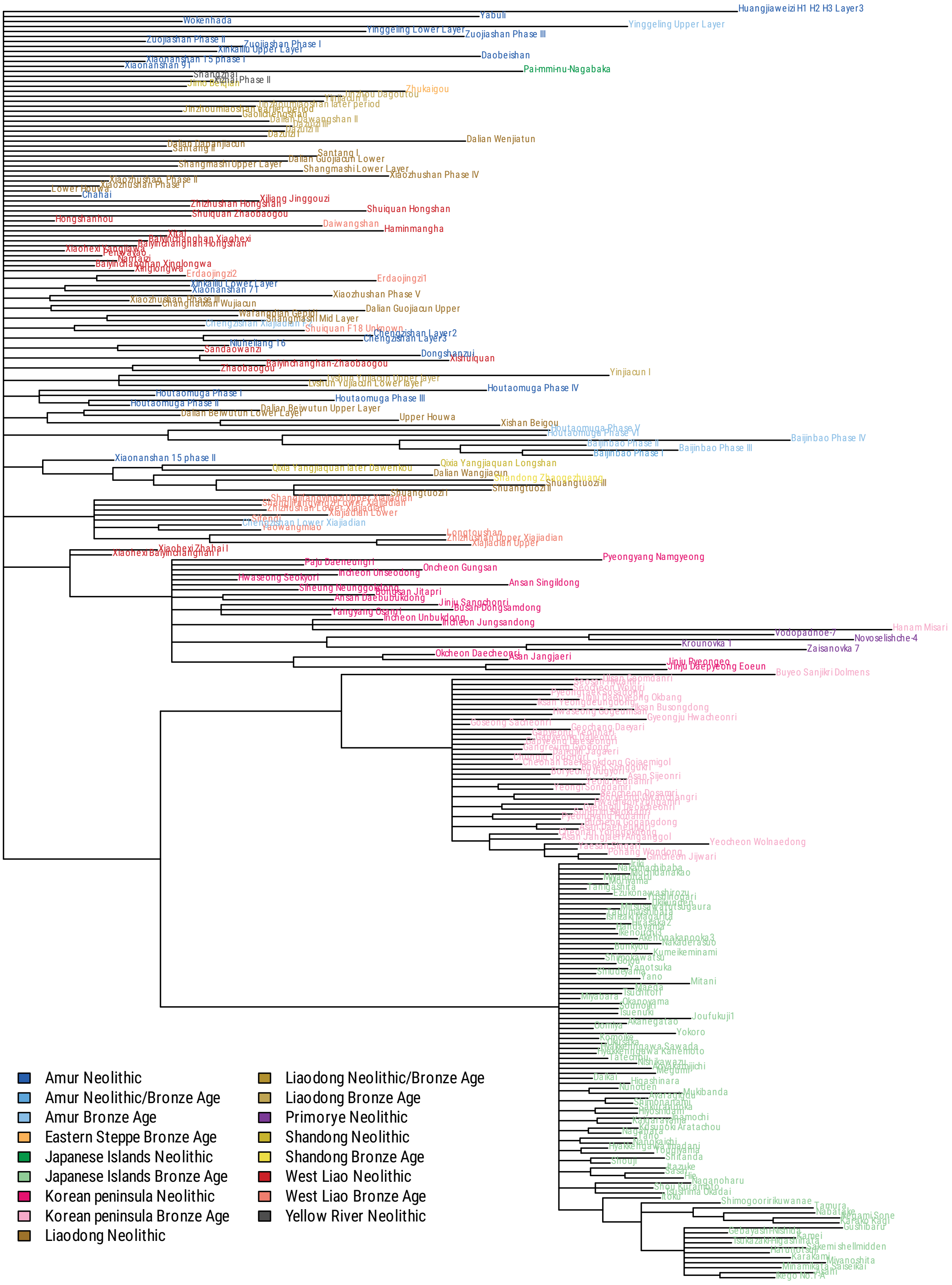

Figure 3.1: Consensus tree for the complete corrected data; only nodes with posterior support above 50% are shown; leaves share a colour when they are in the same region and date to the same period.

and 78.9%, respectively), with also some support for the Middle & Late Yayoi (42.6%) and Zuojiashan (24%) cultures.

For Chulmun and Mumun, the low support is due to the misplacement of the Mumun site Hanam Misari (#117), which occurs in the Chulmun clade. If this site is removed from Mumun and added to Chulmun, we obtain posterior support 40% for Chulmun and 91.1% for Mumun.

| Culture | Number of sites | All data | All data with constraints | Ceramics | Tools | FHBB |
|---|---|---|---|---|---|---|
| Xinglongwa | 6 | 0 | 0.2 | 0.9 | 0 | 0 |
| Hongshan | 11 | 0 | 1.6 | 0 | 0 | 0 |
| Xiaohexi | 4 | 0 | 0 | 0 | 0 | 0 |
| Zhaobaogou | 4 | 1.3 | 4.3 | 5.4 | 0 | 0.2 |
| Haminmangha | 2 | 0 | 0 | 0 | 0 | 0 |
| Lower Xiajiadian | 9 | 0 | 0 | 0 | 0 | 0 |
| Upper Xiajiadian | 5 | 0 | 0 | 0 | 0 | 0 |
| Xiajiadian (all) | 15 | 1.7 | 1.3 | 0 | 0 | 0 |
| Zaisanovka | 4 | 78.9 | 63.8 | 48.4 | 0.8 | 0.1 |
| Lower Xiaozhushan | 2 | 0 | 0 | 0 | 0 | 0 |
| Middle Xiaozhushan | 2 | 6.5 | 6.5 | 0.1 | 0.1 | 26.6 |
| Upper Xiaozhushan | 4 | 0 | 0 | 0 | 0 | 0 |
| Xiaozhushan (all) | 16 | 0 | 0 | 0 | 0 | 0 |
| Pianpuzi | 2 | 11 | 7.4 | 0.7 | 23.9 | 0 |
| Shuangtuozi III | 6 | 0 | 0 | 0 | 0 | 0 |
| Shuangtuozi I | 5 | 0 | 0 | 0 | 0 | 0 |
| Shuangtuozi II | 3 | 0 | 0 | 11.1 | 0 | 0 |
| Shuangtuozi (all) | 15 | 0 | 0 | 0 | 0 | 0 |
| Chulmun | 18 | 1.1 | 19.3 | 0 | 0 | 0 |
| Mumun | 39 | 8.1 | 26.1 | 0 | 0 | 0.6 |
| Yayoi (all) | 84 | 100 | 96.7 | 96 | 0 | 0 |
| Baijinbao | 2 | 0 | 0 | 0 | 0 | 0 |
| Xinkailiu | 4 | 0 | 0 | 0 | 0 | 0 |
| Zuojiashan | 3 | 24 | 28.4 | 0.3 | 0 | 0.1 |

Table 3.1: Posterior probability that each culture is monophyletic, as a percentage.

##### Clades above the culture level and migration hypotheses

In Robbeets et al.'s results, the Xiaohexi sites #7 and #8 occur in a clade with Chulmun sites, whereas Zaisanovka sites are found in a clade with Liaodong sites, supporting according to them two migration events, corresponding to Koreanic and Tungusic, respectively. In our results, however, there is nearly no support for grouping the Xiaohexi sites #7 and #8 with Chulmun, even if the problematic Hanam Misari (#117) Mumun site is added. What we find instead is that a clade comprising Xiaohexi (#7 and #8), Chulmun, Hanam Misari and Zaisanovka (but not Yabuli) has a support of 64.6%, with Zaisanovka sites occurring among Chulmun sites. If these results are interpreted in terms of migration hypotheses, the only possible conclusion is the occurrence of a single migration event over a large zone comprising both Primorye and Korea, and does not support the idea of two distinct migrations.

A second migration hypothesis proposed by Robbeets et al. concerns the grouping of Yayoi and Mumun sites with Bronze age sites from West Liao (e.g. Xiajiadian) or Liaodong (e.g. Houwa). However, in our results, even a Yayoi/Mumun clade is only supported (with 75.5% posterior probabilities) if the Hanam Misari (#117) Mumun site is removed from it. There is no evidence from this dataset for a grouping of Yayoi, Mumun and Liaodong Bronze Age sites, and no support for any migration from West Liao or Liaodong to these areas.

| Culture | Number of sites | All data | All data with constraints | Ceramics | Tools | FHBB |
|---|---|---|---|---|---|---|
| Zhaobaogou+Hongshan+Xinglongwa | 21 | 0 | 100 | 0 | 0 | 0 |
| Zaisanovka+Yabuli | 5 | 0 | 100 | 0 | 0.1 | 0 |
| Yayoi+Mumun | 123 | 6.9 | 18.6 | 0 | 0 | 0 |
| Yayoi+Mumun-117 | 122 | 75.5 | 58.4 | 1.1 | 0 | 0 |
| Xiaohexi(7+8)+Chulmun+117 | 21 | 2.2 | 11.6 | 0.3 | 0 | 0 |
| Xiaohexi(7+8)+Chulmun+117+ Zaisanovka | 25 | 64.6 | 0 | 0 | 0 | 0 |

Table 3.2: Posterior probability that groupings involving sites from several cultures are monophyletic, as a percentage.

#### 3.3 Analysing subsets of the data

The archaeological features used by Robbeets et al. are classified into the following categories:

1. ceramics (69 traits);
2. stone tools (37 traits);
3. buildings/houses (9 traits);
4. food remains (36 traits);
5. shell and bone artefacts (17 traits);
6. burials (13 traits) .

An implicit but essential assumption for Robbeets et al.'s phylogenetic reconstruction to be valid is that there exists a single tree along which all traits evolved. On the other hand, if there exist different trees for the history of (say) ceramics one the one hand, and stone tools on the other hand, then attempting to reconstruct a single tree by merging these traits into a single data set will produce an invalid tree.

We split the data into three subsets: one with the ceramics traits (Fig. 3.2), one with the stone tools traits (Fig. 3.3), and on with all remaining traits (food remains, buildings/houses, shell and bone artefacts, and burials, hereafter denoted by FHBB; Fig. 3.4), to ensure that there are enough traits in each subset. If the implicit assumption is correct, then we expect these analyses to yield a posterior distribution on trees which is similar to that on the complete data, although with larger variance (since we are using less data). If however the tree distributions are significantly different, that is evidence that the different traits do not share a common history.

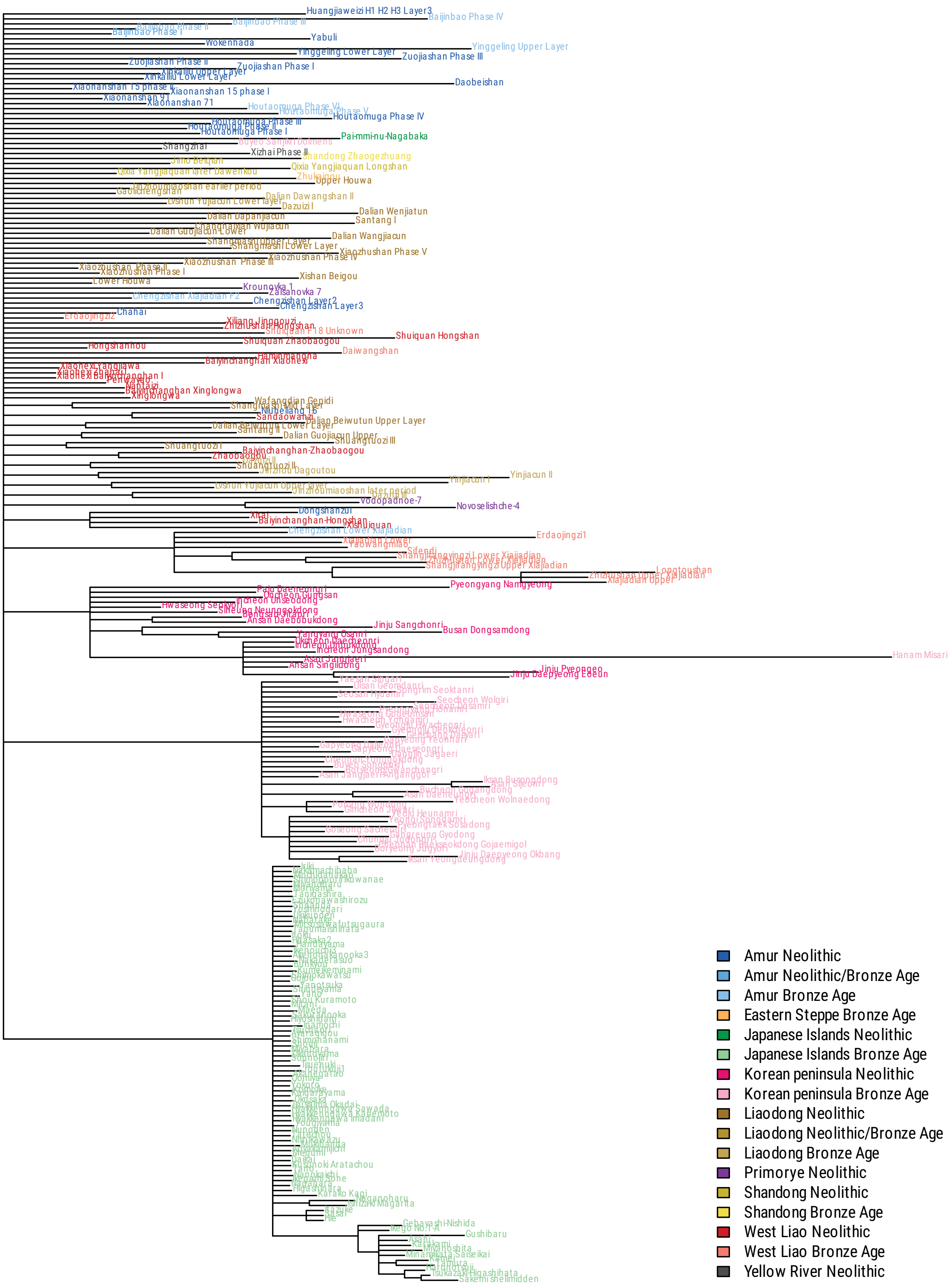

Figure 3.2: Consensus tree for ceramics features only; only nodes with posterior support above 50% are shown; leaves share a colour when they are in the same region and date to the same period.

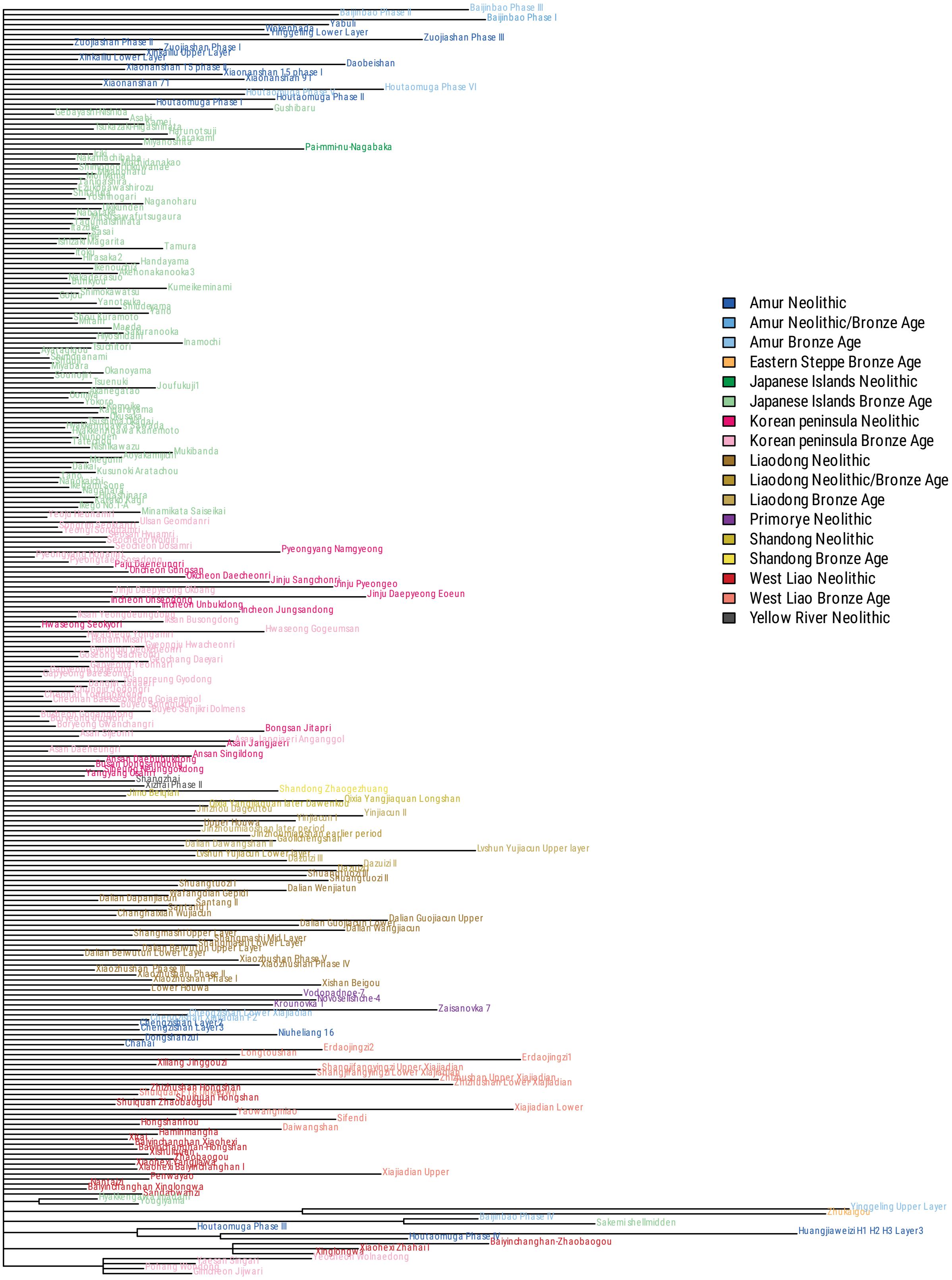

Figure 3.3: Consensus tree for stone tools features only; only nodes with posterior support above 50% are shown; leaves share a colour when they are in the same region and date to the same period.

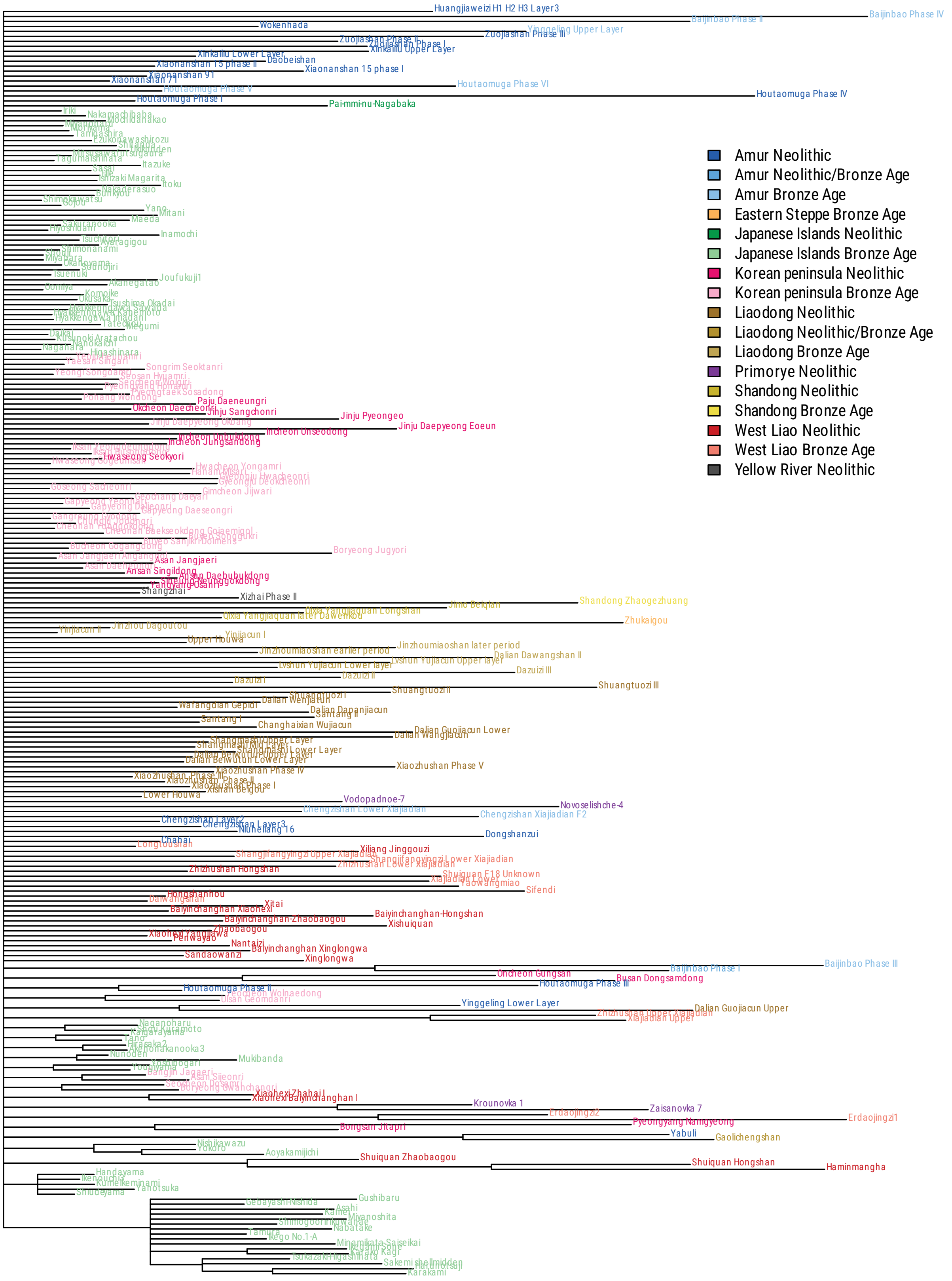

Figure 3.4: Consensus tree for buildings/houses, food remains, sheel and bone artefacts, and burials features only; only nodes with posterior support above 50% are shown; leaves share a colour when they are in the same region and date to the same period.

##### 3.3.1 Root age

The inferred root age is very different between the analyses, as shown in Figure 3.5 and Table 3.3. Only the Ceramics subset agrees with the original analysis.

| Subset | Mean root age | 95% HPD |
|---|---|---|
| All | 7911 | [7134–8867] |
| Ceramics | 8079 | [6914–9492] |
| FHBB | 9649 | [7347–12604] |
| Tools | 11285 | [7700–16025] |

Table 3.3: Posterior root age

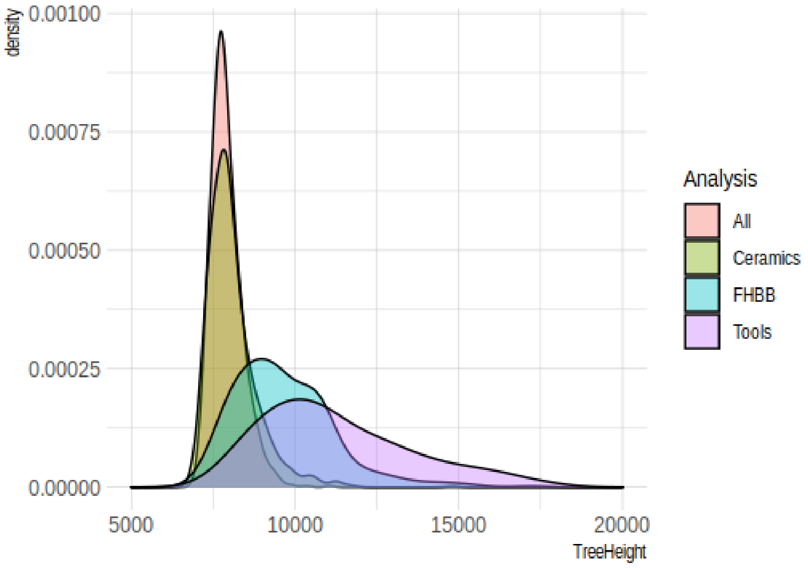

Figure 3.5: Posterior root age

##### 3.3.2 Clade probabilities

We use the BEAST plugin CladeSetComparator to compare distributions of trees (Bouckaert 2020). Each plot below compares two analyses on subsets of the data. In these plots, the tree distributions are close if:

- the points are between the diagonal lines; red points outside of the lines indicate clades with high support in one analysis, low support in the other analysis;
- the histogram at the top left is peaked towards the middle.

These plots all show that there is massive disagreement between the analyses on the various subsets of the data. This indicates that no single tree can capture the history of all these traits.

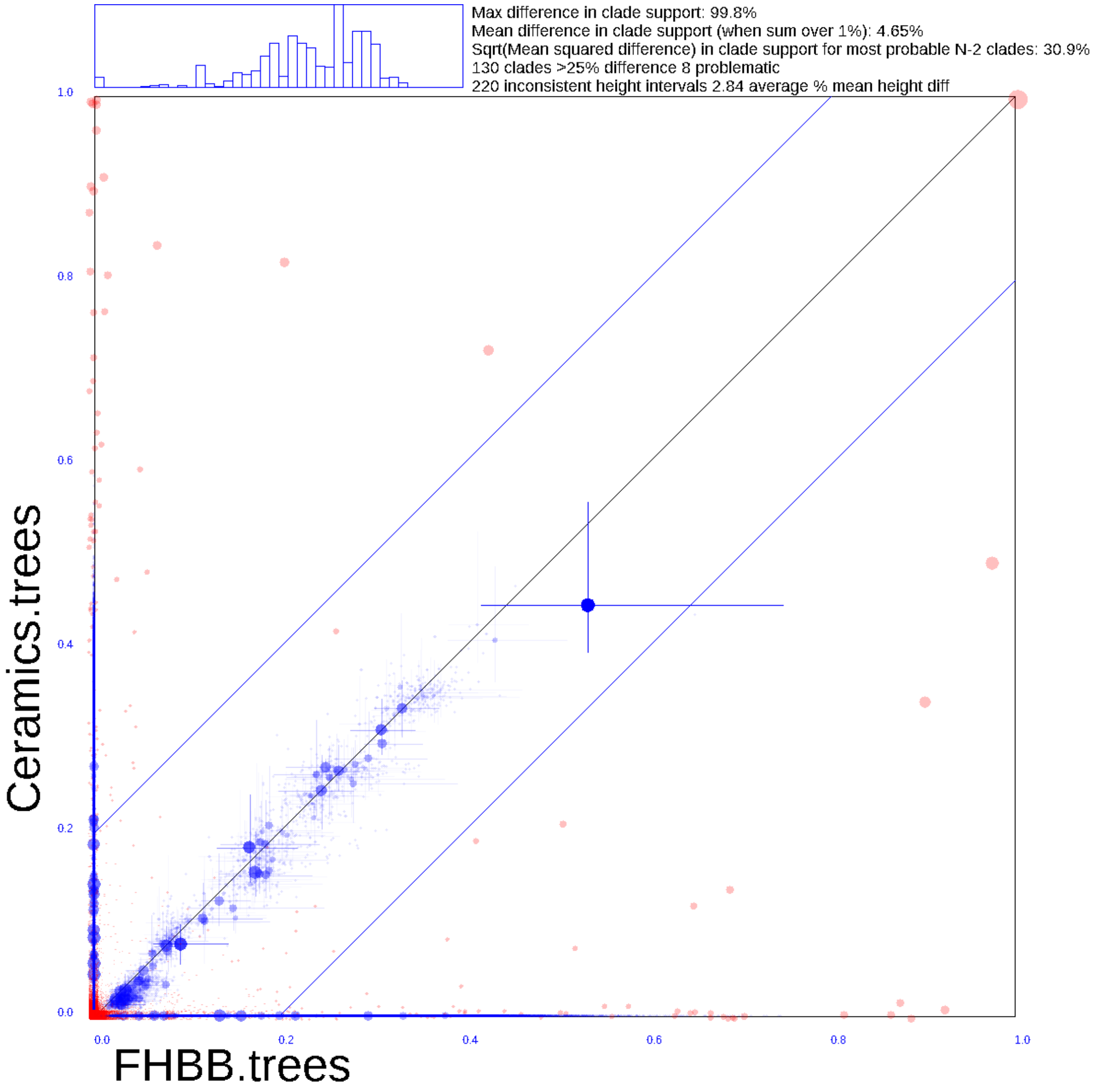

Figure 3.6: Comparison of clades between only FHBB and only Ceramics.

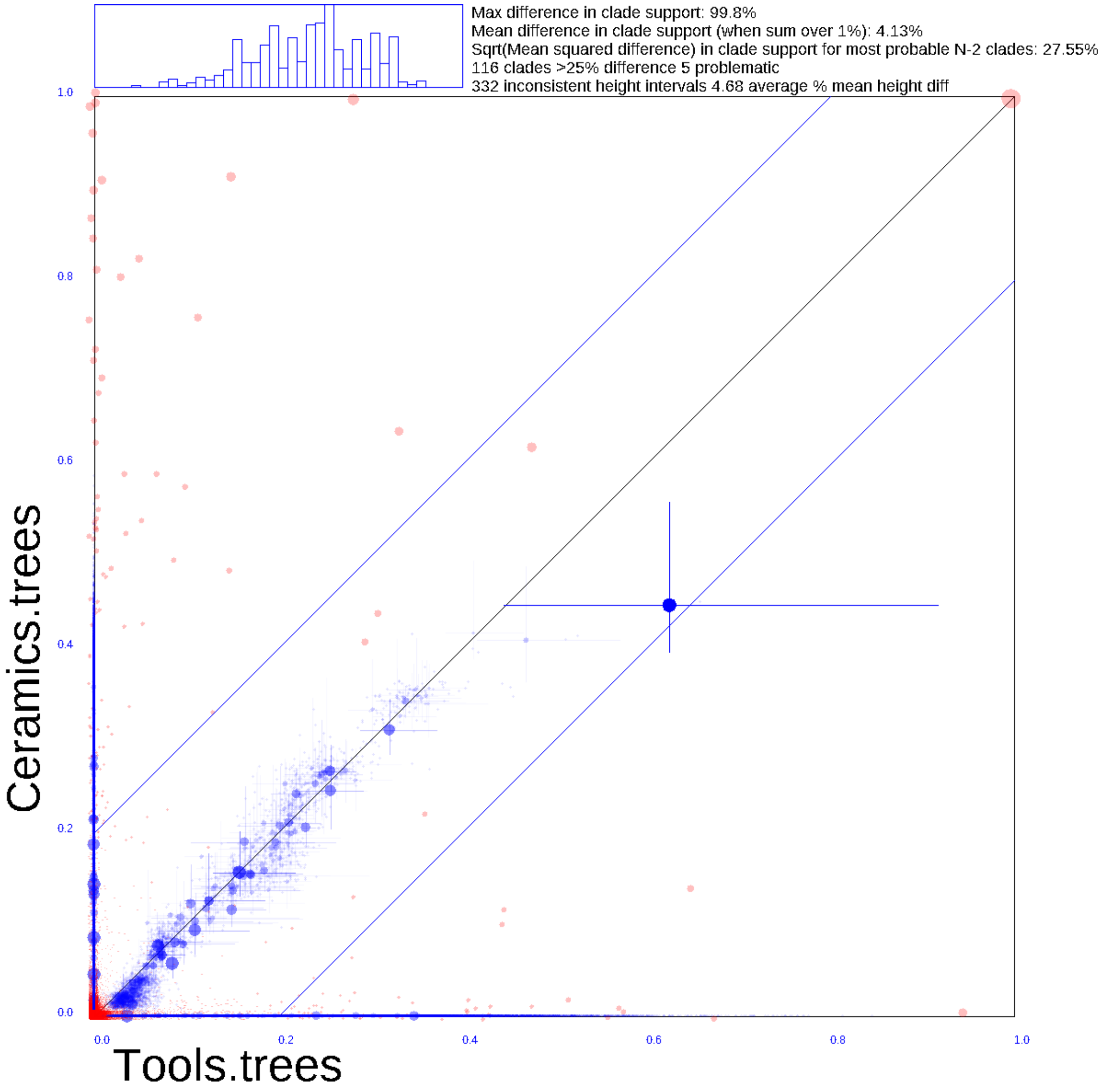

Figure 3.7: Comparison of clades between only Tools and only Ceramics.

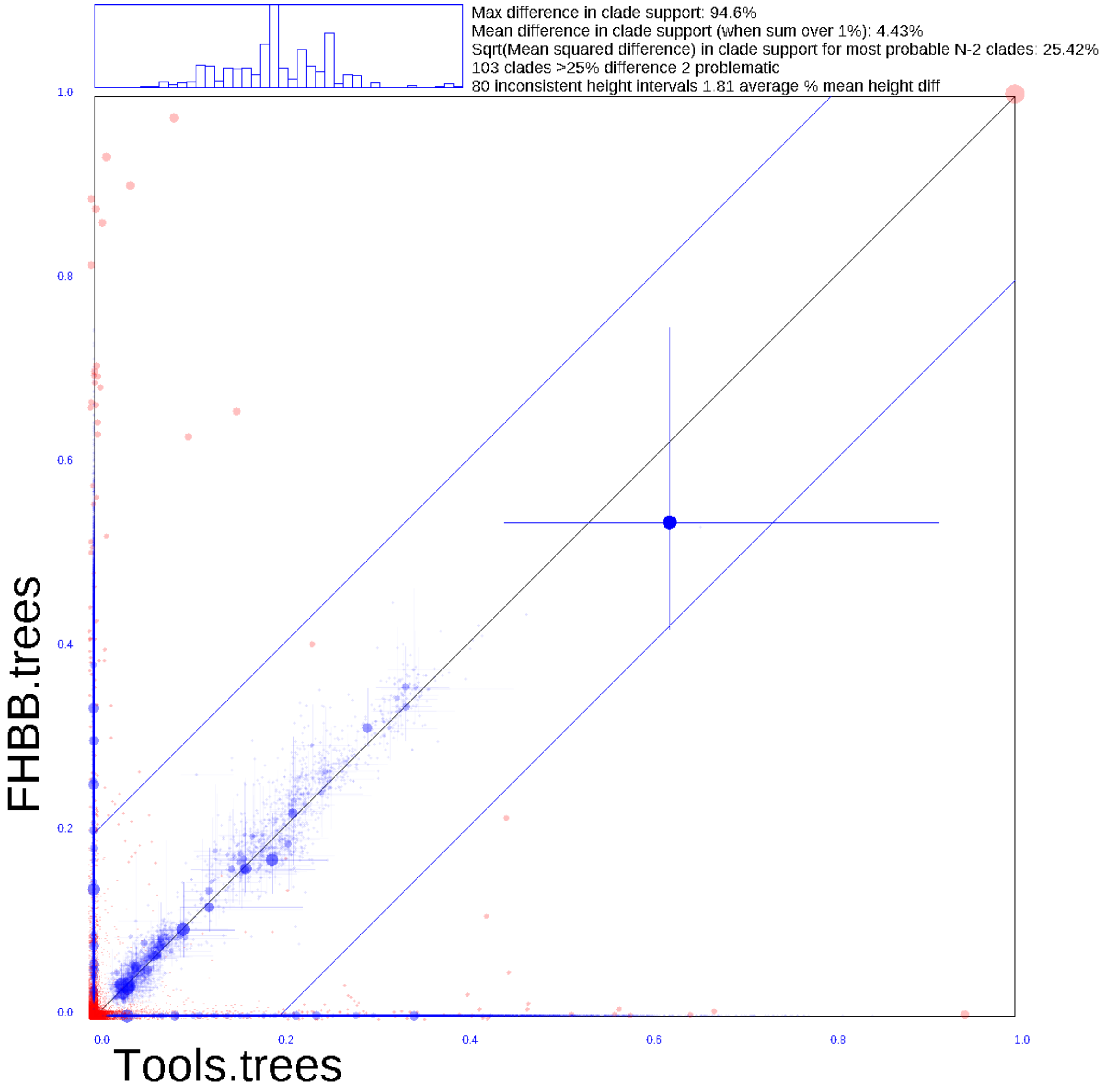

Figure 3.8: Comparison of clades between only Tools and only FHBB.

#### 3.4 Conclusions

Our results make evident that there is little tree structure in the archaeological data selected by Robbeets et al. (2021). Furthermore, the analysis of subsets of the data show that the evolution of these different features followed different histories. We therefore conclude that the data were collected in such a way that using a tree model is inappropriate, and that no faith can be put in the output. In addition, in the few places of the data where there is tree-like signal, the inferred histories does not correspond to the expected output, nor to the conclusions described by Robbeets et al.

### 4 Evaluation of the genetic evidence

#### 4.1 Data mapping

We downloaded the haploid genotyping data used in Robbeets et al.'s (2021) study,[1] then merged it with the previously published data of the Hongshan, the Upper Xiajiadian, and other publicly available modern and ancient reference data from the Allen Ancient DNA Resource[2] using the `mergeit` function from the EIGENSOFT software (Patterson et al. 2006).

#### 4.2 Calculating *f*-statistics

The $f_4$-statistics were calculated in the form of $f_4$ (Mbuti, X; Hongshan, Upper Xiajiadian) using Robbeets et al.'s (2021) newly published populations as X by *qpDstat* v755 implemented in ADMIXTOOLS with the parameters `f4-mode` set to `YES` and `printsd` set to `YES` (Patterson et al. 2012; Peter 2016).

#### 4.3 Conducting the *qpWave* analysis

We used *qpWave* (version 410) implemented in ADMIXTOOLS to test whether the Hongshan and the Upper Xiajiadian population used in Robbeets et al. (2021) were genetically homogenous with respect to their newly published population from Korea, the Ryukyu Islands and Japan. We calculated the *qpWave* statistics using the Hongshan and the Upper Xiajidian populations as left populations and Mbuti, Ando, Changhang, Kuma-Nishioda Yayoi, Nagabaka early, Nagabaka historic, Nagabaka late, Taejungni, Yokchido and Yondaedo as right populations with the parameters `allsnps` set to `YES` and `details` set to `YES` (Patterson et al. 2012).

[1] `https://edmond.mpdl.mpg.de/imeji/collection/59JGAaOpSxRb96Vh`.

[2] AADR, `https://reich.hms.harvard.edu/allen-ancient-dna-resource-aadr-downloadable-genotypes-present-day-and-ancient-dna-data`.

### Abbreviations

Asa. Amami Asama
Aze. Azeri
Bao. Baoan
Bas. Bashkir
BTat. Baraba Tatar
Bur. Buriat
CC Codex Cumanicus
Chu. Chuvash
CTat. Crimean Tatar
Dag. Dagur
Dgx. Dongxiang
Dol. Dolgan
EMJ Early Middle Japanese
Evk. Evenki (Kamnigan)
Evn. Even
Fuk. Fukuoka
Gag. Gagauz
GG Gyeonggi
GW Gangwon
Hac. Hachijo
Hez. Hezhe
HH Hwanghae
Htm. Yaeyama Hatoma
Huz. Huzhu
Ira. Miyako Irabu
Ish. Yaeyama Ishigaki
JJ Jeju
Jpn Japanese
Jur. Jurchen
Kag. Kagoshima
Kalm. Kalmyck
Kar. Karaim
Kaz. Kazakh
KBal. Karachay Balkar
Kgj. Kangjia
Kh. Khalkha
Khk. Khakas
Khl. Khalaj
Kir. Kirghiz
KKal. Kara Kalpak
Kmg. Kamnigan
Kos. Koshiki islands
KTat. Kazan Tatar
KU Kur-Urmi
Kum. Kumamoto
Kum. Kumyk
Man. Manchu
MChu. Middle Chulym
MK (Late) Middle Korean
MMoMuq. Middle Mongolian (Muqaddimat al-adab)
MMoSH Middle Mongolian (Secret History)
Mnh. Minhe
Mog. Moghol
NAlt. North Altai
NanA. Nanai (Middle Amur)
NanB. Nanai (Bikin)
NCC Northern Chungcheong
Neg. Negidal
NETur. Northern Evenki (Tura)
NETut. Northern Evenki (Tutonchany)
NGS Northern Gyeongsang
NHG Northern Hamgyong
NJL Northern Jeolla
Nog. Nogai
NPA Northern Pyongan
Oir. Oirat
OJ Old Japanese
Orc. Oroch
Ork. Orok
Orq. Oroqen
OT Old Turkic
pJ proto-Japonic
pK proto-Koreanic
pM proto-Mongolic
pR proto-Ryukyuan
pTEA proto-Transeurasian
pTg proto-Tungusic

pTk proto-Turkic
Sal. Salar
SAlt. South Altai
SCC Southern Chungcheong
SEChi. Southern Evenki (Chiringda)
SGS Southern Gyeongsang
SHG Southern Hamgyong
Sho. Shor
Shu. Okinawa Shuri
ShYu. Shira-Yughur
SJL Southern Jeolla
Sol. Solon
SPA Southern Pyongan
StEvk. Stony Evenki
Tkm. Turkmen
Tks. Turkish
Tof. Tofa
Tuv. Tuvan
Udi. Udihe
Ulc. Ulcha
Uyg. Uyghur
Uzb. Uzbek
WYug. West Yugur
Xib. Xibe
Yak. Yakut
Yam. Amami Yamatohama
Yng. Yonaguni
Ynm. Okinawa Yonamine
Yor. Amami Yoron

Anthony, David W. 2007. *The horse, the wheel, and language: How Bronze-Age riders from the Eurasian steppes shaped the modern world.* Princeton: Princeton University Press.

Bǎo, Cháolǔ. 1984. *Dōngbù Yùgùyǔ cíhuì.* Huhehaote: Nèi Mēnggǔ Rénmín Chūbǎnshè.

Baskakov, N. A. 1968. *Turkmensko-russkij slovar'.* Moscow: "Sovetskaja èncikolpedija" neširjaty.

Bawden, Charles. 1997. *Mongolian-English dictionary.* London: Kegan Paul International.

Blust, Robert. 1988. The Austronesian homeland: A linguistic perspective. *Asian Perspectives* 26(1). 45–67. http://hdl.handle.net/10125/16918.

Bouckaert, Remco. 2020. *Are these tree sets the same?* https://www.beast2.org/2020/04/20/comparing-tree-sets.html.

Chén, Nǎixióng. 1986. *Bǎoānyǔ cíhuì.* Huhehaote: Nèi Mēnggǔ Rénmín Chūbǎnshè.

Cheremisov, K. M. 1951. *Buryat-Mongol'sko-Russkij slovar'.* Moscow: Gosudarstvennoye izdatel'stvo inostrannykh i nacional'nykh slovarey.

Clauson, Gerard. 1972. *An etymological dictionary of pre-thirteenth-century Turkish.* Oxford: Clarendon Press.

Dunn, Michael & Tiago Tresoldi. 2021. *evotext/ielex-data-and-tree: IELex data and tree (2021/11/08).* Version r20211108. Zenodo. https://doi.org/10.5281/zenodo.5556801.

Enhébātú. 1984. *Dáwòěryǔ cíhuì.* Huhehaote: Nèi Mēnggǔ Rénmín Chūbǎnshè.

Fellner, Hannes A. & Nathan W. Hill. 2019. The differing status of reconstruction in Trans-Himalayan and Indo-European. *Cahiers de Linguistique Asie Orientale* 48(2). 159–172. https://doi.org/10.1163/19606028-04802006.

Forkel, Robert, Johann-Mattis List, Simon J. Greenhill, Christoph Rzymski, Sebastian Bank, Michael Cysouw, Harald Hammarström, Martin Haspelmath, Gereon A. Kaiping & Russell D. Gray. 2018. Cross-Linguistic Data Formats, advancing data sharing and re-use in comparative linguistics. *Scientific Data* 5(180205). 1–10. https://doi.org/10.1038/sdata.2018.205.

Fujimoto, Kenshin. 2011. *Kumamoto-ken hōgen jiten.* Kumamoto: Sōsōsha.

Garnier, Romain, Laurent Sagart & Benoît Sagot. 2017. Milk and the Indo-Europeans. In Martine Robbeets & Alexander Savelyev (eds.), *Language dispersal beyond farming*, 291–311. Amsterdam: John Benjamins. https://doi.org/10.1075/z.215.13gar.

Haenisch, Eric. 1939. *Wörterbuch zu Manghol un Niuca Tobca'an (Yüan-ch'ao pi-shi): Geheime geschichte der mongolen.* Leipzig: Otto Harassowitz.

Hirayama, Teruo (ed.). 1992–1994. *Gendai Nihongo hōgen daijiten.* 9 vols. Tokyo: Meiji Shoin.

Hōsei Daigaku Okinawa Bunka Kenkyūjo (ed.). 1987. *Ryūkyū no Hōgen* 11: *Yaeyama Yonaguni-jima.* http://hdl.handle.net/10114/11991.

Hú, Zēngyì. 1994. *Xīn Mǎn Hàn dà cídiǎn.* Wūlǔmùqí shì: Xīnjiāng Rénmín Chūbǎnshè.

Ikema, Nae. 2003. *Yonagunigo jiten.* Yonaguni-chō: Ikema Nae.

Iwakura, Ichirō. 1941. *Kikai-jima hōgenshū*. Tokyo: Chūō Kōronsha. https://doi.org/10.11501/1852965.

Janhunen, Juha A. 2013. Personal pronouns in Core Altaic. In Martine Robbeets & Hubert Cuyckens (eds.), *Shared grammaticalization: With special focus on the Transeurasian languages*, 211–226. Amsterdam: John Benjamins. https://doi.org/10.1075/slcs.132.16jan.

Jōdaigo jiten henshū iinkai (ed.). 1967. *Jōdaihen* (Jidaibetsu Kokugo Daijiten). Tokyo: Sanseidō.

Kajiku, Shin'ichi. 2020. *Hatoma hōgen jiten*. Tachikawa: National Institute for Japanese Language and Linguistics. https://doi.org/10.15084/00002991.

Kałużyński, Stanisław. 1970. Dagurisches Wörterverzeichnis (part 2). *Rocznik orientalistyczny* 33(2). 109–143.

Khabtagaeva, Bayarma. 2022. On color terms in Dagur. In Bayarma Khabtagaeva (ed.), *Historical linguistics and philology of Central Asia: Essays in Turkic and Mongolic studies*, 473–497. Leiden: Brill. https://doi.org/10.1163/9789004499966_031.

Kiku, Chiyo & Toshizō Takahashi. 2005. *Yoron hōgen jiten*. Tokyo: Musashino Shoin.

Kim, Juwon, Dongho Ko, D. O. Chaoke, Youfeng Han, Lianyu Piao & B.V. Boldyrev. 2008. *Materials of Spoken Manchu*. Seoul: Seoul National University Press.

Kokuritsu Kokugo Kenkyūjo (ed.). 1963. *Okinawago jiten*. Tokyo: Ōkurashō Insatsukyoku. https://doi.org/10.15084/00002266.

Kubo, Tomoyuki, Norikazu Kogura & Shēng Zhuāng. 2011a. *Shibego goishū*. Tōkyō: Tōkyō Gaikokugo Daigaku Ajia Afurika Gengo Bunka Kenkyūjo. https://doi.org/10.15026/95112.

Kubo, Tomoyuki, Norikazu Kogura & Shēng Zhuāng. 2011b. *Shibego no kiso*. Tōkyō: Tōkyō Gaikokugo Daigaku Ajia Afurika Gengo Bunka Kenkyūjo. https://doi.org/10.15026/95113.

Kungnip Kugŏwŏn (ed.). 2016. *Uli mal saem*. https://opendict.korean.go.kr.

Lee, Ki-Moon & S. Robert Ramsey. 2011. *A history of the Korean language*. Cambridge: Cambridge University Press. https://doi.org/10.1017/CBO9780511974045.

Li, Fengxiang & Lindsay Whaley. 2009. Loanwords in Oroqen, a Tungusic language of China. In Martin Haspelmath & Uri Tadmor (eds.), *Loanwords in the world's languages: A comparative handbook*, 525–544. Berlin: De Gruyter Mouton. https://doi.org/10.1515/9783110218442.525.

Lǐ, Kèyù. 1988. *Mongghul Qidar merlong*. Xīníng: Qīnghǎi Rénmín Chūbǎnshè.

Lǐ, Shùlán & Qiān Zhòng. 1986. *Xībó yǔ jiǎnzhì*. Běijīng: Mínzú Chūbǎnshè.

List, Johann-Mattis. 2019. Automatic inference of sound correspondence patterns across multiple languages. *Computational Linguistics* 45(1). 137–161. https://doi.org/10.1162/coli_a_00344.

List, Johann-Mattis, Robert Forkel, Simon J. Greenhill, Christoph Rzymski, Johannes Englisch & Russell D. Gray. 2021. Lexibank: A public repository of standardized wordlists with computed phonological and lexical features. Version 1. *Research Square*. Preprint. https://doi.org/https://doi.org/10.21203/rs.3.rs-870835/v1.

Liu, Li & Xingcan Chen. 2012. *The archaeology of China: From the Late Paleolithic to the Early Bronze Age*. Cambridge: Cambridge University Press. https://doi.org/10.1017/CBO9781139015301.

Mǎ, Guózhōng & Yuánlóng Chén. 2012. *Dōngxiāngyǔ Hànyǔ cídiǎn*. 2nd edn. Lánzhōu: Gānsù Mínzú Chūbǎnshè.

Martin, Samuel E. 1966. Lexical evidence relating Korean to Japanese. *Language* 42(2). 185–251. https://doi.org/10.2307/411687.

Martin, Samuel E. 1970. Shodon: A dialect of the Northern Ryukyus. *Journal of the American Oriental Society* 90(1). 97–139. https://doi.org/10.2307/598434.

Martin, Samuel E. 1987. *The Japanese language through time*. New Haven: Yale University Press.

Martin, Samuel E. 1996. *Consonant lenition and the Macro-Altaic question*. Honolulu: Center for Korean Studies, University of Hawai'i.

Miyagi, Shin'yū. 2003. *Ishigaki hōgen jiten*. 2 vols. Naha: Okinawa Taimusu.

Nadelyayev, Vladimir Mikhaylovich (ed.). 1969. *Drevnetyurkskiy slovar'*. Leningrad: Nauka.

Nakasone, Seizen. 1983. *Okinawa Nakijin hōgen jiten*. Tokyo: Kadokawa Shoten.

Nam, Kwang-U. 1997. *Koŏ sajŏn*. Seoul: Kyohaksa.

Nihon Kokugo Daijiten Dai 2-han Henshū Iinkai, Shōgakkan Kokugo Jiten Henshūbu (ed.). 2000–2002. *Nihon kokugo daijiten*. 2nd edn. 13 vols.

Nugteren, Hans. 2011. *Mongolic phonology and the Qinghai-Gansu languages*. Leiden: Leiden University dissertation. http://hdl.handle.net/1887/18188.

Nugteren, Hans & Marti Roos. 1996. Common vocabulary of the Western and Eastern Yugur languages the Turkic and Mongolic loanwords. *Acta Orientalia Academiae Scientiarum Hungaricae* 49(1–2). 25–91. https://www.jstor.org/stable/43391252.

Osada, Suma & Nahoko Suyama. 1977–1980. *Amami hōgen bunrui jiten*. 2 vols. Tokyo: Kasama Shoin.

Patterson, Nick, Priya Moorjani, Yontao Luo, Swapan Mallick, Nadin Rohland, Yiping Zhan, Teri Genschoreck, Teresa Webster & David Reich. 2012. Ancient admixture in human history. *Genetics* 192(3). 1065–1093. https://doi.org/10.1534/genetics.112.145037.

Patterson, Nick, Alkes L. Price & David Reich. 2006. Population structure and eigenanalysis. *PLoS Genetics* 2(12). e190. https://doi.org/10.1371/journal.pgen.0020190.

Pellard, Thomas. 2022. Ryukyuan and the reconstruction of proto-Japanese-Ryukyuan. In Bjarke Frellesvig, Satoshi Kinsui & John Whitman (eds.), *Handbook of Japanese historical linguistics*. Berlin: De Gruyter Mouton. In press.

Peter, Benjamin M. 2016. Admixture, population structure, and $F$-statistics. *Genetics* 202(4). 1485–1501. https://doi.org/10.1534/genetics.115.183913.

Poppe, N. N. 1931. *Materialy po solonskomu jazyku*. Leningrad: Izdatel'stvo akademii nauk SSSR.

Ramsey, S. Robert. 1991. Proto-Korean and the origin of Korean accent. In William G. Boltz & Michael C. Shapiro (eds.), *Studies in the historical phonology of Asian languages*, 215–238. Amsterdam: John Benjamins. https://doi.org/10.1075/cilt.77.11ram.

Robbeets, Martine. 2020. Basic vocabulary in the Transeurasian languages. In Martine Robbeets & Alexander Savelyev (eds.), *The Oxford guide to the Transeurasian languages*, 645–659. Oxford: Oxford University Press. https://doi.org/10.1093/oso/9780198804628.003.0037.

Robbeets, Martine, Remco Bouckaert, Matthew Conte, Alexander Savelyev, Tao Li, Deog-Im An, Ken-ichi Shinoda, Yinqiu Cui, Takamune Kawashima, Geonyoung Kim, Junzo Uchiyama, Joanna Dolińska, Sofia Oskolskaya, Ken-Yōjiro Yamano, Noriko Seguchi, Hirotaka Tomita, Hiroto Takamiya, Hideaki Kanzawa-Kiriyama, Hiroki Oota, Hajime Ishida, Ryosuke Kimura, Takehiro Sato, Jae-Hyun Kim, Bingcong Deng, Rasmus Bjørn, Seongha Rhee, Kyou-Dong Ahn, Ilya Gruntov, Olga Mazo, John R. Bentley, Ricardo Fernandes, Patrick Roberts, Ilona R. Bausch, Linda Gilaizeau, Minoru Yoneda, Mitsugu Kugai, Raffaela A. Bianco, Fan Zhang, Marie Himmel, Mark J. Hudson & Chao Ning. 2021. Triangulation supports agricultural spread of the Transeurasian languages. *Nature* 599. 616–621. https://doi.org/10.1038/s41586-021-04108-8.

Rozycki, William. 1994. *Mongol elements in Manchu*. Bloomington: Indiana University Press.

Sagart, Laurent, Guillaume Jacques, Yunfan Lai, Robin J. Ryder, Valentin Thouzeau, Simon J. Greenhill & Johann-Mattis List. 2019. Dated language phylogenies shed light on the ancestry of Sino-Tibetan. *Proceedings of the National Academy of Sciences* 116(21). 10317–10322. https://doi.org/10.1073/pnas.1817972116.

Saito, Yoshio. 2008. *The Mongolian words in the Muqaddimat al-Adab: Romanized text and word index (as of January 2008)*. Tech. rep. The Japan society for the Promotion of Science Grant-in-Aid for Scientific Research (B), 2005-2007, research report (2). Tokyo.

Shōgaku Tosho (ed.). 1989. *Nihon hōgen daijiten*. 3 vols. Tokyo: Shōgakkan.

Starostin, Sergei, Anna Dybo & Oleg Mudrak. 2003. *Etymological dictionary of the Altaic languages*. 3 vols. Leiden: Brill.

Toadeva, Buljash Xojchievna (ed.). 2009. *Kalmycko-russkij slovar'*. Elista: Gerel.

Todaeva, B. X. 1986. *Dagurskij jazyk*. Moscow: Nauka.

Tomihama, Sadayoshi. 2013. *Miyako Irabu hōgen jiten*. Naha: Okinawa Taimusu.

VanBik, Kenneth. 2009. *Proto Kuki-Chin: A reconstructed ancestor of the Kuki-Chin languages*. Berkeley: University of California, Berkeley. https://stedt.berkeley.edu/pubs_and_prods/STEDT_Monograph8_Proto-Kuki-Chin.pdf.

Vovin, Alexander. 2006. Why Manchu and Jurchen look so non-Tungusic? In Alessandra Pozzi, Juha Antero Janhunen & Michael Weiers (eds.), *Tumen jalafun jecen akū: Manchu studies in honour of Giovanni Stary*, 255–266. Wiesbaden: Harrassowitz.

Vovin, Alexander. 2010. *Koreo-Japonica: A re-evaluation of a common genetic origin*. Honolulu: University of Hawai'i Press. http://www.jstor.org/stable/j.ctt6wqz03.

Vovin, Alexander. 2018. Two Tungusic etymologies. *Studia Orientalia Slovaca* 17(1). 125–134. https://fphil.uniba.sk/fileadmin/fif/katedry_pracoviska/kvas/SOS_abstracts_17.1/SOS_17_1__2018_.pdf.

Wang, Wei, Christopher Bae & Xin Xu. 2020. Chinese prehistoric eyed bone needles: A review and assessment. *Journal of World Prehistory* 33(3). 385–423. https://doi.org/10.1007/s10963-020-09144-2.

Weiers, Michael. 1963. *Die Sprache der Moghol der Provinz Herat in Afghanistan*. Wiesbaden: VS Verlag für Sozialwissenschaften. https://doi.org/10.1007/978-3-663-01721-9.

Wu, Mei-Shin, Nathanael E. Schweikhard, Timotheus A. Bodt, Nathan W. Hill & Johann-Mattis List. 2020. Computer-assisted language comparison: State of the art. *Journal of Open Humanities Data* 6(2). 1–14. https://doi.org/10.5334/johd.12.

Yamamoto, Kengo. 1969. *Manshūgo kōgo kiso goishū*. Tōkyō: Tōkyō Gaikokugo Daigaku Ajia Afurika Gengo Bunka Kenkyūjo.

Yonaguni Hōgen Jiten Henshū Iinkai (ed.). 2019. *Dunan munui jiten*. Yonaguni-chō: Yonaguni-chō Yakuba.

Yudakhin, Konstantin Kuz'mich. 1965. *Kirgizsko-russkiy slovar': Okolo 40 000 slov*. Moscow: Sovetskaya entsiklopediya.

Zhang, Menghan, Shi Yan, Wuyun Pan & Li Jin. 2019. Phylogenetic evidence for Sino-Tibetan origin in northern China in the Late Neolithic. *Nature* 569. 112–115. https://doi.org/10.1038/s41586-019-1153-z.

Zikmundová, Veronika. 2013. *Spoken Sibe: Morphology of the inflected parts of speech*. Prague: Karolinum Press.
